# Development of non-sedating antischistosomal benzodiazepines

**DOI:** 10.1101/2024.01.26.577323

**Authors:** Md Yeunus Mian, Dishary Sharmin, Prithu Mondal, Jawad Bin Belayet, M Mahmun Hossain, Paul McCusker, Kaetlyn T. Ryan, Alexander Y Fedorov, Heather A Green, Spencer S. Ericksen, Mostafa Zamanian, V. V. N. Phani Babu Tiruveedhula, James M. Cook, John D. Chan

**Author notes:** Address correspondence to John D. Chan.

## Abstract

The neglected tropical disease schistosomiasis infects over 200 million people worldwide and is treated with just one broad spectrum antiparasitic drug (praziquantel). Alternative drugs are needed in the event of emerging praziquantel resistance or treatment failure. One promising lead that has shown efficacy in animal models and a human clinical trial is the benzodiazepine meclonazepam, discovered by Roche in the 1970’s. Meclonazepam was not brought to market because of dose-limiting sedative side effects. However, the human target of meclonazepam that causes sedation (GABA_A_Rs) are not orthologous to the parasite targets that cause worm death. Therefore, we were interested in whether the structure of meclonazepam could be modified to produce antiparasitic benzodiazepines that do not cause host sedation. We synthesized 18 meclonazepam derivatives with modifications at different positions on the benzodiazepine ring system and tested them for *in vitro* antiparasitic activity. This identified five compounds that progressed to *in vivo* screening in a murine model, two of which cured parasite infections with comparable potency to meclonazepam. When these two compounds were administered to mice that were run on the rotarod test, both were less sedating than meclonazepam. These findings demonstrate the proof of concept that meclonazepam analogs can be designed with an improved therapeutic index, and point to the C3 position of the benzodiazepine ring system as a logical site for further structure-activity exploration to further optimize this chemical series.

## Introduction

While the neglected tropical disease schistosomiasis infects over 200 million people worldwide (1), control of the disease is almost entirely reliant on one drug, praziquantel (PZQ). A recent metaanalysis of PZQ efficacy over four decades indicates cure rates between 57-88% (2), but there is the concern that resistance may emerge with repeated rounds of mass drug administration (3). Parasite resistance can be selected in the laboratory (4–6), and there are numerous reports of treatment failure in the field (7, 8). An alternative antischistosomal drug would be of obvious use in the event of either PZQ treatment failure or resistance. However, older antischistosomal chemotherapies tend to either have unacceptable toxicity (ex. antimony compounds) or lack broad efficacy against the two species of schistosomes infecting humans in Africa (e.g. oxamniquine is only effective against *Schistosoma manoni* (9) and metrifonate is only effective against *Schistosoma haematobium* (10)). Benzodiazepines are promising antischistosomal candidates because they are effective against both of these African species(11, 12) Binding and functional assays indicate that MCLZ and PZQ do not appear to act on the same target (13, 14). Additionally, unlike PZQ, MCLZ can cure schistosomiasis at the juvenile, liver stage of parasite infections (11, 15).

The antischistosomal activity of meclonazepam (MCLZ, also referred to as Ro 11-3128 or 3-methyl clonazepam) was discovered by Roche in the late 1970’s in a screen of over 400 benzodiazepines (11). The drug was advanced to a small human trial and found to cure infections with *S. mansoni* and *S. haematobium* (12). However, patients also experienced pronounced sedative side effects at doses required for antischistosomal activity. Attempts were made to block this sedation - the GABA_A_R antagonist flumazenil was developed and was shown to have no impact on the antiparasitic effect of MCLZ (16). However, flumazenil only transiently antagonized MCLZ sedation when the two drugs were given together (17) due to pharmacokinetic differences between the two drugs (flumazenil has a half life of less than one hour (18), while MCLZ’s half life can be up to 40 hours (19)).

While MCLZ and other benzodiazepines cause sedation due to positive allosteric modulator activity at human GABA_A_R chloride channels, these compounds exert antiparasitic effects through different schistosome targets. The sequenced genomes of parasitic flatworms lack GABA_A_Rs (20–22), indicating that it is unlikely the parasite targets of benzodiazepines such as MCLZ are homologous to the human GABA_A_Rs receptors that cause sedation (23). The initial studies on MCLZ’s effects on schistosomes in the 1970’s showed that the drug causes rapid Ca^2+^ influx, muscle contraction and tegument damage (24). MCLZ was recently shown to act as an agonist at a Ca^2+^ permeable schistosome TRP (transient receptor potential) channel (14), identifying the likely parasite receptor for this drug. This finding that MCLZ acts on a TRP channel is consistent with the original Roche screening data that the vast majority of benzodiazepines which are known potent agonists or PAMs of mammalian GABA_A_Rs do not have antischistosomal activity (11, 13, 21). Because the host and parasite targets are different, we reasoned it should be feasible to develop MCLZ derivatives that retain antischistosomal activity but exhibit reduced sedation. Past work from our labs supported this hypothesis. We found benzodiazepines known to lack ɑ1GABA_A_R affinity exhibit *in vitro* antischistosomal activity (21), which led us to expand our screen to further explore the structure-activity relationship of these compounds on parasites.

Here, we have screened 18 new compounds with a focus on molecules with modifications near the C3 position of MCLZ, which could be tolerated without loss of antiparasitic activity. Two compounds were confirmed to exhibit *in vivo* efficacy against *S. mansoni* while displaying a lack of sedation, or greatly reduced sedation, relative to the parent molecule MCLZ.

## Results

### *In vitro* screen of meclonazepam derivatives on adult schistosomes

In order to identify antiparasitic benzodiazepines with decreased sedative potential, we synthesized MCLZ derivatives with modifications at various positions and first determined whether they retained antischistosomal activity. We synthesized 17 derivatives with modifications to the N1, C3, N4, C7 and C2’ positions of the benzodiazepine ring system (**Figure 1A, Supplemental Table 1**) and screened these compounds at a high concentration (30 µM for 14 hours) against adult *S. mansoni in vitro* to triage inactive compounds. Active compounds caused worm contractile paralysis, while worms incubated with inactive compounds remained motile with no visible change in worm movement or morphology (**Figure 1A**). Many positions on MCLZ were not amenable to substitution. Analogs with modifications at the N1 (MYM-II-53), N4 (MYM-V-48, MYM-V-49) and C7 (MYM-V-03, MYM-IV-01, MYM-IV-02, MYM-IV-03, MYM-IV-12, MYM-IV-63, MYM-II-72) lacked activity in this initial assay. However, modification of the C3 position was more variable. While some compounds were inactive (MYM-II-53, MYM-II-74, MYM-V-50), other modifications did retain activity (MYM-V-19, MYM-II-83, MYM-V-56, MYM-III-10, MYM-V-58).

**Figure 1.**
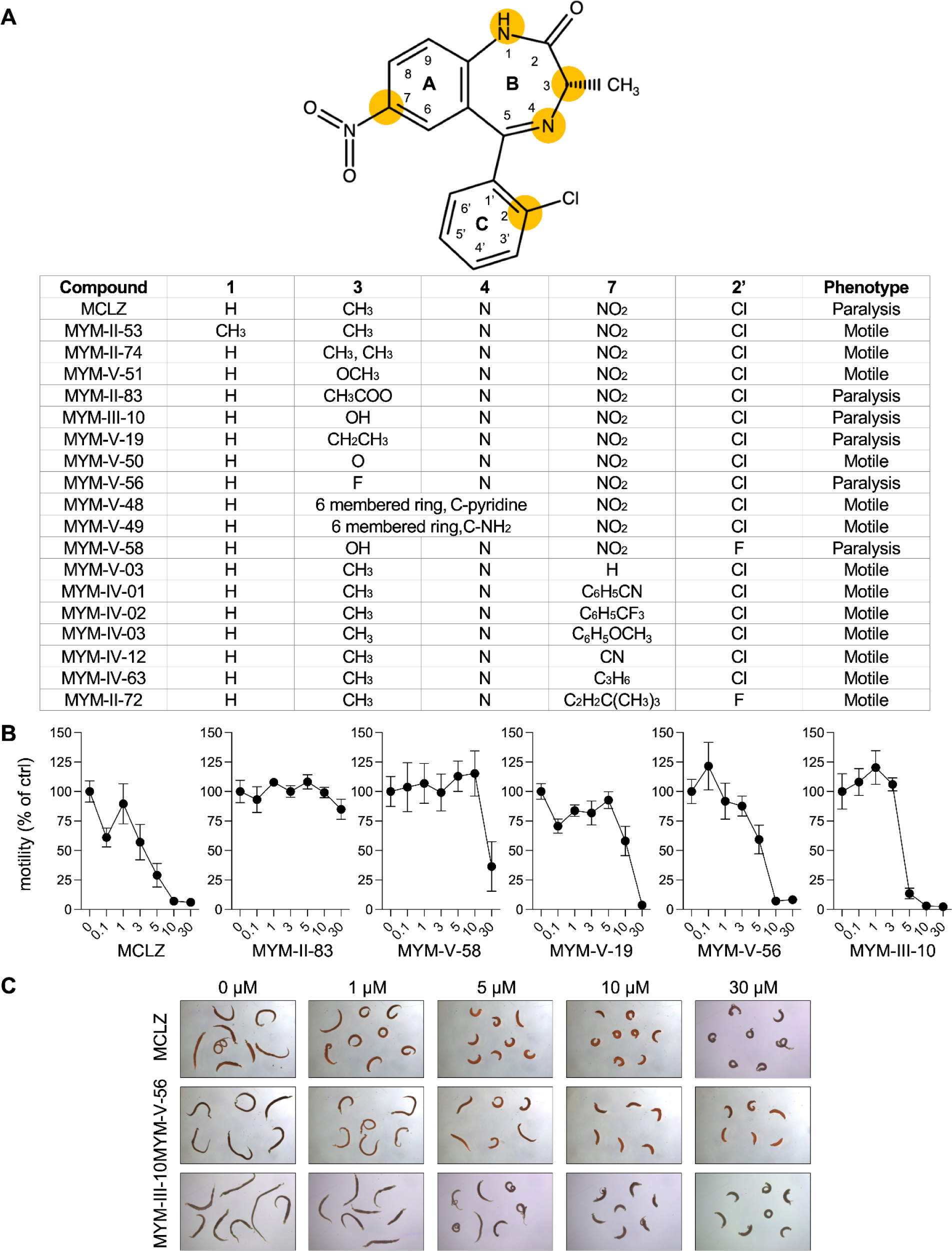
*In vitro* effects of MCLZ analogs on adult *S. mansoni*. **(A)** Top - structure of MCLZ with the benzodiazepine ring system numbered and the modified positions highlighted in yellow. Bottom - MCLZ analogs synthesized with modifications to various positions (1, 3, 4, 7 and 2’) on the benzodiazepine ring system. Adult S. mansoni (5-10 worm pairs) were incubated with compound *in vitro* (30 µM for 14 hours) and the resulting phenotype is noted (paralysis indicating active compounds and motile indicating compounds with no noticeable effects on movement / morphology). **(B)** Concentration-response curves showing potency of compounds shown to be active in (A). Motility is normalized to movement of DMSO control treated worms (values reflect mean ± standard error of n=8 worms). **(C)** Comparison of the morphology of worms treated with MCLZ and the two most potent compounds, MYM-V-56 and MYM-III-10.

Compounds that exhibited antischistosomal activity in this initial screen were then assayed at a series of decreasing concentrations to explore their structure-activity relationship. Adult *S. mansoni* were incubated in compound (0.1 - 30 µM for 14 hours) and motility was measured using a high content imaging system and the wrmXpress pipeline to quantify worm movement (25, 26). Most of these compounds were less active than MCLZ, but two compounds (MYM-V-56 and MYM-III-10) with substitutions at the C3 position retained antischistosomal activity at potency near the parent molecule MCLZ (**Figure 1B&C**). The effects of these compounds on motility (**Figure 1B**) and morphology (**Figure 1C**) were observed to be slightly less potent than MCLZ. However, MYM-V-56 and MYM-III-10 were initially screened as racemic mixtures, while MCLZ is the purified (S) enantiomer of 3-methyl clonazepam and the (R) enantiomer has been reported to be inactive on parasites (27).

### *In vivo* efficacy of benzodiazepine leads in a murine model of schistosomiasis

In order to determine whether hit compounds identified as having *in vitro* activity at 30 µM displayed *in vivo* antischistosomal activity, they were administered to mice infected with *S. mansoni* and the hepatic shift assay was used to observe acute antischistosomal activity, similar to the original Roche study on MCLZ (11). Adult worms normally live within the mesenteric vasculature, but upon exposure to anthelmintic compounds paralyzed worms are swept through the portal vein to the liver (28). Mice were administered compound by oral gavage at a dose of 100 mg / kg and then euthanized three hours later to observe the portion of worms in the liver versus the mesenteries (**Figure 2A**). Only 1.8 ± 1.8% of worms were found in the livers of mice treated with vehicle control. Following MCLZ treatment, 100% of worms were found in the liver and no worms were found in the mesenteries of any mice. Following treatment with the two compounds that showed greatest *in vitro* efficacy, MYM-V-56 and MYM-III-10, 100% of worms were located in the liver and no worms were found in the mesenteries. Generally, compounds that showed weaker potency *in vitro* were less active in the hepatic shift assay. MYM-V-19 was only active at the highest concentration (30 µM) *in vitro* and caused little hepatic shift (5.0 ± 3.2% of worms were found in the liver). MYM-V-58 was similarly only weakly active *in vitro*, but caused a moderate hepatic shift (60.0 ± 23.4% of worms were found in the liver). MYM-II-83, which contains an ester at the C3 position that is likely hydrolyzed *in vivo*, was not active *in vitro* but displayed moderate activity *in vivo* (81.0 ± 10.6% hepatic shift).

**Figure 2.**
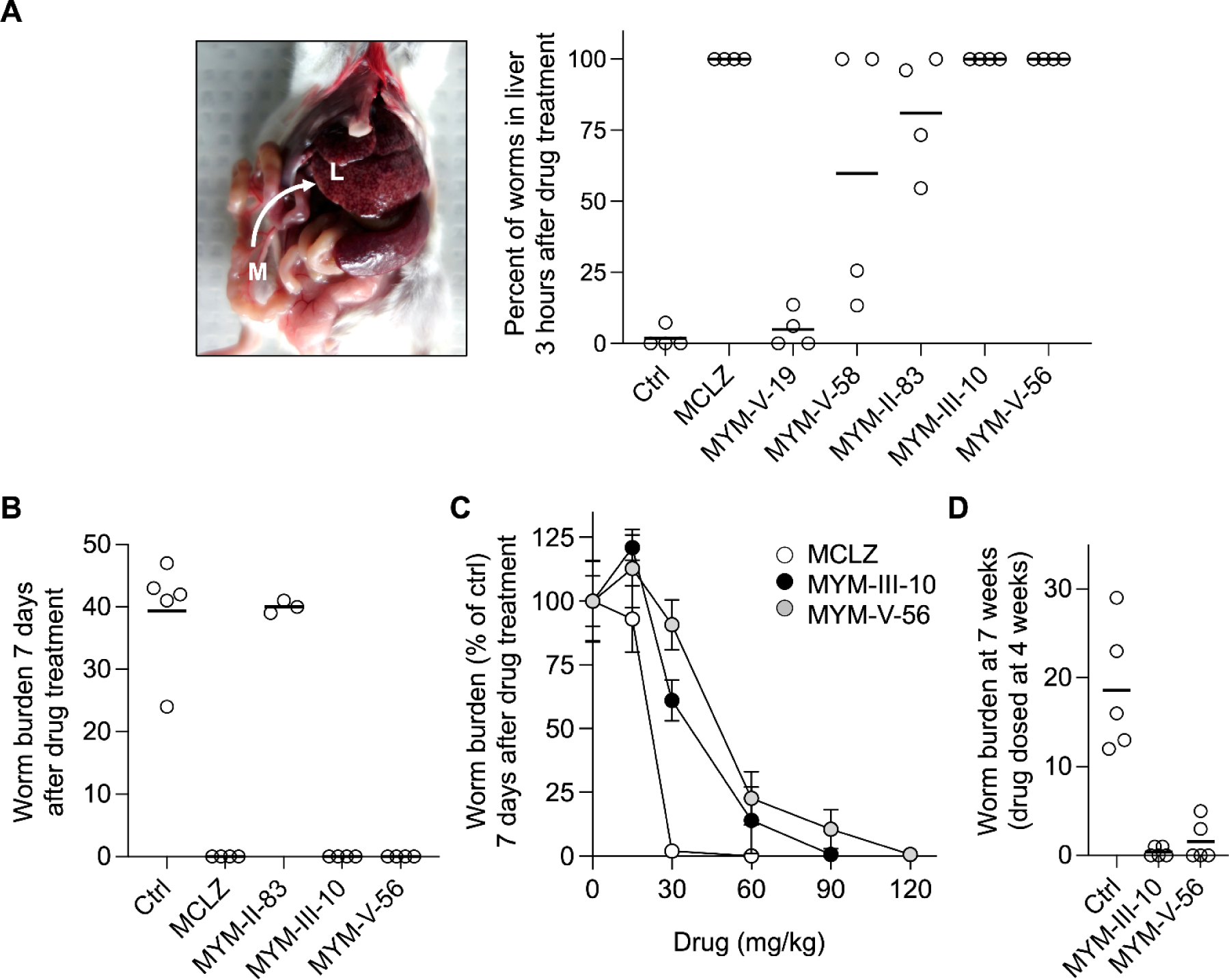
*In vivo* effect of MCLZ analogs on *S. mansoni*. **(A)** Acute *in vivo* antischistosomal activity of MCLZ derivatives, assayed by measuring the percentage of worms in the liver (L) as opposed to the mesenteric vasculature (M) following drug treatment (oral dose of 100 mg / kg). Open symbols reflect percentages for individual mice, solid bars indicate the mean percentage for the treatment cohort. **(B)** Curative activity of MCLZ derivatives following a single oral dose of 100 mg / kg. Mice were euthanized one week after drug administration and worm burden was measured. **(C)** Potency of MCLZ (white symbols), MYM-III-10 (black symbols) and MYM-V-56 (gray symbols) at curing infections of adult *S. mansoni* (mean ± standard error). **(D)** Curative activity of MYM-III-10 and MYM-V-56 against juvenile, liver stage parasites. (open symbols reflect the worm burden of individual mice, solid bars indicate the mean percentage for the treatment cohort).

The three most efficacious compounds, MYM-III-10, MYM-V-56 and MYM-II-83, were then administered to mice (single oral dose of 100 mg / kg) and euthanized after one week to count the number of surviving worms. Mice treated with vehicle control harbored 39.4 ± 4.0 worm pairs, but no live worms were found in MCLZ, MYM-III-10 or MYM-V-56 treated mice (**Figure 2B**). Mice treated with MYM-II-83 did not have any reduction in worm burden compared to control (40.0 ± 0.6 worm pairs per mouse), in contrast to the moderate acute activity of this compound in the hepatic shift assay.

Given that MYM-V-56 and MYM-III-10 exhibited acute *in vivo* efficacy and cured infections at 100 mg / kg, we assessed the potency of these compounds relative to MCLZ by treating infected mice with a range of doses (10-120 mg / kg) and counting the live worm burden after one week (**Figure 2C**). MCLZ cured infected mice at a dose of 30 mg / kg, in line with the effective dose between 33-57 mg / kg reported in the initial Roche study (11). MYM-V-56 and MYM-III-10 displayed curative activity at slightly higher doses of 90-120 mg / kg. This is consistent with the *in vitro* results where the racemate of these compounds were slightly less potent than MCLZ, which is the purified active (S) isoform (**Figure 1B&C**).

Finally, one of the unique features of MCLZ compared to other antischistosomal drugs such as PZQ or oxamniquine is that MCLZ is active against juvenile liver stages of worms (11, 29). Therefore, we wanted to confirm that MYM-V-56 and MYM-III-10 were also capable of clearing these stages of parasites. Infected mice were dosed with both compounds (100 mg / kg) at 4 weeks post infection, which corresponds to the period where PZQ displays the least efficacy. Mice were then euthanized at 7 weeks post infection to count the surviving worms. A single dose of either MYM-V-56 or MYM-III-10 reduced parasite burden by over 90% (**Figure 2D**).

We conducted these initial experiments on MYM-III-10 and MYM-V-56 with racemic preparations of compound, given the resources involved in achieving enantiomeric separation of the (R) and (S) isomers at sufficient scale for *in vivo* mouse studies. However, we reasoned that the purified (S) enantiomers would be more potent, as is the case with (S) versus (R)-MCLZ (27). The two enantiomers of MYM-III-10 and MYM-V-56 were separated by preparative HPLC and adult worms were incubated in each compound *in vitro* to observe paralysis. For MYM-V-56, both the (S) and (R) enantiomer caused contractile paralysis of worms above 5 µM after 14 hours incubation (**Figure 3A**). While these two compounds initially appear equipotent from these data, the kinetics of drug action were quite different. Worms were incubated in 10 µM of each enantiomer and tracked over time. (S)-MYM-V-56 caused contractile paralysis within minutes, while the (R) enantiomer did not cause this phenotype until after several hours incubation in drug (**Figure 3B**). It is likely that the (S) enantiomer is more active *in vivo*, because of the rapid kinetics of the hepatic shift observed with racemic MYM-V-56 treatment (**Figure 2A**). This was confirmed by dosing mice harboring adult parasites orally with both enantiomers. Mice were euthanized one week after drug administration to assess worm burden and *in vivo* antischistosomal activity of each compound. (S)-MYM-V-56 caused a reduction in worm burden of more than 90% at 30 mg / kg (2 ± 3 worm pairs per mouse, compared to 27 ± 3 worm pairs per mouse with vehicle control), while (R)-MYM-V-56 did not decrease worm burden up to the highest dose tested (90 mg / kg) (**Figure 3C**).

**Figure 3.**
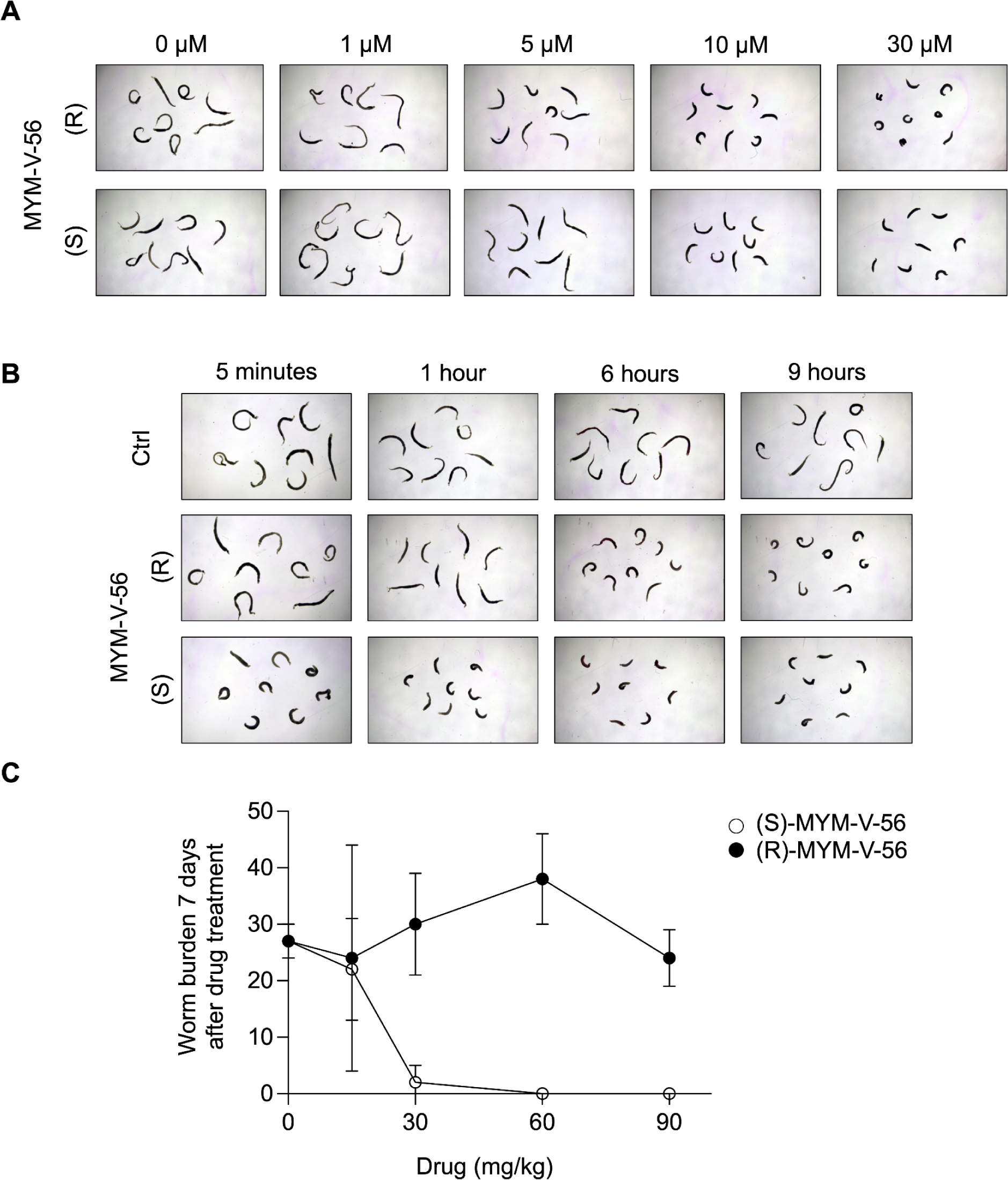
Antischistosomal activity of MYM-V-56 enantiomers. **(A)** *In vitro* effect of purified (R) and (S)-MYM-V-56 enantiomers on adult worms (14 hour incubation). **(B)** Time course of the onset of contractile paralysis caused by each enantiomer (10 µM incubation for each). **(C)** *In vivo* efficacy of each MYM-V-56 enantiomer against adult parasites. Mice were dosed with various amounts of each compound six weeks post infection and euthanized one week later to assess worm burden.

Worms incubated with various concentrations of both (S) and (R)-MYM-III-10 for 14 hours also displayed contractile paralysis, with the (S) enantiomer displaying greater potency than the (R) enantiomer (**Figure 4A**). Unlike MYM-V-56, both enantiomers displayed *in vivo* activity, with (S)-MYM-III-10 curing infections at 60 mg / kg while (R)-MYM-III-10 cured infections at 90 mg / kg (**Figure 4B**). However, given the hydroxyl group on the C3 position of this compound, we anticipated that this compound may racimerize over time in aqueous solution. This has been shown for other benzodiazepines with a C3 hydroxy substituent (e.g. oxazepam, lorazepam, and temazepam)(30). Indeed, purified MYM-III-10 enantiomers racemized within one hour (**Supplementary File 1**). Therefore, (S)-MYM-III-10 may still be more active than the (R) isoform, if much of the antischistosomal activity seen with the (R) enantiomer is in fact due to conversion to the (S) form.

**Figure 4.**
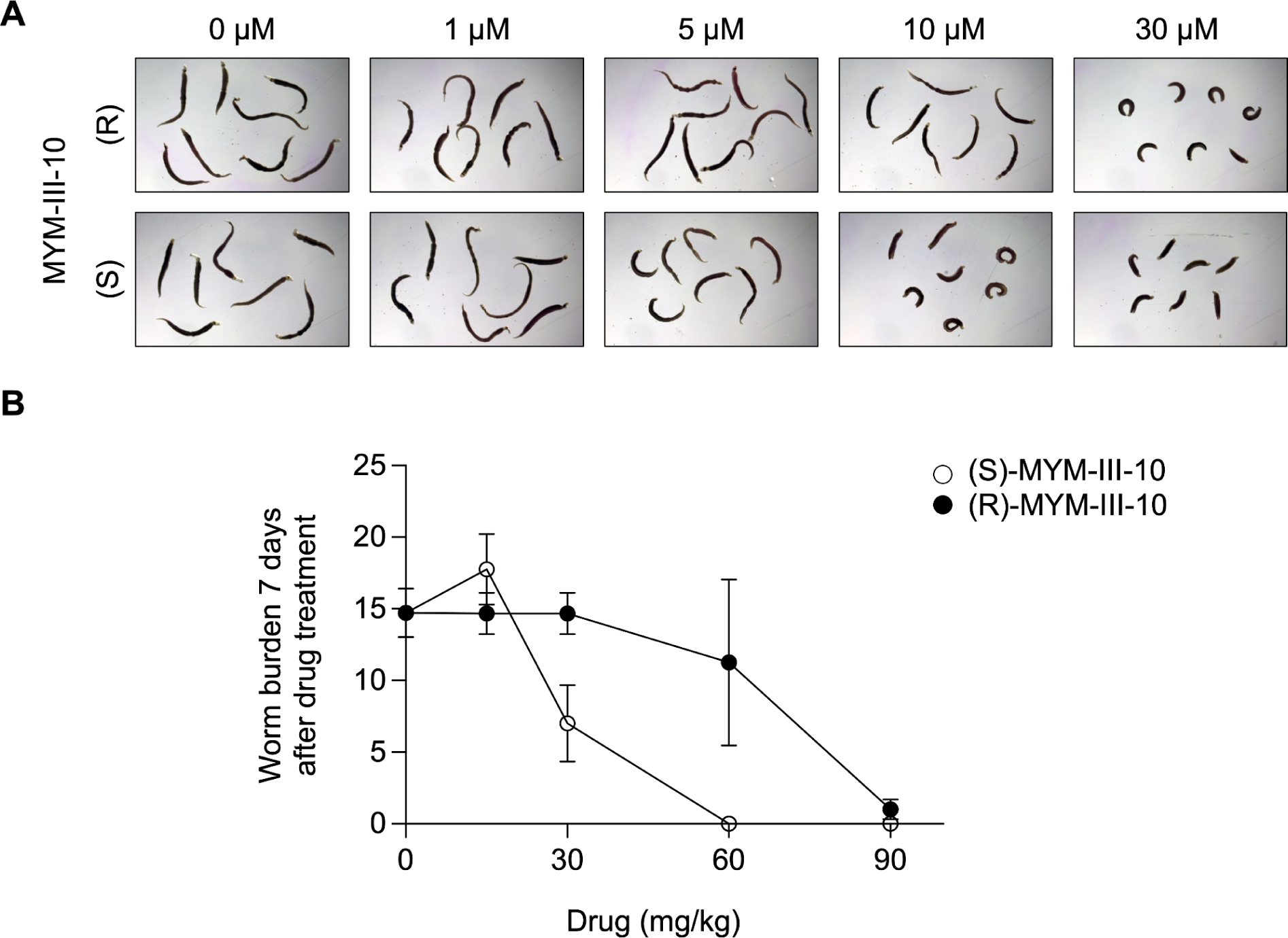
Antischistosomal activity of MYM-III-10 enantiomers. **(A)** *In vitro* effect of purified (R) and (S)-MYM-III-10 enantiomers on adult worms (14 hour incubation). **(B)** *In vivo* efficacy of each MYM-III-10 enantiomer against adult parasites. Mice were dosed with various amounts of each compound six weeks post infection and euthanized one week later to assess worm burden.

### *In vivo* sedative activity of benzodiazepine lead compounds

Given that both MYM-V-56 and MYM-III-10 both retain the antischistosomal activity of MCLZ, we assessed the sedative activity of these compounds. Mice were trained to perform the rotarod test, running on the rod for 300 seconds at a speed of 20 RPM. Animals not infected with schistosomiasis were chosen for this assay, because the hepatomegaly caused by parasite infection can interfere with mouse mobility. Animals dosed orally with increasing amounts of MCLZ or test compound and after 5 minutes were placed on the rotarod device to perform the test. Whenever a mouse fell off of the device, this time point was noted and the test was ended for that animal. For all other animals, the test was ended at 300 seconds. Mice dosed with MCLZ displayed impaired performance on the rotarod test, consistent with this assay serving as a measurement of sedation. At the dose of 30 mg / kg (which corresponds to the dose that cures infection) only 2 / 14 mice were able to complete the test, and the average time they were able to run on the device was 102 ± 26 seconds (**Figure 5**). Mice treated with MYM-III-10 (30 mg / kg) exhibited less impairment, running on the device for an average of 246 ± 26 seconds, and mice treated with MYM-V-56 (30 mg / kg) ran on the device for an average of 232 ± 30 seconds. The purified (R) and (S)-MYM-V-56 enantiomers were also tested and displayed minimal sedation, running for 293 ± 8 seconds and 299 ± 1 seconds, respectively, at the 30 mg / kg dose.

**Figure 5.**
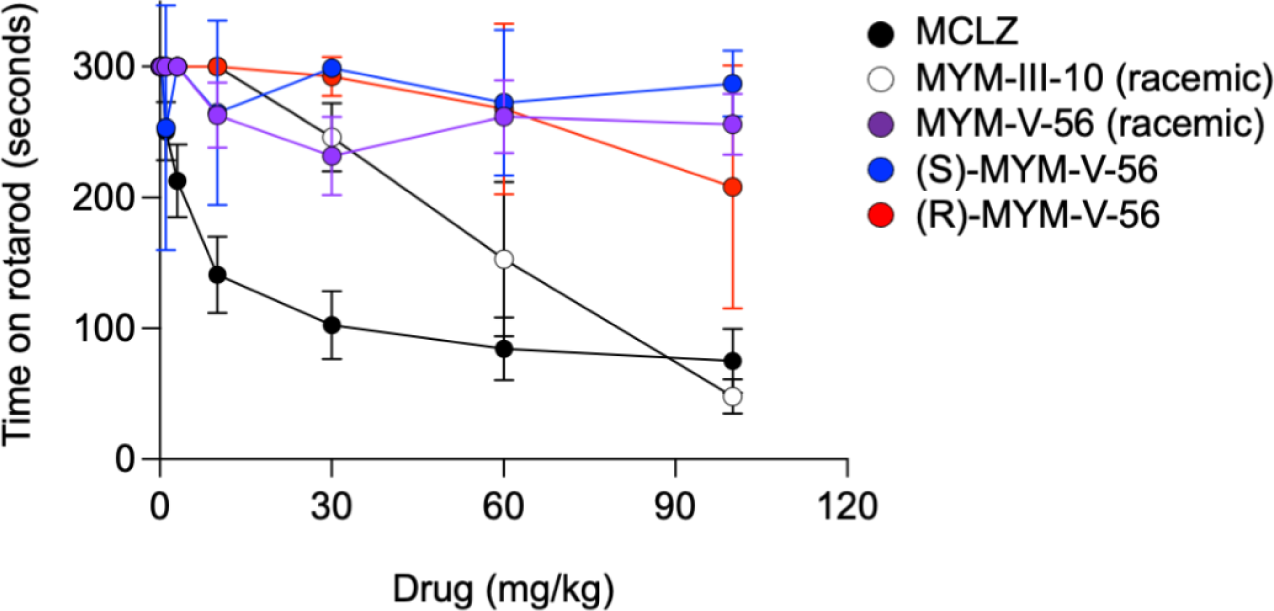
Sedating activity of antischistosomal benzodiazepines. Sedation was measured by ability to complete the rotarod test. Mice were dosed with varying amounts of test compound or vehicle control and assayed for their ability to run on the device for 3 minutes (symbols denote mean ± standard error, black = MCLZ, white = MYM-III-10, purple = MYM-V-56, blue = (S)-MYM-V-56 and red = (R)-MYM-V-56.

While it may appear from these results that MYM-V-56 does not impair mouse locomotion / coordination, observing the mice in their cages following treatment indicates that this compound does have psychoactive effects, even if it is not sedating. Videos of mice treated with vehicle control, MCLZ and MYM-V-56 (100 mg / kg each) are shown in **Supplementary Files 2 - 4**. MYM-V-56 treated mice appear unusually hyperactive. In fact, in order to record a 20 second video of these mice it was necessary to place a plexiglass cover over the cage to keep them from jumping out and escaping. We also observed mice treated with MYM-V-56 that did not complete the rotarod test did not appear sedated. Instead, they would jump off of the device and attempt to escape during the assay. Therefore, while this compound does not have pronounced sedating activity, it does appear to have other psychoactive effects.

In order to investigate the action of the two hit compounds, MYM-III-10 and MYM-V-56, on host targets, we performed a series of binding and functional assays. Binding assays revealed that both compounds displace diazepam radioligand from rat brain membrane preparations with similar potency as MCLZ. Therefore, both MYM-III-10 and MYM-V-56 appear to have affinity for host GABA_A_Rs (**Figure 6A**), even if they are less sedating *in vivo*. Similarly, both compounds have similar activity in functional assays measuring positive allosteric modulator activity at GABA_A_Rs, stimulating peak currents measured from ɑ1β2γ2 GABA_A_ channels recombinantly expressed in mammalian CHO-K1 cells (**Figure 6B**).

**Figure 6.**
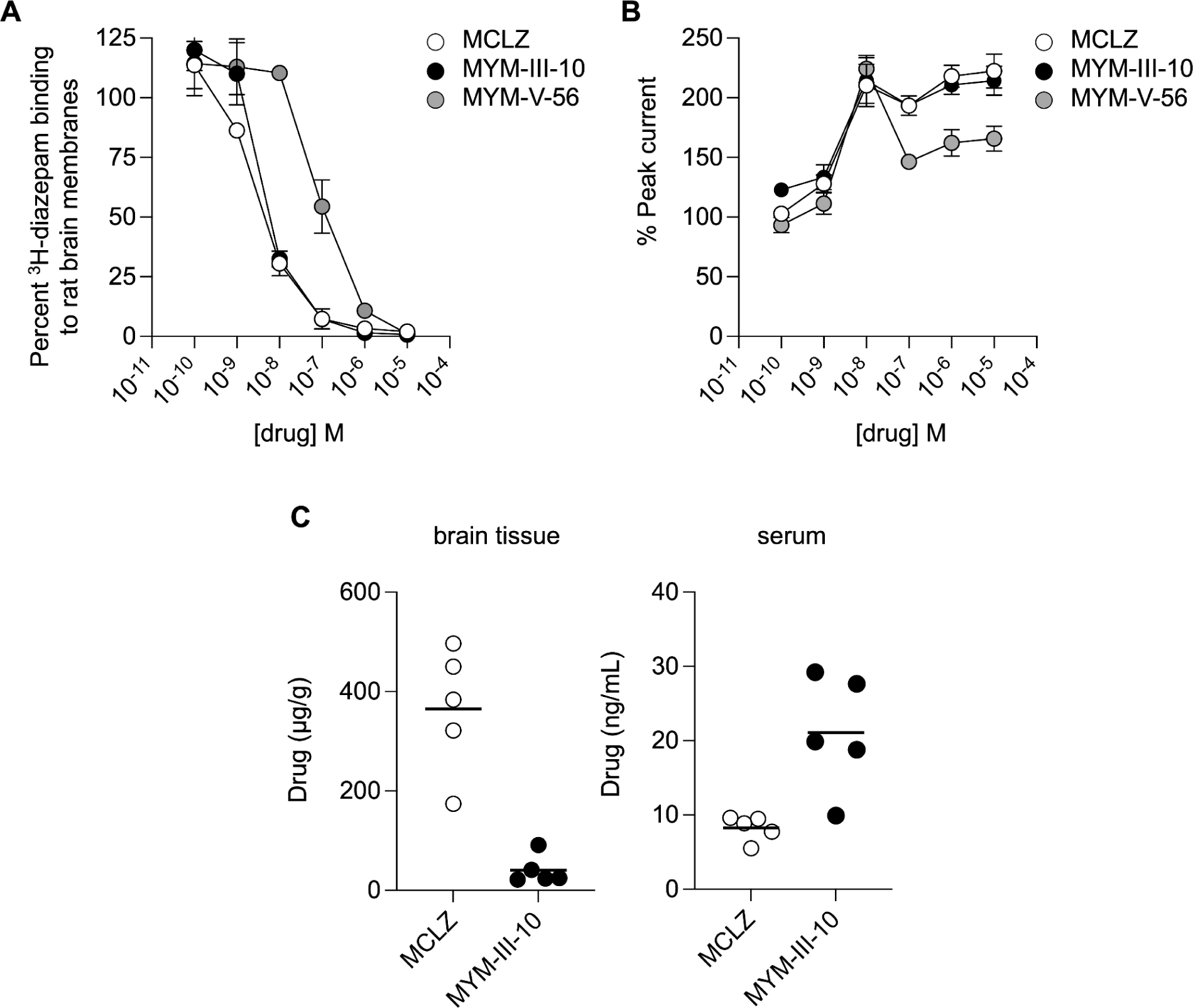
Activity of antischistosomal benzodiazepines on mammalian CNS targets. **(A)** Affinity for CNS GABA_A_Rs was measured by assaying the ability of MCLZ (open symbols), MYM-III-10 (black symbols) or MYM-V-56 (gray symbols) to displace ^3^H-diazepam radioligand from rat brain membranes. **(B)** Positive allosteric modulator (PAM) activity of MCLZ, MYM-III-10 and MYM-V-56 was measured by recording GABA evoked Cl^-^ currents from ɑ1β2γ2 GABA_A_ channels recombinantly expressed in mammalian CHO-K1 cells. All three compounds displayed PAM activity. **(C)** Levels of MYM-III-10 and MCLZ in either mouse brain tissue or serum were measured by HPLC following oral dosing (100 mg / kg).

Given the increased polarity of MYM-III-10 (ClogP 1.01) relative to MCLZ (ClogP 2.39), we hypothesized that the decrease in sedation observed with MYM-III-10 treatment may be due to less compound crossing the blood brain barrier. This was confirmed by measuring levels of drug in serum and brain tissue. Mice were treated orally with either MCLZ or MYM-III-10 as in the antischistosomal and rotarod assays, then euthanized to harvest whole blood and dissect brains from each animal. Drug levels were measured in each sample, revealing that indeed MCLZ but not MYM-III-10 accumulated in the brain tissue. This was not due to a lack of MYM-III-10 bioavailability, since this compound was found at high levels in the serum (**Figure 6C**).

An explanation for the lack of MYM-V-56 sedation is less straightforward. Like MCLZ, it is a relatively nonpolar compound (ClogP 2.42) that would be expected to cross the blood brain barrier. We reasoned that activity at receptors other than GABA_A_Rs may mediate the psychoactive effects of this compound, and submitted MYM-V-56 to a panel of 44 different binding and functional assays (Eurofins Safety Screen44 Panel, **Supplementary Table 2**) to profile activity against a range of targets at a concentration of 10 µM. These results confirmed that MYM-V-56 binds GABA_A_Rs, but no other assay showed significant displacement of radioligand. MYM-V-56 was also submitted to the NIH NIMH Psychoactive Drug Screening Program (PDSP)(31), where binding assays were performed against 46 different targets. These data also did not show significant displacement of radioligand at any of the targets screened (**Supplementary Table 2)**. Therefore, future work will be required to elaborate on the mammalian targets of MYM-V-56 and minimize the psychoactive effects of this compound.

## Discussion

It has been known for over 40 years that certain benzodiazepines exhibit antischistosomal activity (27), including the compound MCLZ which was curative in a human clinical trial (12). While several studies have investigated additional MCLZ derivatives (21, 32, 33), until now it has not been shown that it is possible to develop a benzodiazepine that retains antiparasitic effects *in vivo* but lacks sedative effects.

The two compounds that resulted from this screen, MYM-V-56 and MYM-III-10, both contain modifications at the C3 position of MCLZ. This position was particularly interesting for exploration given the comparison of MCLZ and the anxiolytic / anti-seizure medication clonazepam. Clonazepam is identical to MCLZ, except it lacks a methyl group at the C3 position. This difference makes clonazepam approximately three times *less* potent on schistosomes (causing paralysis at 10 µM, compared to 3 µM for MCLZ), but it binds host GABA_A_Rs with approximately three times *higher* potency (displacement of diazepam radioligand with a K_i_ = 0.82 nM, compared to 2.4 nM for MCLZ) (21, 33). Other pairings with a functional group on the benzodiazepine C3 position can also slightly impair GABA_A_R binding affinity, such as nordazepam (K_i_ = 8.9 nM) versus oxazepam which has a C3 hydroxy group (K_i_ = 38 nM)(34). While substitution of the MCLZ methyl group with a hydroxyl group (MYM-III-10) or fluorine (MYM-V-56) was tolerated, larger substitutions lost or attenuated antiparasitic activity (for example, putting two methyl groups at the C3 position (MYM-II-74), increasing the length of the alkyl chain to an ethyl group (MYM-V-19), or adding an ester (MYM-II-83)). Putting a ketone at this position (MYM-V-50) also resulted in a loss of activity, which is somewhat expected since stereochemistry is known to impact antischistosomal activity (27) and this modification results in a loss of the chiral stereocenter at the C3 position. Future work elaborating on the structure-activity relationship of compounds with modifications at this position may further optimize the antischistosomal activity of this series.

The recent finding that MCLZ acts as an agonist on a schistosome TRP channel enables modeling of the chemical series screened in this study within the benzodiazepine binding pocket of the putative parasite target and a comparison of the SAR at human receptors (**Figure 7, Supplemental Files 5-8**). Interestingly, even though the worm TRP channel and human GABA_A_Rs belong to distinct classes of ion channels with little structural similarity, several common positions on the benzodiazepine ring system are required for activity at both targets. It has been well established that the chirality of the C3 position and presence of electron withdrawing groups at the C7 and phenyl C2’ positions was required for benzodiazepine binding affinity to GABA_A_Rs (34–38), and these interactions were confirmed by resolving the structure of diazepam and alprazolam-bound α1β3γ2L GABA_A_Rs (39). As expected, meclonazepam docked into this binding pocket adopts a similar pose interacting with side chains known to mediate binding affinity (**Figure 7**). However, our SAR data shows similar criteria exists at these same positions of the benzodiazepine molecule for anti-parasitic effects. The importance of the nitro functional group at the C7 position is clear from our chemical series; removing the nitro so that the C7 is unmodified causes a loss of activity (MYM-V-03), as does replacing the nitro with another functional group (MYM-IV-01, MYM-IV-02, MYM-IV-03, MYM-IV-12, MYM-IV-63). This is likely due to the electrostatic interactions between the nitro group and tyrosine and histidine side chains in the binding pocket (**Figure 7**, (14)). Similarly, the C2’ position was known to be critical due to the inactivity of nitrazepam (21) and modeled interaction with Y864 supported by mutagenesis experiments (14). The importance of the chlorine at this position is demonstrated by the reduced activity of MYM-V-58, which is identical to the active MYM-V-56 but contains a fluorine at the C2’ position. Finally, the compound MYM-II-53, which contains a methyl group on the N1 position, was inactive in our screen and was also inactive against the recombinantly expressed TRP channel Smp_333650 (14). This compound was also inactive on schistosomes in reference (33), along with numerous other N1 alkylated compounds. The patent filing by Roche (US4031078A, (27)) for anthelmintic benzodiazepine derivatives stated the N1 could contain a lower alkyl group, but this is not supported by our data or those in references (14, 33). This is one distinction compared to the criteria for GABA_A_R affinity, where N1 alkyl groups are tolerated (e.g. N1 methyl on diazepam).

**Figure 7.**
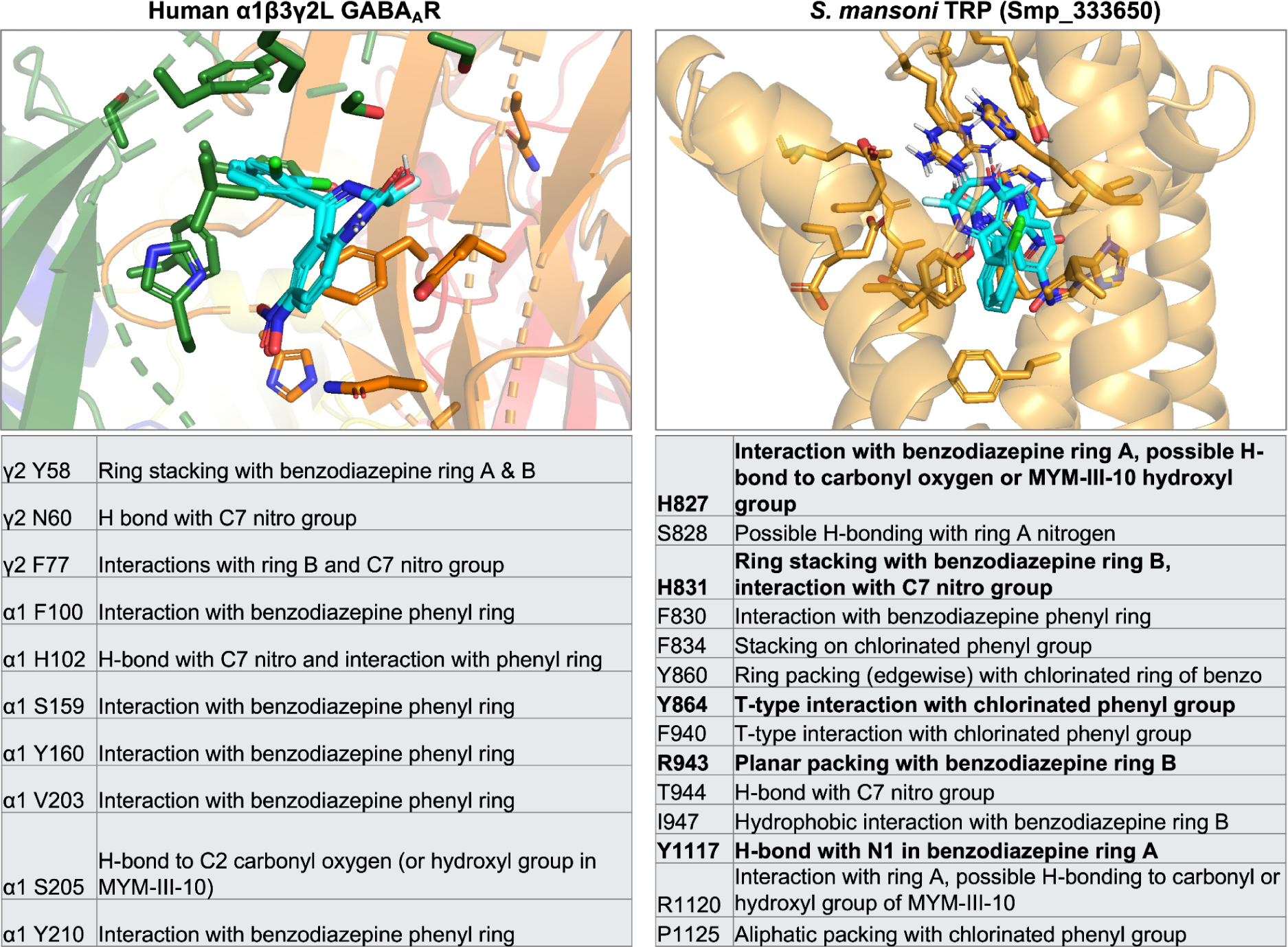
Modeled benzodiazepine interactions with host and parasite receptors. Left - Meclonazepam, (S)-MYM-V-56 and (S)-MYM-III-10 (cyan) docked into the benzodiazepine binding pocket of the human α1β3γ2L GABA_A_ receptor (cyro-EM structure 6HUP, reference(39). γ2 subunit is shown in green, α1 subunit in orange. Right - the same three benzodiazepines docked into the VSLD cavity of the *S. mansoni* TRP Smp_333650 (alphafold structure prediction for A0A5K4FCC0, reference(14)). Bottom - predicted potential interactions between ligands and amino acid side chains. Bold = amino acids experimentally validated by mutagenesis studies in reference (14). Panels are taken from the docked ligands provided in supplementary files 5 and 6 (.pdb files), docking scores are provided in supplementary file 7, and ligand - side chain interactions are elaborated upon in supplementary file 8.

While both MYM-III-10 and MYM-V-56 were less sedating than MCLZ, both of these compounds still retained positive allosteric modulator activity against GABA_A_Rs that would be expected to confer sedation (**Figure 5 & 6**). So why were MYM-III-10 and MYM-V-56 not as sedating as MCLZ? For MYM-III-10, an explanation may relate to pharmacokinetic properties of this molecule, since it is more polar with a hydroxyl group at the C3 position. A lower ClogP would be expected to have less blood brain barrier penetration and sedation (40), which is indeed what is observed when levels of this compound were measured in brain tissue (**Figure 6C**).

While there is a plausible explanation for why MYM-III-10 is much less sedating than MCLZ despite GABA_A_R affinity, it is less clear why MYM-V-56 is not sedating. The activity of MYM-V-56 on GABA_A_Rs does differ slightly from MCLZ and MYM-III-10. MYM-V-56 does not have quite as potent affinity as these two compounds for the central benzodiazepine binding site in radioligand binding assays (**Figure 6A**), but does still bind with relatively high affinity. MYM-V-56 also displays potent PAM activity at recombinantly expressed GABA_A_Rs (**Figure 6B**), even though receptors desensitize at higher concentrations. From these data we may still expect that MYM-V-56 would be sedating, but this was not the case in the rotarod assay even at doses up to 100 mg / kg (**Figure 5**). However, from observing these mice it was clear that MYM-V-56 does have unique psychoactive effects (**Supplementary File 3**). These mice were hyperactive, indicating MYM-V-56 may have activity on targets other than GABA_A_Rs, or GABA_A_Rs with different subunit compositions than the typical ɑ1-containing channels that cause sedation (e.g. ɑ2-containing channels that mediate anxiolytic effects). The identity of what those receptors may be is not clear and remains a topic for future investigation (**Supplementary Table 2**). Indeed, while the rotarod data showing a lack of sedation appears promising for this compound, the distinct psychoactive effects observed would likely not be acceptable for further development. Exploration of additional molecules with C3 modifications is warranted to determine whether related compounds (e.g. different halogens or other electrophilic groups) would have the same effect.

The structure activity relationship of meclonazepam derivatives explored in this study, as well as the comparative modeling of the host and parasite receptors and the identification of two new lead molecules, will advance benzodiazepines as promising candidates for development as new antischistosomal therapies. This class of compounds displays excellent anti-parasitic activity, through mechanisms distinct from the existing monotherapy, praziquantel. Benzodiazepines also have an established track record as highly prescribed drugs (>5% of adults in the United States) with known safety profiles (41). The primary barrier so far to developing these compounds to treat schistosomiasis has been dose-limiting sedation, which we demonstrate can be reduced or eliminated in a murine model.

## Materials and methods

### Chemical synthesis

Meclonazepam was purchased from Anant Pharmaceuticals and verified as greater than 99% pure by HPLC. For HPLC experiments, a meclonazepam reference standard was purchased from Sigma Aldrich (catalog number M-197-1ML). The reactions were performed in round-bottom flasks with magnetic stir bars under an argon atmosphere or air condition. Organic solvents were purified when necessary by standard methods or purchased from Sigma-Aldrich Chemicals. Reagents and other chemicals were purchased from either Sigma-Aldrich, Oakwood Chemical, Alfa Aesar, Matrix Scientific, Admiral Chemical Company, or Acros Organic. The progress of reactions was visualized with TLC plates from Dynamic Adsorbents, Inc. under a UV light. LCMS 2020 was used to monitor some reactions. Flash column chromatography was done for purification of some analogs on silica gel (230-400 mesh, Dynamic Adsorbents). A normal phase Agilent HPLC was used to determine the ratio of optically active enantiomers as well as to determine %ee. The ^1^H NMR and ^13^C NMR spectra were obtained on Bruker Spectrospin 500 MHz instrument in CDCl_3_ and chemical shifts were reported in δ (ppm). Multiplicities are represented as follows: singlet (s), broad signal (br), doublet (d), triplet (t), quartet (q), dd (doublet of doublets), and multiplet (m). The technique employed for HRMS was carried out on a LCMS-IT-TOF at the Milwaukee Institute for Drug Discovery in the Shimadzu Laboratory for Advanced and Applied Analytical Chemistry. Detailed synthesis methods, reaction schemes and NMR spectra for compounds shown in Figure 1 are provided in **Supplementary File 1**.

### Purification of enantiomers

MYM-III-10 and MYM-V-56 enantiomers were purified by preparative HPLC. The (R) and (S) enantiomers of MYM-III-10 were separated by using an isocratic mixture of 88% n-hexane and 12% ethanol as the mobile phase in an HPLC system consisting of a quaternary pump, auto sampler and a DAD detector. The method time was set for 55 min. A chiral preparative HPLC column, Reflect I cellulose C 5 µm, 25 cm x 21.1 mm was used for separating the enantiomers. The first fraction (*R* isomer) for MYM-III-10 was collected from 35.2 to 40.2 min and the second fraction was collected from 45.5 to 50 min. Approximately, 15 mL of sample was collected for each fraction after each injection. The fractions were evaporated separately just after collection and dried. Compounds were then analyzed by NMR. The DAD wavelength range was set at 200-400 nm. A 254 nm wavelength was used for relative % area determination. The flow rate was set at 21 mL/min.

The (R) and (S) enantiomers of MYM-V-56 were separated using the same method, but set for 25 min. The first fraction (*R* isomer) for MYM-V-56 was collected from 19.5 min to 21.2 min and the second fraction was collected from 22.4 min to 23.4 min. Approximately, 15 mL of sample was collected for each fraction after each injection. The fractions were evaporated separately just after collection, dried, and analyzed by NMR.

To confirm the absolute configuration of MYM-V-56, the X-ray crystal structure of the first fraction was determined and it turned out to be the (R) configuration, indicating the other enantiomer is (S)-configuration. A clear colorless chunk crystal of dimensions 0.236 x 0.185 x 0.080 mm was mounted on a MiteGen MicroMesh using a small amount of Cargille Immersion Oil. Data were collected on a Bruker three-circle platform diffractometer equipped with a PHOTON II CPAD detector. The crystals were irradiated using a 1μs microfocus CuK_a_ source (l = 1.54178) with Montel optics. Data was collected at room temperature (20°C) and the unit cell was initially refined using *APEX3* [v2015.5-2](42). Data Reduction was performed using *SAINT* [v8.34A] and *XPREP* [v2014/2] (Bruker AXS Inc.). Corrections were applied for Lorentz, polarization, and absorption effects using *SADABS* [v2014/2] and the structure was solved and refined with the aid of SHELXL-2014/7 (Bruker AXS Inc.). The full-matrix least-squares refinement on F^2^ included atomic coordinates and anisotropic thermal parameters for all non-H atoms. Hydrogen atoms were located from the difference electron-density maps and added using a riding model.

### *In vitro* antischistosomal activity assays

Female Swiss Webster mice infected with NMRI strain *S. mansoni* were provided by the NIH-NIAID Schistosomiasis Resource Center and euthanized by CO_2_ asphyxiation and cervical dislocation 6-7 weeks post-infection. Adult schistosomes were harvested from the mesenteric vasculature and placed in culture media consisting of high-glucose DMEM supplemented with 5% fetal calf serum, penicillin-streptomycin (100 units/mL), HEPES (25 mM) and sodium pyruvate (1 mM) and cultured at 37°C / 5% CO_2_.

Worms were treated with either DMSO control or test compounds listed in Supplementary Table 1 for 14 hours at 37°C / 5% CO_2_ prior to observing phenotypes. Worms were visually observed on a Zeiss Stemi 305 Stereo Microscope to assess changes in morphology and images were captured on an Axiocam 208 color camera. For motility experiments, serotonin (5-HT) was added at 250 µM to increase worm movement at least 1 hour prior to imaging on an ImageXpress Nano (Molecular Devices). Time lapse recordings were acquired for each well (15 second videos at a rate of 4 frames / second) using a 2X objective. Movement was analyzed using the wrmXpress package (25).

All animal work was carried out with the oversight and approval of UW-Madison Research Animal Resources and Compliance (RARC), adhering to the humane standards for the health and welfare of animals used for biomedical purposes defined by the Animal Welfare Act and the Health Research Extension Act. Experiments were approved by the UW-Madison School of Veterinary Medicine IACUC committee (approved protocol #V006353-A08).

### *In vivo* antischistosomal activity assays

For hepatic shift assays, mice harboring seven week old *S. mansoni* infections were administered either DMSO control, MCLZ or test compound solubilized in vegetable oil (0.25 mL) by oral gavage and then euthanized as outlined above (CO_2_ asphyxiation followed by cervical dislocation) three hours later. Worms were dissected from the liver and mesenteries, and the percentage of worms in each location was calculated. For longer term assays measuring curative activity of drugs, compounds were administered orally to mice harboring six week old *S. mansoni* infections. Mice were euthanized one week later, and the total parasite burden for each mouse was counted. To assess the curative effects of compounds on juvenile, liver stage parasites mice harboring four week old *S. mansoni* infections were dosed orally with test compound and euthanized at seven weeks to count parasite burden.

### Rotarod assay for sedation

Six week old female Swiss Webster mice were chosen for study since this matches the strain used in the parasite assays. Uninfected mice were ordered from taconic or charles river and trained to perform the rotarod test using a IITC Life Science rotarod device with the following settings: 20 RPM max speed, 20 ramp speed, and 300 second duration test. Mice were run five at a time and only used for drug assays if they were capable of completing the training and able to run on the device for the entire 300 seconds. Mice were then dosed with test compounds by oral gavage and tested on the rotarod device after 5 minutes.

### Detection of MCLZ and MYM-III-10 in brain and serum

Six week old female Swiss Webster mice were dosed with MCLZ or MYM-III-10 (100 mg / kg) as described in the antischistosomal assays and then euthanized after 10 minutes by CO_2_ asphyxiation. Whole blood samples were collected from each mouse and allowed to clot at room temperature for 15 minutes. Serum was separated by centrifugation at 2,000 x g for 10 minutes. The skull was opened with scissors and all brain tissue was removed. Both samples were frozen at -80°C until ready for analysis by HPLC.

To prepare samples for HPLC brain tissue homogenized with BeadBug microtube homogenizer and the solid phase extraction procedure was carried out on Hypersep C18 cartridges. Separations were obtained on Shimadzu HPLC System with Waters Symmetry C18 column (150x3.9 mm id, 5 μm) kept at 30°C. The mobile phase was composed of a mixture of MeOH (65% v/v) + 0.1% Formic acid and a ultrapure water (35% v/v) + 0.1% Formic acid. The flow rate was set at 0.5 mL/min. Detection was carried out on a Shimadzu Diode Array Detector SPD-M20A at 254 nm. Spiked in meclonazepam used as an internal standard for MYM-III-10 treated samples, and vice versa.

### Binding and functional assays

MYM-V-56 binding was assessed by two panels of assays. The Safety Screen 44 Panel was performed by Eurofins Pharma Discovery. Per vendor recommendations, each assay was performed using 10 µM of MYM-V-56 in technical duplicates. MYM-V-56 (10 µM) binding to receptors in the NIMH Psychoactive Drug Screening Program (PDSP) in technical quadruplicate per SOPs in the PDSP assay protocol book found at https://pdsp.unc.edu/pdspweb/

### Modeling benzodiazepine binding to TRP and GABA_A_Rs

Targets were obtained from public structure repositories. GABA_A_R coordinates were taken from RSCB, PDB accession 6HUP for a cryo-EM derived structure in its diazepam-bound state. The AlphaFold model AF-A0A5K4FCC0-F1-model provided starting coordinates for the *S. mansoni* TRP channel Smp_333650. Structure models were prepared for docking using the DockPrep utility of ChimeraX v1.7 (43) to fill in missing atoms, add hydrogens, assign partial charges (AMBER ff14SB), and exported in mol2 formats. SMILES for the benzodiazepine chemical series were generated using ChemDraw. Library preparation utilities fixpka (QUACPAC v2.1.2.1), OMEGA2 v4.1.1.1, and molcharge (QUACPAC v2.1.2.1) were used to assign protonation states, build energetically reasonable starting 3D conformers for docking, and assign partial charges (MMFF), respectively (OpenEye Scientific, Cadence Molecular Sciences, Inc.). Single conformations were exported in mol2 format for docking. Docking was performed using the program Gnina v1.0 (44). To localize the search space in docking GABA_A_R, the binding pocket was specified based on the position of diazepam using the autobox option with a bounding box of 10 Angstroms. No sidechain flexibility was allowed for docking into the GABA_A_R. For the schistosome TRP channel, the site center was determined using the most probable site in the VSLD domain using the prediction tool p2rank (45). Due to steric restrictions within the putative binding pocket of the TRP channel, sidechain flexibility was enabled for pocket residues H827, H831, Y864, E867, E868, D893, F940, R943, H946, I947, Y1117, R1120. For both targets, extensive sampling was achieved using an exhaustiveness parameter of 64. CNN scoring was used for re-scoring final poses by setting cnn_scoring option to “rescore.” For CNN-based pose scoring, the “crossdock_default2018” model was selected. Docking poses were inspected and images created using PyMOL v2.5.4 (Schrodinger, LLC).

## Acknowledgments.

This work was supported by funding from NIH-NIAID R21AI146540 (JDC) and a University of Wisconsin System Regent Scholar Award (JDC and JMC). JMC is also supported by R01DA011792, R01DA043204 and R01NS076517, Office of Naval Research grant N00014–15-WX-0–0149 and NSF Chemistry Research Instrumentation Program award #1625735. We acknowledge the UW-Milwaukee Shimadzu Laboratory for Advanced and Applied Analytical Chemistry and support from the Milwaukee Institute of Drug Discovery and the University of Wisconsin-Milwaukee Research Foundation. Schistosome infected mice were provided by the NIH-NIAID Schistosomiasis Resource Center for distribution through BEI Resources, NIH-NIAID Contract HHSN272201700014I. Receptor binding profile for MYM-V-56 was generously provided by the National Institute of Mental Health’s Psychoactive Drug Screening Program, Contract # HHSN-271-2018-00023-C (NIMH PDSP). We thank Jonathan Marchant for sharing the pdb file of meclonazepam docked into Smp_333650, facilitating *in silico* modeling of the compounds reported in this study.

---

**Supplementary File 1. Supporting data on synthesis and purification of meclonazepam derivatives.** Detailed synthesis methods along with HPLC chromatograms, X-ray crystal structure data, and NMR spectra for benzodiazepines screened.

**Supplementary Files 2 - 4. Videos of mice treated with vehicle control, MCLZ, or MYM-V-56**. Mice were given either vehicle control or either MCLZ or MYM-V-56 (100 mg / kg) by oral gavage.

**Supplementary File 5. Benzodiazepines modeled in the human GABA_A_R**. Compounds from Figure 1A are shown docked in the diazepam binding site of the human α1β3γ2L GABAAR. Nine poses are shown, ranked in order of gnina docking score (values provided in Supplementary File 7).

**Supplementary File 6. Benzodiazepines modeled in the *S. mansoni* TRP.** Compounds from Figure 1A are shown docked in the VSLD cavity of Smp_333650. Nine poses are shown, ranked in order of gnina docking score (values provided in Supplementary File 7).

**Supplementary File 7. Scores for benzodiazepines docked into the human GABA_A_R and *S. mansoni* TRP channel**. Docking scores from gnina for the 9 poses generated for each benzodiazepine are shown.

**Supplementary File 8. LigPlot visualization of meclonazepam binding in the human GABA_A_R and *S. mansoni* TRP channel.** Plots were generated using LigPlot+ v2.2.8 for the top meclonazepam poses in Supplementary Files 5 and 6.

**Supplementary Table 1. Structures of test compounds.** Structure and SMILES ID provided for each compound.

**Supplementary Table 2. Binding of MYM-V-56 to human receptors.** (S)-MYM-V-56 was screened at a concentration of 10 µM against the Eurofins SafetyScreen panel (Sheet 1, 44 targets consisting of 36 binding assays and 6 functional assays) and the NIMH Psychoactive Drug Screening Program (PDSP; Sheet 2 - 46 assays). Binding is indicated by percent inhibition of radioligand binding caused by MYM-V-56.

