## Supplementary figures and images for "Development of non-sedating antischistosomal benzodiazepines"

### Supplementary File 8

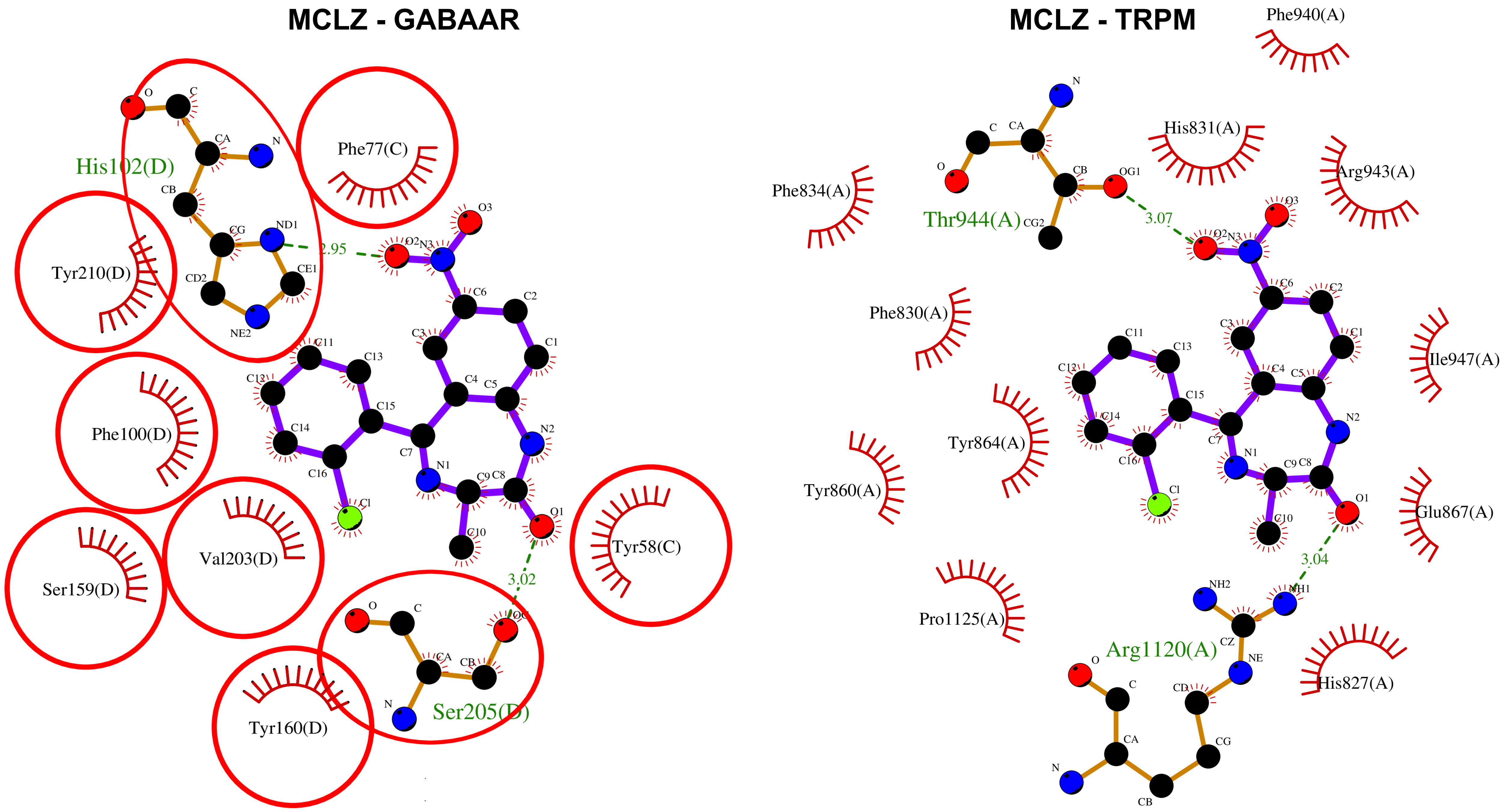
