## Supplementary File 1 for "Development of non-sedating antischistosomal benzodiazepines"

**Supplementary File 1. Supporting data on synthesis and purification of meclonazepam derivatives.**

**Scheme 1. Synthesis of N-methyl meclonazepam**

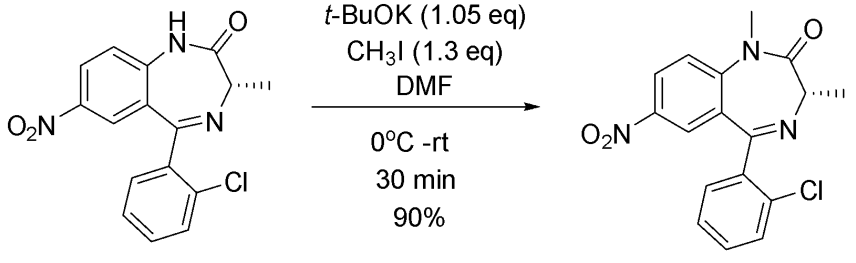


**1** **2**, MYM-II-53

**(*S*)-5-(2-chlorophenyl)-1,3-dimethyl-7-nitro-1,3-dihydro-2H-benzo[e][1,4]diazepin-2-one (2, MYM-II-53)**

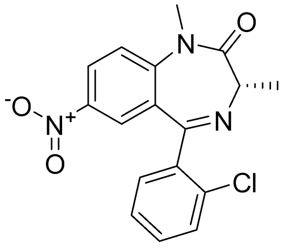

**2** (MYM-II-53)

An oven dried 3 neck round bottom flask was charged with meclonazepam **15** (80 mg, 0.243 mmol) (Anant Pharmaceuticals Pvt. Ltd., India), anhydrous *N*,*N*-dimethyl formamide (5 ml) and methyl iodide (0.02 ml, 0.32 mmol). The mixture was stirred for 10 min until homogeneous and cooled to -30 °C. Then potassium *tert*-butoxide (27 mg, 0.254 mmol) was added to the reaction mixture. The solution was stirred at 25 °C for 2 hours. The reaction was monitored by TLC. After completion, the reaction was quenched by adding ice cold water and then extracted with ethyl acetate (3 × 10 mL). The combined organic layer was washed with 25% aq. ammonium chloride (1 × 10 mL), brine (3 × 10 mL) and dried ($Na_2SO_4$). The solvent was removed under reduced pressure and the solid obtained was purified by flash chromatography (neutral alumina, EtOAc: hexane, 3:2) to afford pure methylated meclonazepam **2** as yellow solid (75 mg, 91%). $^1$H NMR (500 MHz, $CDCl_3$) δ 8.38 (dd, *J* = 9.1, 2.7 Hz, 1H, Ar), 7.96 (d, *J* = 2.6 Hz, 1H, Ar), 7.71 – 7.65 (m, 1H, Ar), 7.50 (d, *J* = 9.1 Hz, 1H, Ar), 7.47 – 7.43 (m, 2H, Ar), 7.39 – 7.35 (m, 1H, Ar), 3.80 (d, *J* = 6.5 Hz, 1H, CH), 3.55 (s, 3H, N-$\underline{CH_3}$), 1.78 (d, *J* = 6.5 Hz, 3H, $CH_3$). $^{13}$C NMR (126 MHz, $CDCl_3$) δ 170.31, 166.62, 147.78, 143.12, 137.22, 133.00, 131.59, 131.27, 130.78, 130.29, 127.47, 126.09, 124.16, 122.06, 59.09, 35.33, 17.28. **HRMS (ESI/ITTOF) m/z: $[M + H]^+$** Calcd for $C_{17}H_{14}N_3O_3Cl$ 344.0796; found 344.0792.

**Scheme 2. Synthesis of C-3 modified analogs of meclonazepam**

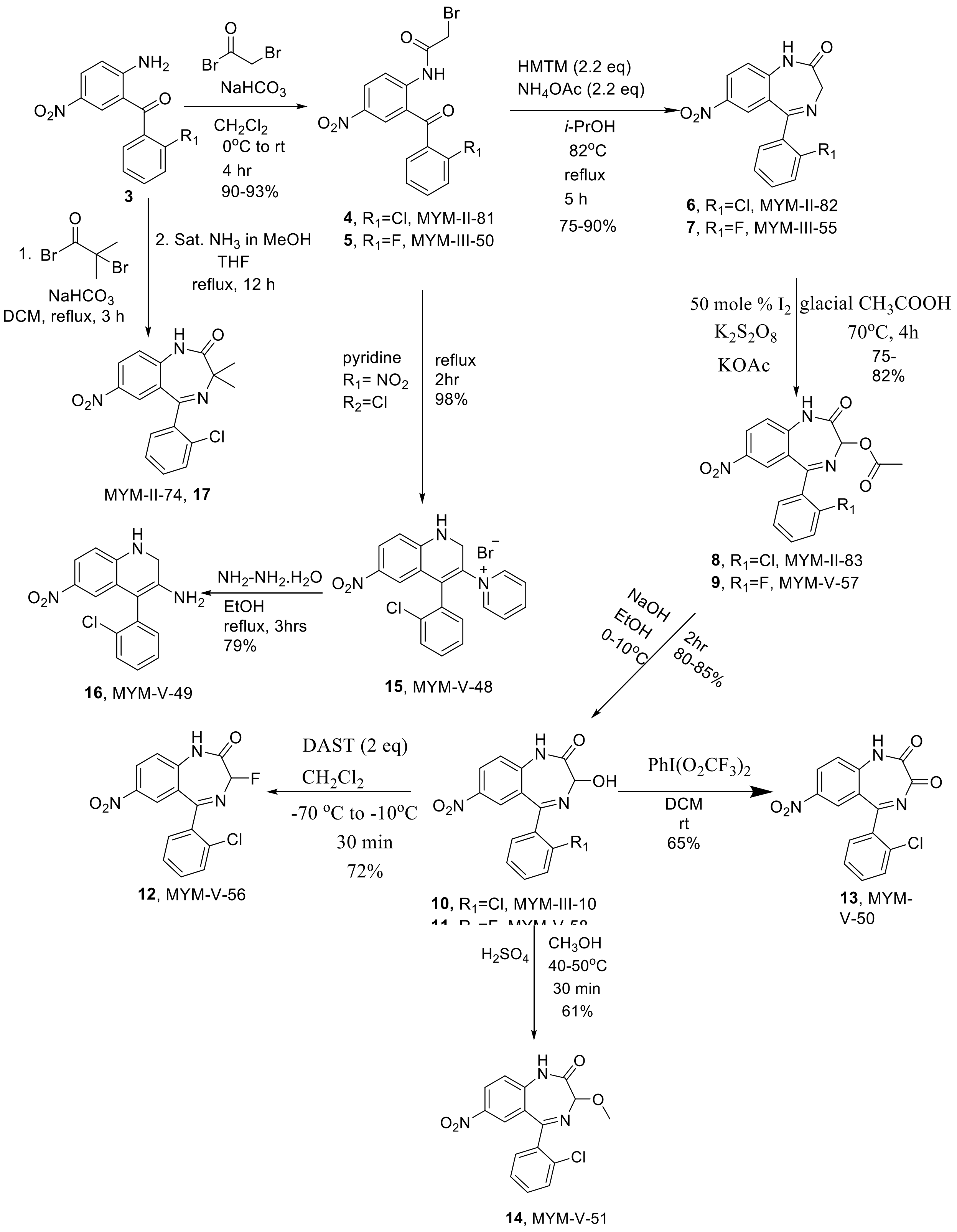

**2-Bromo-N-(4-bromo-2-(2-chlorobenzoyl)phenyl)acetamide (4, MYM-II-81)**

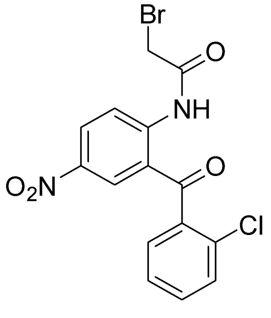

**4**

A round bottom flask was charged with (2-amino-5-nitrophenyl)(2-chlorophenyl)methanone ( 30 g, 108 mmol) and anhydrous DCM (300 mL) and stir to make a homogenous solution. The sodium bi-carbonate (18.15 g, 216 mmol, solid) was added once to the reaction mixture. Bromoacetyl bromide (19.6 mL, 224.4 mmol) was added dropwise to the mixture at rt over a period of 30 min. The mixture was the stirred for additional 3 hours after complete addition of bromoacetyl bromide. The completion of the reaction was confirmed by silica gel TLC (30% EtOAc-hexane). The reaction mixture was quenched slowly by adding water (100 mL) over a 10 min as carbon dioxide gas evolved. The biphasic mixture, which resulted, was allowed to stand for 5 min and the layers were separated. The organic layer was separated and the aq layer was extracted with dichloromethane (2x100 mL).The combined organic layers were washed with 5% aq sodium bicarbonate solution (1x 250 mL), 10% aq sodium chloride solution (2x 250 mL) and dried ($Na_2SO_4$). The residue was slurried in ethanol (150 mL) and stirred for 20 minutes at 50°C. Upon cooling to rt, the residue was filtered, washed with cold ethanol (2x20 mL), and dried under vacuum at 40 °C to afford the product **4** as an off-white powder (42 g, 97.5%). **$^1$H NMR** (500 MHz, $CDCl_3$) δ 12.35 (s, 1H), 8.99 (d, *J* = 9.3 Hz, 1H), 8.45 (dd, *J* = 9.3, 2.7 Hz, 1H), 8.29 (d, *J* = 2.7 Hz, 1H), 7.59 – 7.52 (m, 2H), 7.49 – 7.46 (m, 1H), 7.42 (dd, *J* = 7.6, 1.4 Hz, 1H), 4.12 (s, 2H); **$^{13}$C NMR** (126 MHz, $CDCl_3$) δ 197.76 (s), 166.00 (s), 145.52 (s), 142.28 (s), 137.03 (s), 132.48 (s), 131.09 (s), 130.58 (s), 130.13 (s), 129.50 (s), 129.09 (s), 127.34 (s), 121.92 (s), 121.10 (s), 29.26 (s). **HRMS (ESI/IT-TOF**) m/z: [M - H]$^-$ Calcd for $C_{15}H_{10}$N2$O_4$Cl 394.9440 found 394.9428.

**2-Bromo-N-(4-bromo-2-(2-fluorobenzoyl)phenyl)acetamide (5, MYM-III-50)**

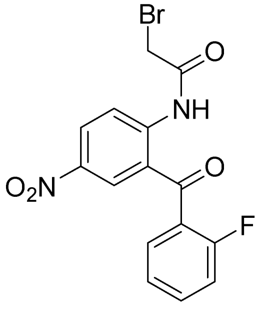

**5**

Bromoacetyl bromide (3.3 mL, 27 mmol) was added dropwise to the mixture of (2-amino-5-nitrophenyl)(2-fluorophenyl)methanone (7 g, 27 mmol), solid sodium bicarbonate (3.4 g, 40.5 mmol), and anhydrous dichloromethane (70 mL) at 0°C. The mixture was the stirred for 4 hour after complete addition of bromoacetyl bromide. The completion of the reaction was confirmed by silica gel TLC (EtOAc: hexane = 3:7). The reaction mixture was quenched slowly by adding water (50 mL) over a 10 min as carbon dioxide gas evolved. The biphasic mixture, which resulted, was allowed to stand for 5 min and the layers were separated. The organic layer was separated and the aq layer was extracted with dichloromethane (2x50 mL) and the combined organic layers were washed with 5% aq sodium bicarbonate solution (1x 50 mL), 10% aq sodium chloride solution (2x 50 mL) and dried ($Na_2SO_4$). The residue was slurried in EtOH (40 mL) and stirred for 20 min at 50°C. Upon cooling to rt and holding for 2 h, the residue was filtered, washed with EtOH (2x10 mL), and dried under vacuum at 40 °C to afford the product **5** as an off-white powder (9.5 g, 92.5%). **$^1$H NMR** (500 MHz, $CDCl_3$) δ 12.17 (s, 1H), 8.97 (d, *J* = 9.9 Hz, 1H), 8.47 (dd, *J* = 5.4, 2.4 Hz, 2H), 7.70 – 7.64 (m, 1H), 7.62 – 7.58 (m, 1H), 7.38 (t, *J* = 7.6 Hz, 1H), 7.26 (t, *J* = 9.2 Hz, 1H), 4.11 (s, 2H). **$^{13}$C NMR** (126 MHz, $CDCl_3$) δ 195.30 (s), 165.84 (s), 159.64 (d, *J* = 253.4 Hz), 144.97 (s), 142.38 (s), 134.69 (d, *J* = 8.5 Hz), 130.63 (d, *J* = 1.9 Hz), 129.78 (s), 129.11 (d, *J* = 2.9 Hz), 125.92 (d, *J* = 14.0 Hz), 125.02 (d, *J* = 3.6 Hz), 122.97 (s), 121.17 (s), 116.81 (d, *J* = 21.5 Hz), 29.18 (s). **HRMS (ESI/IT-TOF)** m/z: $[M+H]^+$ Calcd for $C_{15}H_{10}N_2O_4F$ 380.98807 found 380.98852.

**7-Nitro-5-(2-chlorophenyl)-1,3-dihydro-2H-benzo[e][1,4]diazepin-2-one (6, MYM-II-82)**

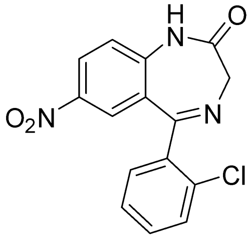


**6**

A round bottom flask was charged with 2-bromo-N-(4-nitro-2-(2-chlorobenzoyl) phenyl)acetamide (**57**, 70 g, 177 mmol), IPA (400 mL), HMTM (54.58 g, 390 mmol), ammonium acetate (30.07 g, 390 mmol) and the mixture was refluxed for 4 h at 82°C at which point the reaction was deemed to complete on TLC analysis (silica gel, EtOAc: Hexane=1:1). The reaction mixture was then cooled to -10 to -5°C and hold for 2 hrs. The residue was filtered, washed cold IPA (100 mL), and water (4x200 mL). The solid was dried under vacuum at 40°C for 6 h to afford pure **6** as an off white (44.4 g, 80%). **$^1$H NMR** (500 MHz, DMSO) δ 11.31 (s, 1H), 8.37 (dd, *J* = 9.0, 2.7 Hz, 1H), 7.75 (d, *J* = 2.6 Hz, 1H), 7.65 (dt, *J* = 7.7, 3.5 Hz, 1H), 7.55 – 7.52 (m, 1H), 7.51 – 7.48 (m, 1H), 7.45 (dd, *J* = 9.0, 3.4 Hz, 1H), 4.32 (s, 2H). **$^{13}$C NMR (**126 MHz, DMSO) δ 169.77 (s), 168.36 (s), 144.78 (s), 142.07 (s), 138.45 (s), 132.27 (s), 132.00 (s), 131.93 (s), 130.28 (s), 128.00 (s), 127.41 (s), 126.91 (s), 125.36 (s), 122.62 (s), 57.55 (s). **HRMS** (LCMS-IT-TOF) Calc. for $C_{15}H_{10}N_3O_3Cl$ $(M-H)^-$ 314.0338, found 314.0316.

**7-Nitro-5-(2-fluorophenyl)-1,3-dihydro-2H-benzo[e][1,4]diazepin-2-one (7, MYM-III-55)**

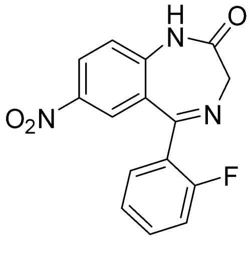


**7**

Compound **7** (7.6 g, 80%) was synthesized from 2-bromo-N-(4-nitro-2-(2-fluorobenzoyl) phenyl) acetamide (**5**, 9.5 g, 23 mmol), IPA (100 mL), HMTM (7.09 g, 50.3 mmol), and ammonium acetate (3.87 g, 50.3 mmol) according to the procedure described for the synthesis of compound **6**. **$^1$H NMR** (500 MHz, DMSO) δ 11.27 (s, 1H), 8.37 (d, *J* = 8.9 Hz, 1H), 7.90 (d, *J* = 2.0 Hz, 1H), 7.65 – 7.55 (m, 2H), 7.45 (d, *J* = 9.0 Hz, 1H), 7.35 (t, *J* =

7.4 Hz, 1H), 7.24 (t, *J* = 9.3 Hz, 1H), 4.30 (s, 2H). **$^{13}$C NMR** (126 MHz, DMSO) δ 169.91 (s), 165.64 (s), 160.16 (d, *J* = 249.0 Hz), 144.30 (s), 142.21 (s), 133.22 (d, *J* = 8.2 Hz), 132.11 (s), 127.68 (s), 127.16 (d, *J* = 12.2 Hz), 126.87 (s), 125.68 (s), 125.26 (s), 122.71 (s), 116.55 (d, *J* = 21.4 Hz), 57.63 (s). **HRMS** (LCMS-IT-TOF) Calc. for $C_{15}H_{10}N_3O_3F$ $(M + H)^+$ 300.0779, found 300.0782.

**7-Nitro-5-(2-chlorophenyl)-2-oxo-2,3-dihydro-1H-benzo[e][1,4]diazepin-3-yl acetate (8, MYM-II-83)**

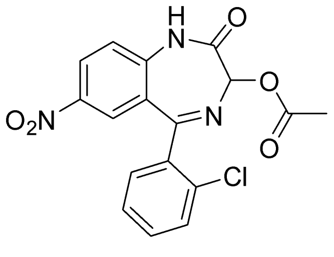

**8**

A round bottom flask was charged with potassium acetate (13.05 g, 133 mmol), iodine (8.44, 33 mmol), 7-nitro-5-(2-chlorophenyl)-1,3-dihydro-2H-benzo[e][1,4]diazepin-2-one (**61**, 21 g, 66.5mmol), glacial acetic acid (200 mL) and the mixture was then allowed to heat at 65°C. Then potassium persulfate (35.96 g, 133 mmol) was added to the reaction mixture at several portions. The reaction mixture was stirred at 65-70°C for additional 5 h. At that point the reaction was deemed to complete on TLC analysis (silica gel, 50% EtOAc-hexane). The acetic acid and iodine were evaporated which resulted a purple color gummy mass. A solution of sodium thiosulfate (40 g) in water (150 mL) was added to the gummy mass and the mixture was at 70°C for 2 h. Upon cooling to 10°C, the residue was filtered, washed with hot water (2x100 mL), and dried under vacuum at 50°C for 4 h. The orange-colored crude products, which resulted was then dissolved in hot DMF (70-80°C, 100 mL) and filtered to remove any inorganic materials. Then IPA (150 mL) was added dropwise very slowly to the clear DMF solution at 70°C and stirred for 30 minutes. The mixture was cooled to rt and kept in the refrigerator (-20°C) for 3 h. The fluffy materials, which resulted was filtered, washed with IPA (2x30 mL), and dried under vacuum at 40°C for 3 h to afford **8** as a white powder (20.4, 82%). **$^1$H NMR** (500 MHz, DMSO) δ 11.77 (s, 1H), 8.44 (dd, *J* = 9.0, 2.6 Hz, 1H), 7.78 (d, *J* = 2.6 Hz, 1H), 7.70 (dd, 1H), 7.62 – 7.55 (m, 2H), 7.55 – 7.49 (m, 2H), 5.96 (s, 1H), 2.23 (s, 3H). **$^{13}$C NMR** (126 MHz, DMSO) δ 170.00 (s), 165.01 (s), 164.66 (s), 143.47 (s), 142.56 (s), 137.33 (s), 132.52 (s), 132.31 (s), 132.04 (s), 130.43 (s), 128.15 (s), 127.65 (s), 127.38 (s), 125.16 (s), 123.10 (s), 85.36 (s), 21.14 (s). **HRMS** (LCMS-IT-TOF) Calc. for $C_{17}H_{12}N_3O_5Cl$ $(M + H)^+$ 374.0538, found 374.0562.

**5-(2-Chlorophenyl)-3-hydroxy-7-nitro-1,3-dihydro-2H-benzo[e][1,4]diazepin-2-one (10, MYM-III-10)**

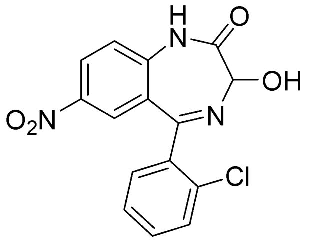

**10, MYM-III-10**

A solution of NaOH (4.28 g, 107 mmol) in water (30 mL) was added dropwise to the stirred solution of 7-nitro-5-(2-chlorophenyl)-2-oxo-2,3-dihydro-1H-benzo[e][1,4]diazepin-3-yl acetate (**65**, 20 g, 53.5 mmol) in ethanol (200 mL) over a period of 30 min. The reaction mixture was then stirred for 1h at +5 to 10°C at which point the reaction was deemed to complete on TLC analysis (silica gel, 100% EtOAc). Glacial acetic acid (10 mL) was then added dropwise to the reaction mixture. The mixture was cooled to 0 to 5$^o$C. The residue was filtered, washed with water (3x 30 mL), and dried under vacuum. The crude products were then slurried in ethanol and refluxed for 20 min. The mixture was cooled to 0 to 5$^o$C and hold for 1 h. The residue was filtered, washed with cold ethanol (2x20 mL), and dried under vacuum for 2 h at 40$^o$C to afford pure **10** as an off-white powder (14.7 g, 83%). **$^1$H NMR** (500 MHz, DMSO) δ 11.44 (s, 1H), 8.39 (dd, *J* = 9.0, 2.6 Hz, 1H), 7.77 (d, *J* = 2.6 Hz, 1H), 7.70 (dd, *J* = 5.8, 3.4 Hz, 1H), 7.58 – 7.54 (m, 2H), 7.52 – 7.49 (m, 1H), 7.44 (d, *J* = 9.1 Hz, 1H), 6.63 (d, *J* = 7.1 Hz, 1H), 4.96 (s, 1H). **$^{13}$C NMR** (126 MHz, DMSO) δ 169.62 (s), 144.06 (s), 142.16 (s), 137.93 (s), 132.27 (s), 132.07 (s), 131.96 (s), 130.32 (s), 128.05 (s), 127.49 (s), 127.10 (s), 124.88 (s), 122.69 (s), 83.59 (s), 56.49 (s), 19.03 (s). **HRMS** (LCMS-IT-TOF) Calc. for $C_{15}H_{10}N_3O_4Cl$ $(M - H)^-$ 330.0287, found 330.0266.

**5-(2-Fluorophenyl)-3-hydroxy-7-nitro-1,3-dihydro-2H-benzo[e][1,4]diazepin-2-one (11, MYM-V-58)**

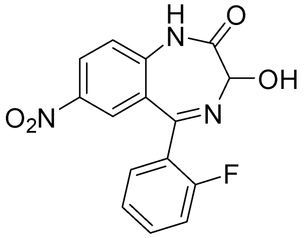

**11, MYM-V-58**

A round bottom flask was charged with 5-(2-fluorophenyl)-7-nitro-2-oxo-2,3-dihydro-1H-benzo[e][1,4]diazepin-3-yl acetate (**66**, 0.5 g, 1.34 mmol), ethanol (10 mL) and stirred to make a suspension. A solution of NaOH (0.13 g, 3.21 mmol) in water (2 mL) was added dropwise to the reaction mixture. The reaction mixture was then stirred for 30 min at +5 to 10$^o$C at which point the reaction was deemed to complete on TLC analysis (silica gel, 100% EtOAc). Glacial acetic acid (1 mL) was then added dropwise to the reaction mixture. The mixture was cooled to 0 to 5$^o$C. The residue was filtered, washed with water (2x 5 mL), and dried under vacuum. The crude products was purified by a flash chromatography (silica gel, EtOAc) to afford pure **11** as a yellow colored powder (308 mg, 70%). **$^1$H NMR** (500 MHz, DMSO) δ 11.40 (s, 1H), 8.40 (dd, *J* = 9.0, 2.6 Hz,1H), 7.92 (d, *J* = 2.3 Hz, 1H), 7.69 – 7.57 (m, 2H), 7.45 (d, *J* = 9.0 Hz, `H), 7.39 (t, *J* = 7.4 Hz, 1H), 7.27 (dd, *J* = 10.3, 8.8 Hz, 1H), 6.65 (d, *J* = 8.7 Hz, 1H), 4.93 (d, *J* = 8.4 Hz, 1H). **HRMS** (LCMS-IT-TOF) Calc. for $C_{15}H_{10}N_3O_4F$ $(M + H)^+$ 316.07281, found 316.0727.

**5-(2-Chlorophenyl)-7-nitro-1H-benzo[e][1,4]diazepine-2,3-dione (12, MYM-V-50)**

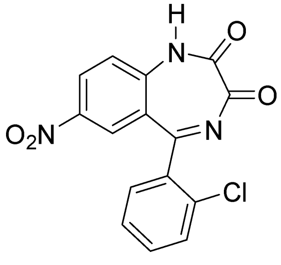

**12, MYM-V-50**

Bis(trifluoroacetoxy)iodobenzene (0.25 g, 0.60 mmol) was added to the stirred solution of 5-(2-chlorophenyl)-3-hydroxy-7-nitro-1,3-dihydro-2H-benzo[e][1,4]diazepin-2-one (**10**, 0.1 g, 0.3 mmol) in anhydrous acetonitrile (10 mL). The mixture was stirred for 5 h at which point the reaction was deemed to complete on TLC analysis (silica gel, 50% EtOAc-hexane, $R_f$=0.4; $R_f$=0.3 for **10**). The reaction was then quenched with water (10 mL) and the aq. layer was extracted with DCM (2x10 mL). The combined organic layer was washed with brine

(2x10 mL), and dried ($Na_2SO_4$). The solvents were evaporated under reduced pressure and the residue was purified by a flash chromatography (silica gel, 40% EtOAc-hexane) to afford **12** as white solid (60.8 mg, 61%). **$^1$H NMR** (500 MHz, DMSO) δ 8.82 (ddd, *J* = 9.6, 7.2, 2.5 Hz, 1H), 8.52 (dd, *J* = 16.4, 9.3 Hz, 1H), 8.42 – 8.36 (m, 1H), 7.81 (d, *J* = 8.1 Hz, 1H), 7.77 – 7.70 (m, 2H), 7.67 (dd, *J* = 10.6, 4.0 Hz, 1H). **$^{13}$C NMR** (126 MHz, $CDCl_3$) δ 171.13 (s), 170.96 (s), 163.77 (s), 154.71 (s), 152.65 (s), 134.17 (s), 132.38 (s), 132.12 (s), 132.07 (s), 131.24 (s), 131.21 (s), 130.44 (s), 128.17 (s), 127.60 (s), 123.85 (s). **HRMS** (LCMS-IT-TOF) found for $C_{15}H_9N_3O_4Cl$ $(M - H)^-$ 329.0287.

**5-(2-Chlorophenyl)-3-fluoro-7-nitro-1,3-dihydro-2H-benzo[e][1,4]diazepin-2-one(13, MYM-V-56)**

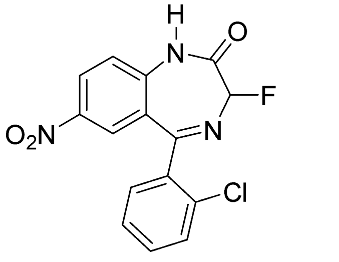


**13, MYM-V-56**

5-(2-chlorophenyl)-3-hydroxy-7-nitro-1,3-dihydro-2H-benzo[e][1,4]diazepin-2-one (**10**, 0.6 g, 1.83 mmol) was dissolved in anhydrous DCM (15 mL) and the mixture was cooled to -70°C using an acetone/dry ice bath. Then diethyl amino sulfur trifluoride (DAST, 0.8 mL, 6.1 mmol) was added dropwise to the reaction mixture over 5 min period while maintaining the temperature -70°C to -65°C. The mixture was then allowed to warm slowly to -20 to -10°C and stirred for 30 min at that temperature. The temperature of the reaction should be less than -10°C; otherwise N1H will be replaced by fluorine atom. The consumption of starting material was monitored by TLC (silica gel, 70% EtOAc-hexane; **$R_f$ of 13** =0.5; **$R_f$ of 10** =0.35). The reaction was quenched with water and the aq layer was extracted with chloroform (30 mL). The organic layer was washed with brine (2x10 mL) and dried ($Na_2SO_4$). The solvents were removed under reduced pressure and the residue was slurried in ethanol (8 mL). The mixture was stirred for 15 min at 80°C. Upon cooling to rt, the mixture was kept at -20°C freezer for 2 h. The residue was filtered, washed with cold ethanol (2x3 mL), and dried under vacuum at 45°C for 2h to afford pure MYM-V-56 (**13**) as white colored powder. **$^1$H NMR** (500 MHz, DMSO) δ 11.74 (s, 1H), 8.43 (dd, *J* = 9.0, 2.5 Hz, 1H), 7.77 (d, *J* = 2.4 Hz, 1H), 7.75 – 7.72 (m, 1H), 7.63 – 7.57 (m, 2H), 7.56 – 7.53 (m, 1H), 7.49 (d, *J* = 9.0 Hz, 1H), 6.00 (d, *J* = 55.4 Hz, 1H). **$^{13}$C NMR** (126 MHz, DMSO) δ 165.72 (d, *J* = 28.7 Hz), 162.64 (d, *J* = 22.4 Hz), 143.56 (s), 142.45 (s), 137.12 (s), 132.59 (s), 132.30 (s), 132.08 (s), 130.47 (s), 128.21 (s), 127.62 (s), 127.41 (s), 125.04 (s), 123.31 (s), 97.24 (d, *J* = 180.5 Hz). **HRMS** (LCMS-IT-TOF) Calc. for $C_{15}H_9N_3O_3FCl$ $(M + H)^+$ 334.0389, found 334.0393.

**5-(2-chlorophenyl)-3-hydroxy-1-methyl-7-nitro-1,3-dihydro-2H-benzo[e][1,4] diazepin-2-one (14, MYM-V-51)**

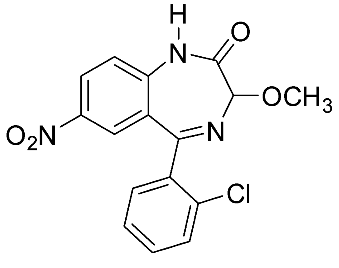


**14**

5-(2-chlorophenyl)-3-hydroxy-7-nitro-1,3-dihydro-2H-benzo[e][1,4]diazepin-2-one (**10**, 0.1 g, 0.3 mmol) was dissolved in anhydrous acetonitrile (3 mL) and 2 drops of concentrated $H_2SO_4$ was added. The mixture was stirred for 5 min and anhydrous methanol (10 mL) was added. The mixture was stirred for 10 min at 50-60°C. The all the solvents were evaporated under reduced pressure. Water (10 mL) and DCM (20 mL) were added and the mixture was allowed to stand to separate layers. The organic layer was separated, washed (brine, 2x 10 mL), and dried ($Na_2SO_4$). The solvents were removed under reduced pressure and the residue was purified by a flash chromatography (silica gel, 50% EtOAc-hexane) to afford **14** as white solid (67 mg, 65%). **$^1$H NMR** (500 MHz, $CDCl_3$) δ 8.71 (dd, *J* = 9.2, 2.5 Hz, 1H), 8.61 (d, *J* = 2.4 Hz, 1H), 8.42 (d, *J* = 9.2 Hz, 1H), 7.64 (d, *J* = 8.0 Hz, 1H), 7.62 – 7.56 (m, 1H), 7.54 (d, *J* = 4.2 Hz, 2H), 5.75 (s, 1H), 3.61 (s, 3H). **$^{13}$C NMR** (126 MHz, $CDCl_3$) δ 170.24 (s), 163.92 (s), 152.84 (s), 146.37 (s), 134.59 (s), 132.75 (s), 131.72 (s), 131.44 (s), 131.12 (s), 130.42 (s), 127.57 (s), 127.41 (s), 123.81 (s), 122.21 (s), 103.90 (s), 54.60 (s). **HRMS** (LCMS-IT-TOF) Calc. for $C_{16}H_{12}N_3O_4Cl$ $(M + H)^+$ 346.05891, found 346.0589.

**<u>1-(4-(2-chlorophenyl)-6-nitro-2-oxo-1,2-dihydroquinolin-3-yl)pyridin-1-ium chloride (15, MYM-V-48)</u>**

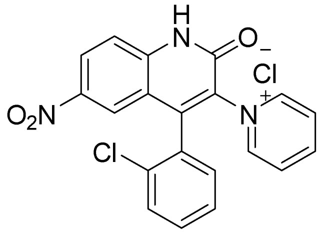


**15, MYM-V-48**

A round bottom flask was charged with 2-bromo-N-(4-nitro-2-(2-chlorobenzoyl) phenyl) acetamide (**4**, 1 g, 2.52 mmol), pyridine (10 mL) and the mixture was refluxed for 2 h at 115°C. The reaction mixture was then cooled to rt and hold for 1 h. The solid residue, which resulted, was filtered and washed with anhydrous DCM (2x10 mL). The residue was then dried under vacuum for 2 h to afford **14** as an off-white powder (1.02 g, 98%). **$^1$H NMR** (500 MHz, DMSO) δ 13.67 (s, 1H), 9.23 (d, *J* = 6.2 Hz, 1H), 9.15 (d, *J* = 6.2 Hz, 1H), 8.91 – 8.88 (m, 1H), 8.76 (tt, *J* = 7.9, 1.4 Hz, 1H), 8.61 (dd, *J* = 9.2, 2.5 Hz, 1H), 8.29 (dt, *J* = 12.5, 6.6 Hz, 2H), 7.82 (d, *J* = 9.2 Hz, 1H), 7.77 (d, *J* = 2.5 Hz, 1H), 7.65 (dd, *J* = 8.0, 1.1 Hz, 1H), 7.60 (td, *J* = 7.5, 1.8 Hz, 1H), 7.56 (dd, *J* = 7.9, 1.9 Hz, 1H), 7.52 (td, *J* = 7.5, 1.2 Hz, 1H). **$^{13}$C NMR** (126 MHz, DMSO) δ 169.62 (s), 162.37 (s), 144.05 (s), 142.16 (s), 137.93 (s), 132.27 (s), 132.07 (s), 131.96 (s), 130.32 (s), 128.05 (s), 127.48 (s), 127.10 (s), 124.87 (s), 122.68 (s), 83.59 (s), 56.49 (s), 19.03 (s). **HRMS** (LCMS-IT-TOF) Calc. for $C_{20}H_{12}N_3O_3Cl$ $(M + H)^+$ 378.0640, found 378.0645.

**<u>3-Amino-4-(2-chlorophenyl)-6-nitroquinolin-2(1H)-one (16, MYM-V-49)</u>**

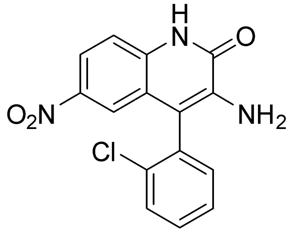


**16, MYM-V-49**

The pyridinium salt, 1-(4-(2-chlorophenyl)-6-nitro-2-oxo-1,2-dihydroquinolin-3-yl)pyridin-1-ium chloride (**15,** 1.5 g  3.62 mmol) was slurried in ethanol (10 mL) and hydrazine hydrate (0.34 mL, 7.24 mmol) was added to that slurry. The mixture was then refluxed for 3 h at 80°C. The mixture was then cooled to rt, diluted with water (30 mL) and DCM (40 mL). After separating the organic layer, the aq. layer was extracted with DCM (2x10 mL). The combined organic layer was washed with water (2x 10 mL), 10% aq. NaCl (20 mL), and dried ($Na_2SO_4$). The solvents were removed under reduced pressure and the residue was slurried in 10% EtOAc-hexane (10 mL). The mixture was stirred for 10 min at 50°C. Upon cooling to rt and holding for 1 h, the residue was filtered, washed with 10% EtOAc-hexane (2x5 mL), and dried under vacuum to afford pure **16** as a yellow colored powder (903 mg, 79%). **$^1$H NMR** (500 MHz, DMSO) δ 12.76 – 12.29 (m, 1H), 8.04 (dd, *J* = 8.9, 2.5 Hz, 1H), 7.75 – 7.72 (m, 1H), 7.60 – 7.56 (m, 2H), 7.45 (d, *J* = 8.9 Hz, 1H), 7.43 – 7.39 (m, 2H), 5.46 (s, 2H). **$^{13}$C NMR** (126 MHz, DMSO) δ 158.39 (s), 142.53 (s), 136.86 (s), 136.59 (s), 134.02 (s), 132.78 (s), 132.77 (s), 131.07 (s), 130.76 (s), 128.97 (s), 121.84 (s), 119.68 (s), 117.83 (s), 116.30 (s), 114.00 (s). **HRMS** (LCMS-IT-TOF) Calc. for $C_{15}H_{10}N_3O_3Cl$ $(M + H)^+$ 316.0483, found 316.0488.

**Scheme 3. Synthesis of C-7 modified analogs of meclonazepam**

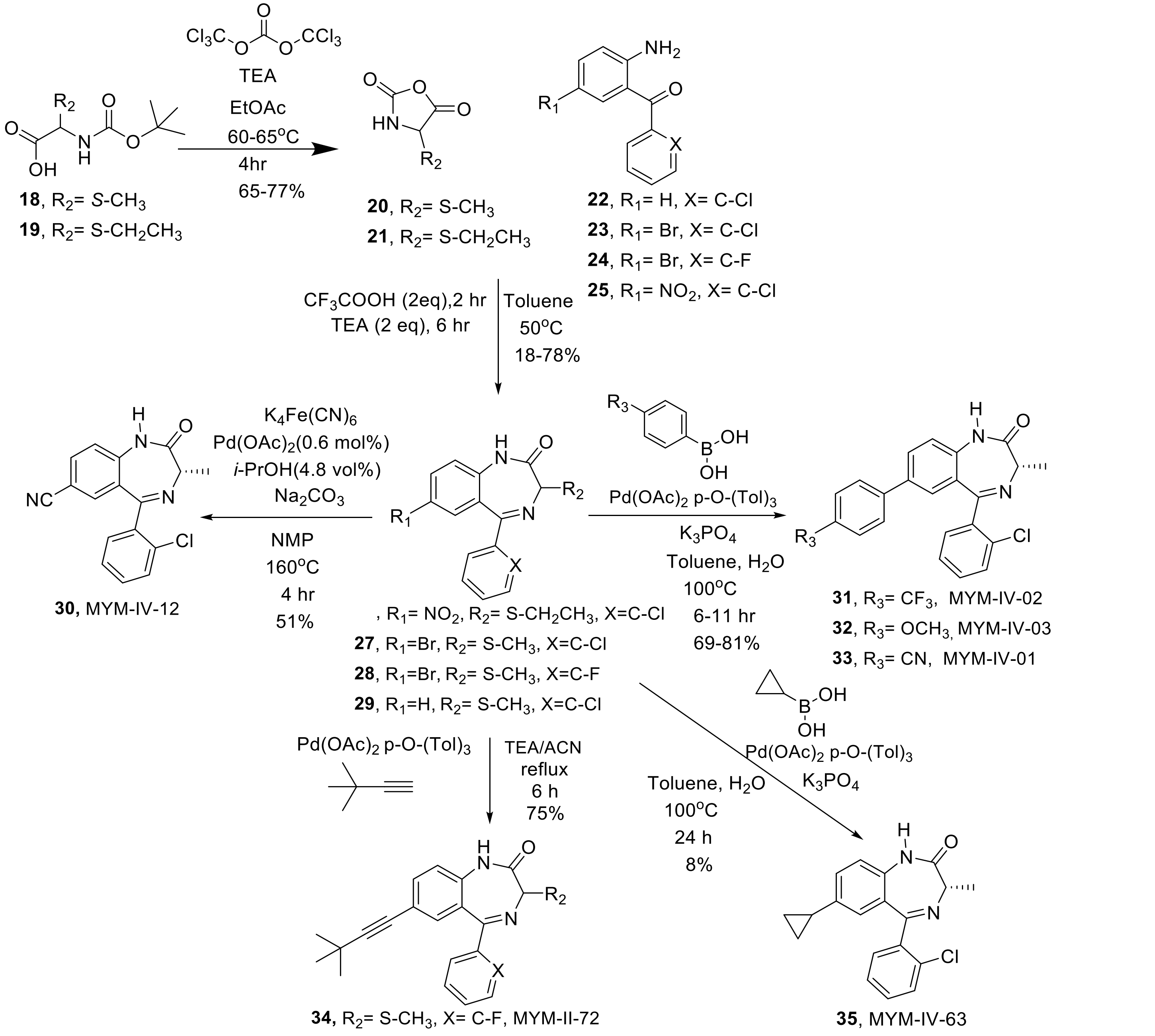

**(S)-4-methyloxazolidine-2,5-dione (20)**

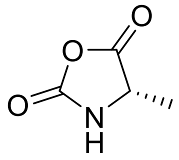

**20**

A three neck round bottom flask was charged with Boc-Ala-OH (50 g, 264.25 mmol) and anhydrous ethyl acetate (2000mL) under argon atmosphere and the mixture was stirred to make a solution. Triphosgene (31.35g, 105.7 mmol) was then added to that reaction mixture followed by dropwise addition of triethylamine (40.5 mL, 290.72 mmol) over a period of 60 min at 25 – 30 °C temperature. $CO_2$(g) and HCl(g) was evolved during the addition of triethylamine. The resulted reaction mixture was then allowed to stir additional 1 h at 25 – 30°C. After stirring 1hr at room temperature, the mixture was refluxed at 65 – 70°C for 2 h at which point the reaction was deemed complete as gas evolution ceased. A white slurry resulted at this point. The reaction mixture was then cooled to rt and hold for 3hr. The white TEA-HCl salt was removed by filtration and the residue was washed with ethyl acetate (2 x 100 mL). The pale yellow colored filtrates were then concentrated under reduced pressure to yield a semi-solid mass. The residue was then dissolved in anhydrous dichloromethane (200 mL). Then anhydrous hexane (200ml) was dropwise slowly over 1 hr with vigorous stirring. The mixture was then placed in a freezer at -20 °C overnight to maximize product precipitation. The hexane needs to be added very slowly, quick addition of hexane to the dichloromethane solution causes oily product which may not precipitate at all and hence difficult to collect the NCA product. The white crystalline solid was filtered and washed with anhydrous hexanes (150 mL x 2). The solid was dried under vacuum at room temperature for 2 h to afford the product as an off-white solid (20.8 g, 70%). **$^1$H NMR** (500 MHz, DMSO) δ 8.98 (s, 1H), 4.47 (q, *J* = 6.7 Hz, 1H), 1.34 (d, *J* = 6.6 Hz, 3H). **$^{13}$C NMR** (126 MHz, DMSO) δ 172.85 (s), 152.15 (s), 53.27 (s), 17.18 (s).

**(S)-5-(2-chlorophenyl)-3-ethyl-1,3-dihydro-2H-benzo[e][1,4]diazepin-2-one (26, MYM-V-19)**

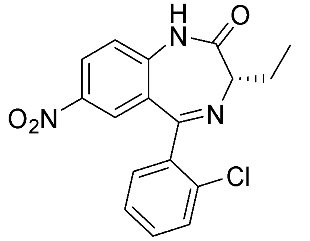

Chemical Formula: $C_{17}H_{14}ClN_3O_3$
Molecular Weight: 343.77

(2-Amino-5-nitrophenyl)(2-chlorophenyl)methanone ( 0.3 g, 1.0 mmol) was dissolved in anhydrous toluene ( 10 mL) and trifluoro acetic acid (0.2 mL, 2 mmol) was added. The mixture was heated to 50 °C. At that point, (*S*)-4-ethyloxazolidine-2,5-dione (NCA, 0.24 g, 2.5 mmol) was added one. The mixture was then stirred for an hour at 50°C at which point the starting material was deemed to consume (TLC, silica gel, 50% EtOAc-hexane) to form the TFA-salt of amide intermediate. Then, triethylamine (0.4 mL, 2.8 mmol) added dropwise and a white fume was observed during the addition. Upon completion of addition, the mixture was stirred for additional one hour. The consumption of starting material was confirmed by TLC (silica gel, 50% EtOAc-hexane). The solvents were evaporated and the residue was dissolved in dichlormethane. Water (20 mL) was added to dilute the mixture. The layers were separated and the aq. layer was extracted with DCM (2x10 mL). The organic layer was washed with water (2x10 mL) and dried ($Na_2SO_4$). The solvents were removed under reduced pressure and the residue was purified by neutral alumina column chromatograpy to afford pure **23** as a white colored powder (67 mg, 18%). **$^1$H NMR** (500 MHz, $CDCl_3$) δ 9.60 (s, 1H), 8.35 (dd, *J* = 8.9, 2.5 Hz, 1H), 8.02 (d, *J* = 2.5 Hz, 1H), 7.59 (dd, *J* = 5.6, 2.4 Hz, 1H), 7.46 – 7.43 (m, 2H), 7.41 – 7.38 (m, 1H), 7.33 (d, *J* = 8.9 Hz, 1H), 3.55 (t, *J* = 7.1 Hz, 1H), 2.37 – 2.25 (m, 2H), 1.14 (td, *J* = 7.4, 4.7 Hz, 3H). **$^{13}$C NMR** (126 MHz, $CDCl_3$) δ 171.18 (s), 167.49 (s), 143.10 (s), 142.59 (s), 137.68 (s), 133.18 (s), 131.52 (s), 131.37 (s), 130.37 (s), 128.32 (s), 127.33 (s), 126.51 (s), 125.65 (s), 121.73 (s), 65.15 (s), 24.20 (s), 10.61 (s). **HRMS** (LCMS-IT-TOF) Calc. for $C_{17}H_{14}N_3O_3Cl$ $(M + H)^+$ 344.0796; found 344.0792.

**(S)-7-bromo-5-(2-chlorophenyl)-3-methyl-1,3-dihydro-2H-benzo[e][1,4]diazepin-2-one (27)**

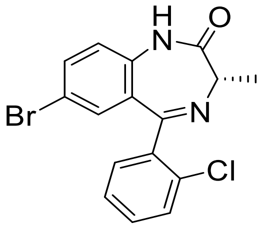


**27**

Anhydrous trifluoroacetic acid (14.70 mL, 192 mmol) was added to a stirred solution of commercially available 2-amino-5-bromo-2'-chlorobenzophenone (30 g, 96 mmol) in anhydrous toluene (500 mL) under argon atmosphere. The mixture was allowed to stir at rt for 30 min to dissolve the starting material which resulted a yellow-colored solution. The N-carboxy-L-alanine anhydride (**18**, 13.34 g, 115 mmol) was then added to the reaction mixture at once. The reaction mixture was then stirred for 1 h at 50 – 55°C for consumption of starting material. TLC (silica gel and 30% ethyl acetate / hexanes) confirmed the formation of TFA salt intermediate. It was observed that less than 5% 2-amino-5-bromo-2'-chlorobenzophenone remained after 1 hr on TLC. Then anhydrous triethylamine (26.9 mL, 192 mmol) was added dropwise slowly to the reaction mixture over a 20 min period at 50 – 55°C temperature. Once the addition was done, the reaction mixture was stirred for 2h at 50 – 55°C. TLC (silica gel; 50% ethyl acetate / hexanes) confirmed the disappearance of TFA salt intermediate. The reaction mixture was then cooled to rt, the solvents were concentrated under reduced pressure, and the residue was diluted with ethyl acetate (500 mL) and water (500 mL). The biphasic mixture was then stirred for 5 min, allowed to stand for 10 min, and transferred to separatory funnel. The layers were separated and the aquous layer was extracted with EtOAc (2x150). The combined organic layers were washed with 10% aq sodium chloride solution (2x300mL) and dried ($Na_2SO_4$). The solvents were removed under reduced pressure and the residue was slurried in 10% ethyl acetate / hexane (200 mL). The slurry was then stirred at 60 – 65°C for 30 min. The mixture was cooled to rt and kept for 2 h at rt to precipitate the product. The solid was then filtered, washed with 10% ethyl acetate / heptane (50 mL x 2) collected by filtration and washed and then heptane (30 mL x 2), and dried under vacuum at 35 – 40°C to afford the product as an off-white solid of **25**(22.93 g, 72.0%). **$^1$H NMR** (500 MHz, $CDCl_3$) δ 9.74 (s, 1H), 7.58 (dd, J = 8.6, 2.2 Hz, 1H), 7.54 – 7.48 (m, 1H), 7.41 – 7.33 (m, 3H), 7.22 (d, J = 2.2 Hz, 1H), 7.11 (d, J = 8.6 Hz, 1H), 3.83 (q, J = 6.5 Hz, 1H), 1.78 (d, J = 6.5 Hz, 3H); **$^{13}$C NMR** (126 MHz, $CDCl_3$) δ 172.28 (s), 167.31 (s), 138.21 (s), 136.90 (s), 134.75 (s), 133.33 (s), 131.89 (s), 131.14 (s), 130.92 (s), 130.19 (s), 129.86 (s), 127.02 (s), 122.81 (s), 116.49 (s), 58.77 (s), 16.88 (s); **HRMS (ESI/IT-TOF):** m/z [M + H]+ calcd for $C_{16}H_{13}BrClN_2O$: 362.9894; found: 362.9926.

**(S)-7-bromo-5-(2-fluorophenyl)-3-methyl-1,3-dihydro-2H-benzo[e][1,4]diazepin-2-one (26)**

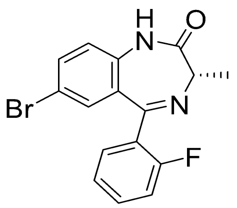

**26**

Compound **26 (**8.31 g, 71%) was synthesized by NCA route following the procedure described for **25** from the commercially available benzophenone **22** ( 10 g, 33.9 mmol**).** $^{1}$**H NMR** (500 MHz, $CDCl_3$) δ 9.62 (s, 1H), 7.59 (ddd, J = 7.1, 5.0, 2.0 Hz, 2H), 7.48 – 7.43 (m, 1H), 7.36 (d, J = 2.1 Hz, 1H), 7.25 (t, J = 7.5 Hz, 1H), 7.12 (d, J = 8.6 Hz, 1H), 7.09 – 7.05 (m, 1H), 3.80 (q, J = 6.5 Hz, 1H), 1.79 (d, J = 6.5 Hz, 3H**).** $^{13}$**C NMR** (126 MHz, $CDCl_3$) δ 172.24 (s), 164.52 (s), 160.43 (d, J = 252.1 Hz), 136.47 (s), 134.74 (s), 132.18 (d, J = 8.3 Hz), 132.08 (s), 131.56 (d, J = 1.6 Hz), 130.07 (s), 127.06 (d, J = 12.2 Hz), 124.42 (d, J = 3.6 Hz), 122.90 (s), 116.50 (s), 116.29 (d, J = 21.5 Hz), 58.83 (s), 16.91 (s). **HRMS** (ESI/IT-TOF) m/z: $[M + H]^+$ Calcd for $C_{16}H_{13}BrFN_2O$ 347.0189; found 347.0190

**((S)-5-(2-chlorophenyl)-3-methyl-1,3-dihydro-2H-benzo[e][1,4]diazepin-2-one (29, MYM-V-03)**

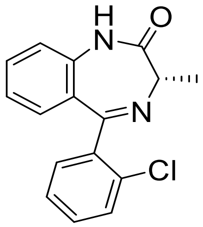

**29**

Anhydrous trifluoroacetic acid was added (0.5 mL, 8.6 mmol) to the stirred solution (2-aminophenyl)(2-chlorophenyl)methanone (1 g, 4.3 mmol) and the reaction mixture was heated to 50°C. N-carboxy alanine anhydride (NCA, 0.65 g, 5.6 mmol) was added one portions .The mixture was stirred for an hour at 50°C until the starting material was fully consumed, as confirmed by TLC analysis. Following this, triethylamine (1.2 mL, 8.6 mmol) was added dropwise, resulting in the formation of white fumes. After one more hour of stirring, TLC analysis again confirmed the complete consumption of the starting material. The solvents were then evaporated under reduced pressure, and the residue was dissolved in dichloromethane. The addition of water (20 mL) facilitated the separation of the layers, and the aqueous layer was extracted with dichloromethane (2x10 mL). The organic layer was washed with water (2x10 mL) and dried using sodium sulfate (Na2SO4). After removing the solvents under reduced pressure, the remaining residue was purified by washing with a 10% ethyl acetate (EtOAc) and hexane mixture followed by drying under vacuum at 50°C to afford compound **27** (0.6 g, 78%) as a white-colored powder. $^{1}$**H NMR** (500 MHz, $CDCl_3$) δ 8.98 (s, 1H), 7.53 (s, 1H), 7.50 (td, 1H), 7.37 (d, J = 3.0 Hz, 3H), 7.17 (t, J = 7.6 Hz, 1H), 7.12 (dd, 2H), 3.86 (q, J = 6.5 Hz, 1H), 1.79 (d, J = 6.5 Hz, 3H). $^{13}$**C NMR** (126 MHz, CDCl3) δ 168.52 (s), 138.99 (s), 137.64 (s), 133.37 (s), 131.77 (s), 131.13 (s), 130.54 (s), 130.02 (s), 129.68 (s), 128.32 (s), 128.19 (s), 126.86 (s), 123.76 (s), 120.90 (s), 58.63 (s), 16.96 (s). **HRMS (ESI/ITTOF) m/z: [M + H]+** Calcd for $C_{16}H_{13}N_2OCl$ 285.0789; found 285.0767.

**(S)-5-(2-chlorophenyl)-3-methyl-2-oxo-2,3-dihydro-1H-benzo[e][1,4]diazepine-7-carbonitrile (30, MYM-IV-12)**

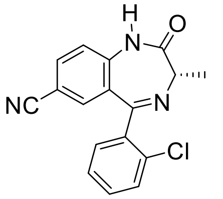

**30**

A round bottom flask was charged with (S)-7-bromo-5-(2-chlorophenyl)-3-methyl-1,3-dihydro-2H-benzo[e][1,4]diazepin-2-one (**25**) (1 g, 2.75 mmol), *i*-PrOH (0.5 ml, 4.8 vol%), $Na_2CO_3$ (0.32 g, 3.02 mmol), Pd(OAc)2 (0.0037 g, .0167 mmol), and NMP (10 mL) under argon atmosphere. The mixture was heated to 140°C. At that point, $K_4Fe(CN)_6.3H_2O$ (0.46 g, 1.1 mmol) was added to the mixture at once. The mixture was heated for 4 h under sealed condition while maintaining temperature 140-150°C. The completion of reaction was monitored by TLC (silica gel, 40% EtOAc-hexane). Upon cooling, the mixture was diluted with water (10 mL) and ethyl acetate (10 mL). The layers were separated and the aq. layer was extracted with ethyl acetate (2x10 mL). The combined organic layer was washed with water (5x 10 mL), brine (2x10 mL), and dried ($Na_2SO_4$). The solvents were evaporated, and the residue was purified by column chromatography (silica gel, 20% EtOAc-hexane to 40% EtOAc-hexane). The appropriate fractions were pooled and the solvents were removed under reduced pressure. The residue was dried under vacuum for 1 h to afford pure **28** as white powder (450 mg, 53%). $\mathbf{R_f}$ = 0.3 (silica TLC, 40% EtOAc-hexane, $R_f$ of **47**=0.5). $^1$**H NMR** (300 MHz, $CDCl_3$) δ 9.84 (s, 1H), 7.71 (dd, *J* = 8.5, 1.7 Hz, 1H), 7.57 – 7.53 (m, 1H), 7.44 – 7.41 (m, 1H), 7.39 (dd, *J* = 7.0, 2.3 Hz, 3H), 7.35 (d, *J* = 8.5 Hz, 1H), 3.80 (q, *J* = 6.5 Hz, 1H), 1.76 (d, *J* = 6.5 Hz, 3H). **HRMS (ESI/IT-TOF)** m/z: $[M + H]^+$ Calcd for $C_{17}H_{12}N_3OCl$ 310.07417; found 310.07403.

**((S)-5-(2-chlorophenyl)-3-methyl-7-(4-(trifluoromethyl)phenyl)-1,3-dihydro-2H-benzo[e][1,4]diazepin-2-one (31, MYM-IV-02)**

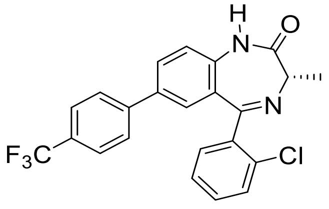

**31**

To a mixture of $Pd(OAc)_2$ (0.044 g, 0.189 mmol), tri-(O-tolyl) phosphine (0.119 g, 0. 378 mmol) in toluene (10 mL), the benzodiazepine, (S)-7-bromo-5-(2-chlorophenyl)-3-methyl-1,3-dihydro-2H-benzo[e][1,4]diazepin-2-one **25** (0.1 g, 2.7 mmol), 4-trifluoromethyl phenyl boronic acid ( 2 g, 10 mmol), tri-basic potassium phosphate (2.5 g, 12.1 mmol), water (0.4 mL, 12.1 mmol) were added sequentially under argon. The reaction mixture was then stirred at 100°C for 6 h. LCMS 2020 (single quadrupole mass analyzer), and TLC (silica gel, 50% EtOAc-hexane) confirmed the consumption of starting material. The reaction mixture was cooled and opened to air once all the starting material consumed. The reaction mixture was passed through a pad of celite bead to remove any palladium salts. The filtrate was diluted with water (20 mL) and ethyl acetate (20 mL). The biphasic mixture, which resulted, was allowed to stand to separate. The organic layer was collected, and the aqueous layer was extracted (2x10 mL). The combined organic layer was washed with 10% aq. NaCl (3x10 mL) and dried ($Na_2SO_4$). The solvents were removed under reduced pressure. The orange-colored residue, which resulted, was purified a flash chromatography (silica gel 100g, 40% EtOAc-hexane). The desired fractions were pooled, and the solvents were removed. The solid residue was dried under vacuum for 2 h to afford a yellow-colored powder of **29** (0.94 g, 80%). **$^1$H NMR** (500 MHz, $CDCl_3$) δ 9.31 (s, 1H), 7.84 (s, 1H), 7.67 (d, *J* = 8.2 Hz, 2H), 7.57 (dd, *J* = 5.6, 3.4 Hz, 2H), 7.43 (dd, *J* = 16.5, 7.9 Hz, 2H), 7.33 (dd, *J* = 5.4, 3.2 Hz, 1H), 7.11 (s, 2H), 6.88 (d, *J* = 7.7 Hz, 1H), 3.80 (q, *J* = 6.3 Hz, 1H), 1.79 (d, *J* = 5.0 Hz, 2H); **$^{13}$C NMR** (126 MHz, $CDCl_3$) δ 172.14 (s), 170.09 (s), 144.62 (s), 142.95 (s), 140.82 (s), 139.03 (s), 137.63 (s), 134.72 (s), 130.90 (s), 130.18 (d, *J* = 9.4 Hz), 129.87 (s), 129.12 (s), 128.39 (s), 128.13 (s), 127.06 (s), 125.94 (q, *J* = 3.7 Hz), 125.18 (d, *J* = 5.4 Hz), 124.54 (q, *J* = 3.5 Hz), 123.02 (d, *J* = 5.4 Hz), 58.82 (s), 16.77 (s); **HRMS** (ESI/IT-TOF) m/z: $[M + H]^+$ Calcd for $C_{23}H_{16}N_2OF_3Cl$ 429.0976; found 429.0975.

**(S)-5-(2-chlorophenyl)-7-(4-methoxyphenyl)-3-methyl-1,3-dihydro-2H-benzo[e][1,4] diazepin-2-one (32, MYM-IV-30)**

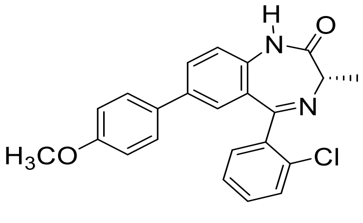


**32**

$Pd(OAc)_2$ (0.025 g, 0.115 mmol), tri-(O-tolyl) phosphine (0.07 g, 0. 23 mmol) were dissolved in toluene (10 mL) and the mixture was stirred for 10 min under argon atmosphere to generate the $Pd(OAc)_2$-p-O-$(tol)_3$-phosphine catalyst *in-situ*. Then benzodiazepine, (S)-7-bromo-5-(2-chlorophenyl)-3-methyl-1,3-dihydro-2H-benzo[e][1,4]diazepin-2-one **25** (0.6 g, 1.65 mmol), 4-methoxy phenyl boronic acid ( 0.51 g, 3.3 mmol), tri-basic potassium phosphate (1.57 g, 7.42 mmol), water (0.2 mL, 7.42 mmol) were added sequentially to the previous reaction mixture under argon. The reaction mixture was then stirred at 100°C for 6 h. LCMS 2020 (single quadrupole mass analyzer), and TLC (silica gel, 50% EtOAc-hexane) confirmed the consumption of starting material. The reaction mixture was cooled and opened to air once all the starting material consumed. The reaction mixture was passed through a pad of celite bead to remove any palladium salts. The filtrate was diluted with water (20 mL) and ethyl acetate (20 mL). The biphasic mixture, which resulted, was allowed to stand to separate. The organic layer was collected, and the aqueous layer was extracted (2x10 mL). The combined organic layer was washed with 10% aq. NaCl (3x10 mL) and dried ($Na_2SO_4$). The solvents were removed under reduced pressure. The orange-colored residue, which resulted, was purified a flash chromatography (silica gel 100 g, 40% EtOAc-hexane). The desired fractions were pooled, and the solvents were removed under reduced pressure. The solid residue was dried under vacuum for 2 h to afford a yellow-colored powder of **30** (0.55 g, 79%). **$^1$H NMR** (500 MHz, $CDCl_3$) δ 9.23 (s, 1H), 7.68 (dd, *J* = 8.4, 2.1 Hz, 1H), 7.57 (dd, *J* = 8.3, 3.6 Hz, 1H), 7.40 – 7.36 (m, 5H), 7.25 (d, *J* = 2.0 Hz, 1H), 7.23 (d, *J* = 4.1 Hz, 1H), 6.96 – 6.92 (m, 2H), 3.93 (q, *J* = 6.5 Hz, 1H), 3.83 (s, 3H), 1.82 (d, *J* = 6.5 Hz, 3H). **$^{13}$C NMR** (126 MHz, $CDCl_3$) δ 172.22 (s), 168.55 (s), 159.44 (s), 138.89 (s), 136.50 (s), 136.39 (s), 133.36 (s), 131.94 (s), 131.16 (s), 130.59 (s), 130.14 (s), 130.11 (s), 128.66 (s), 128.04 (s), 127.38 (s), 126.92 (s), 121.40 (s), 114.35 (s), 58.75 (s), 55.38 (s), 16.99 (s). **HRMS** (ESI/IT-TOF) m/z: $[M + H]^+$ Calcd for $C_{23}H_{19}N_2O_2Cl$ 391.1207; found 391.1206.

**(S)-4-(5-(2-chlorophenyl)-3-methyl-2-oxo-2,3-dihydro-1H-benzo[e][1,4]diazepin-7-yl)benzonitrile (33, MYM-IV-01)**

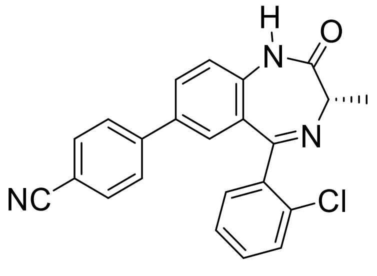

**33**

$Pd(OAc)_2$ (0.025 g, 0.115 mmol), tri-(O-tolyl) phosphine (0.07 g, 0. 23 mmol) were dissolved in toluene (10 mL) and the mixture was stirred for 10 min under argon atmosphere to generate the $Pd(OAc)_2$-p-O-$(tol)_3$-phosphine catalyst *in-situ*. Then the bwnzodiazepine, (S)-7-bromo-5-(2-chlorophenyl)-3-methyl-1,3-dihydro-2H-benzo[e][1,4]diazepin-2-one **25** (0.6 g, 1.65 mmol), 4-cyano phenyl boronic acid ( 0.5 g, 3.3 mmol), tri-basic potassium phosphate (1.57 g, 7.42 mmol), water (0.2 mL, 7.42 mmol) were added sequentially to the previous reaction mixture under argon. The reaction mixture was then stirred at 100°C for 6 h. LCMS 2020 (single quadrupole mass analyzer), and TLC (silica gel, 50% EtOAc-hexane) confirmed the consumption of starting material. The reaction mixture was cooled and opened to air once all the starting material consumed. The reaction mixture was passed through a pad of celite bead to remove any palladium salts. The filtrate was diluted with water (20 mL) and ethyl acetate (20 mL). The biphasic mixture, which resulted, was allowed to stand to separate. The organic layer was collected and the aqueous layer was extracted (2x10 mL). The combined organic layer was washed with 10% aq. NaCl (3x10 mL) and dried ($Na_2SO_4$). The solvents were removed under reduced pressure. The orange colored residue, which resulted, was purified a flash chromatography (silica gel 100g, 40% EtOAc-hexane). The desired fraction were pooled and the solvents were removed. The solid residue was dried under vacuum for 2 h to afford an yellow colored powder of **31** (0.42 g, 69%). **$^1$H NMR** (500 MHz, $CDCl_3$) δ 9.27 (s, 1H), 7.74 – 7.67 (m, 3H), 7.59 (s, 1H), 7.53 (d, *J* = 8.1 Hz, 2H), 7.39 (dd, *J* = 8.4, 5.0 Hz, 3H), 7.30 (dd, *J* = 12.7, 8.9 Hz, 2H), 3.92 (q, *J* = 6.3 Hz, 1H), 1.82 (d, *J* = 6.4 Hz, 3H). **$^{13}$C NMR** (126 MHz, $CDCl_3$) δ 172.11 (s), 168.16 (s), 143.79 (s), 138.56 (s), 138.01 (s), 134.69 (s), 133.31 (s), 132.73 (s), 131.24 (s), 130.89 (s), 130.44 (s), 130.19 (s), 128.86 (s), 128.66 (s), 128.20 (s), 127.56 (s), 127.36 (s), 127.08 (s), 121.82 (s), 118.66 (s), 111.33 (s), 58.87 (s), 16.95 (s). **HRMS** (ESI/IT-TOF) m/z: [M + H] Calcd for $C_{23}H_{16}N_3OCl$ 386.10547; found 386.10550

**(S)-7-(3,3-dimethylbut-1-yn-1-yl)-5-(2-fluorophenyl)-3-methyl-1,3-dihydro-2H-benzo[e][1,4]diazepin-2-one (34, MYM-II-72)**

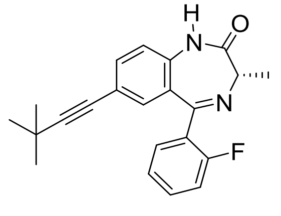

**32**

Tri-(O-tolyl) phosphine (0.15 g, 0.518 mmol), $Pd(OAc)_2$ (0.058 g, 0.259 mmol) were dissolved in ACN (5 mL) and the mixture was stirred for 10 min under argon atmosphere to generate the Pd catalyst *in-situ*. Then, the benzodiazepine **26** (1.5 g, 4.32 mmol), triethylamine (1.5 mL, 12.9 mmol), t-butyl acetylene (0.6 mL, 4.7 mmol) and additional acetonitrile (10 mL) were added sequentially to the previous reaction mixture under argon. The reaction mixture was then refluxed for 4 h and the consumption of starting material was confirmed by TLC (silica gel; 60% ethyl acetate-hexane). The mixture was then cooled to rt and silica gel (5g) was added to the flask and the mixture was stirred for 15 min. The contents were filtered through a pad of celite and the residue was washed with DCM (2x50 mL). The solvents were removed under reduced pressure. DCM (20 mL) and water (10 mL) was added to the resulting black oily mass to dissolve the contents. The mixture was stirred for 5 minutes and allowed to stand to separate layers for 5 min. The layers were separated, and the aqueous layer was extracted with DCM (2x10 mL). The combined organic layers were washed with 10% aq. NaCl (2x10 mL) and dried ($Na_2SO_4$). The DCM was removed under reduced pressure. The residue was purified by silica gel (50g) flash chromatography using 40% EtOAc-hexane. The desired fractions were collected, and the solvents were removed under reduced pressure. The yellow colored solid, which resulted was then dried under vacuum for 1hr to afford pure **32** (1.13 g, 75%) as yellow powder. $R_f$ = 0.6 (silica gel, 60% EtOAc-hexane). **$^1$H NMR** (500 MHz, $CDCl_3$) δ 9.41 (s, 1H), 7.60 (td, *J* = 7.5, 1.7 Hz, 1H), 7.49 (dd, *J* = 8.4, 1.9 Hz, 1H), 7.47 – 7.42 (m, 1H), 7.26 (dd, *J* = 7.5, 1.0 Hz, 1H), 7.24 (d, *J* = 1.5 Hz, 1H), 7.11 (d, *J* = 8.5 Hz, 1H), 7.08 – 7.04 (m, 1H), 3.77 (q, *J* = 6.5 Hz, 1H), 1.78 (d, *J* = 6.5 Hz, 3H), 1.28 (s, 9H); **$^{13}$C NMR** (126 MHz, $CDCl_3$) δ 172.37 (s), 165.26 (s), 160.52 (d, *J* = 252.0 Hz), 136.37 (s), 134.87 (s), 132.26 (s), 131.83 (d, *J* = 8.3 Hz), 131.60 (d, *J* = 2.3 Hz), 128.45 (s), 127.62 (d, *J* = 12.5 Hz), 124.30 (d, *J* = 3.5 Hz), 121.11 (s), 119.79 (s), 116.26 (d, *J* = 21.5 Hz), 99.14 (s), 77.72 (s), 58.83 (s), 30.90 (s), 27.92 (s), 16.97 (s); **HRMS (ESI/IT-TOF)** m/z: $[M + H]^+$ Calcd for $C_{23}H_{23}N_2OF$ 363.1867; found 363.1858.

***(S)*-5-(2-chlorophenyl)-7-cyclopropyl-3-methyl-1,3-dihydro-2H-benzo[e][1,4]diazepin-2-one (35)**

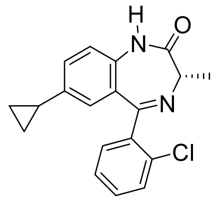

**35**

$Pd(OAc)_2$ (0.013 g, 0.075 mmol), tri-(O-tolyl) phosphine (0.04 g, 0. 12 mmol) were dissolved in toluene (10 mL) and the mixture was stirred for 10 min under argon atmosphere to generate the $Pd(OAc)_2$-p-O-$(tol)_3$-phosphine catalyst *in-situ*. Then the benzodiazepine, (S)-7-bromo-5-(2-chlorophenyl)-3-methyl-1,3-dihydro-2H-benzo[e][1,4]diazepin-2-one **26** (0.3 g, 0.83 mmol), 4-cyclopropyl boronic acid ( 0.2 g, 1.6 mmol), tri-basic potassium phosphate (0.75 g, 3.8 mmol), water (0.1 mL, 3.8 mmol) were added sequentially to the previous reaction mixture under argon. The reaction mixture was then stirred at 100°C for 6 h. LCMS 2020 (single quadrupole mass analyzer), and TLC (silica gel, 50% EtOAc-hexane) confirmed the consumption of starting material. The reaction mixture was cooled and opened to air once all the starting material consumed. The reaction mixture was passed through a pad of celite bead to remove any palladium salts. The filtrate was diluted with water (20 mL) and ethyl acetate (20 mL). The biphasic mixture, which resulted, was allowed to stand to separate. The organic layer was collected and the aqueous layer was extracted (2x10 mL). The combined organic layer was washed with 10% aq. NaCl (3x10 mL) and dried ($Na_2SO_4$). The solvents were removed under reduced pressure. The orange colored residue, which resulted, was purified by chromatography (longer bed of alumina 100 g, 2% EtOAc-hexane to 30% EtOAc-hexane). The desired fraction were pooled and the solvents were removed. The solid residue was dried under vacuum for 2 h to afford an yellow colored powder of **35** (0.015 g, 5.7%). **$^1$H NMR** (500 MHz, $CDCl_3$) δ 9.26 (s, 1H), 7.49 (t, *J* = 11.4 Hz, 1H), 7.42 – 7.35 (m, 2H), 7.24 (dt, *J* = 11.4, 2.9 Hz, 1H), 7.15 – 7.10 (m, 1H), 7.08 (dd, *J* = 8.3, 2.1 Hz, 1H), 6.84 (dd, *J* = 23.9, 1.5 Hz, 1H), 3.85 – 3.80 (m, 1H), 1.77 (d, *J* = 9.7 Hz, 3H), 1.28 (t, *J* = 7.1 Hz, 1H), 0.89 (dd, *J* = 17.6, 10.7 Hz, 3H), 0.55 – 0.48 (m, 2H). **$^{13}$C NMR** (126 MHz, $CDCl_3$) δ 170.91 (s), 168.43 (s), 141.58 (s), 140.80 (s), 138.97 (s), 133.33 (s), 131.13 (s), 130.43 (s), 129.99 (s), 129.56 (s), 129.37 (s), 128.80 (s), 128.57 (s), 126.80 (s), 60.41 (s), 16.95 (s), 14.85 (s), 9.19 (s), 7.16 (s).

**Chromatograms for separation of MYM-III-10 and MYM-V-56 enantiomers**

SHIMADZU LabSolutions **Analysis Report**

**<Sample Information>**

| | | | |
|---|---|---|---|
| Sample Name | : MYM-III-10 0.5mL | | |
| Sample ID | : MYM-III-10 0.5mL | | |
| Data Filename | : MYM-III-10 0.5mL.lcd | | |
| Method Filename | : Chiral separation MYM-III-10 by Prep LC.lcm | | |
| Batch Filename | : | | |
| Vial # | : 2 | Sample Type | : Unknown |
| Injection Volume | : 500 uL | | |
| Date Acquired | : 8/30/2021 5:14:45 PM | Acquired by | : Md Yeunus Mian |
| Date Processed | : 8/30/2021 6:04:53 PM | Processed by | : Md Yeunus Mian |

**<Chromatogram>**

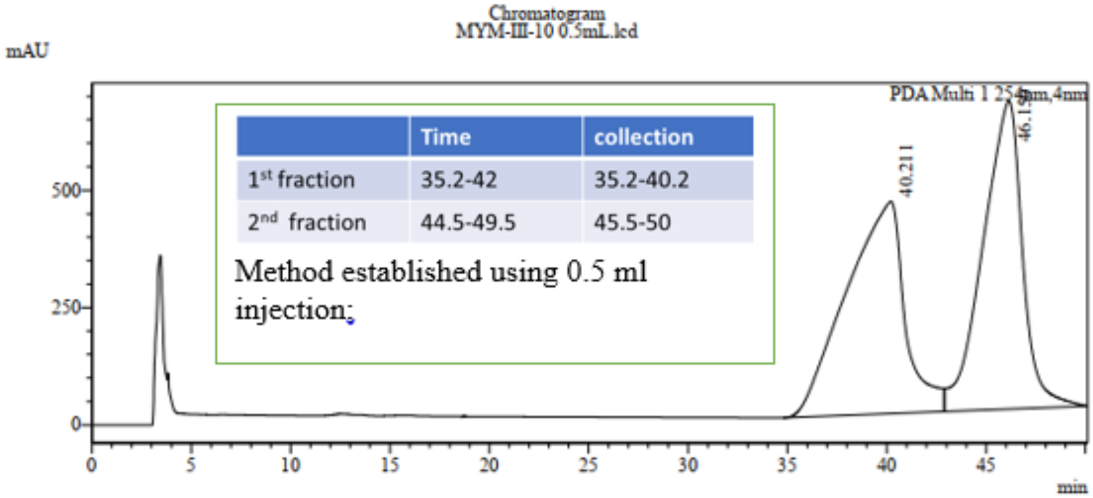


| | Time | collection |
|---|---|---|
| 1st fraction | 35.2-42 | 35.2-40.2 |
| 2nd fraction | 44.5-49.5 | 45.5-50 |

**<Peak Table>**

MYM-III-10 0.5mL.lcd

PDA Ch1 254nm

| Peak# | Ret. Time | Area% |
|---|---|---|
| 1 | 40.211 | 50.697 |
| 2 | 46.150 | 49.303 |
| Total | | 100.000 |

**Separation of MYM-III-10 enantiomers by preparative HPLC.** Reflect I cellulose C 5 µm, 25 cm x 21.1 mm, mobile phase 10 % EtOH-hexane. Flow rate 21.1 mL/min

SHIMADZU LabSolutions

### Analysis Report

#### <Sample Information>

| | | | |
|---|---|---|---|
| Sample Name | : MYM-III-10-1 | | |
| Sample ID | : MYM-III-10-1 | | |
| Data Filename | : MYM-III-10-1_8-30-21-EtOH-Hex=12-88_52 min.lcm_8-31_001.lcd | | |
| Method Filename | : 8-30-21-EtOH-Hex=12-88_52 min.lcm | | |
| Batch Filename | : 8-31.lcb | | |
| Vial # | : Vial 1 | Sample Type | : Unknown |
| Injection Volume | : 20 uL | Level | : 1 |
| Date Acquired | : 8/31/2021 3:39:21 PM | Acquired by | : Md Yeunus Mian |
| Date Processed | : 8/31/2021 4:48:01 PM | Processed by | : Md Yeunus Mian |

#### <Chromatogram>

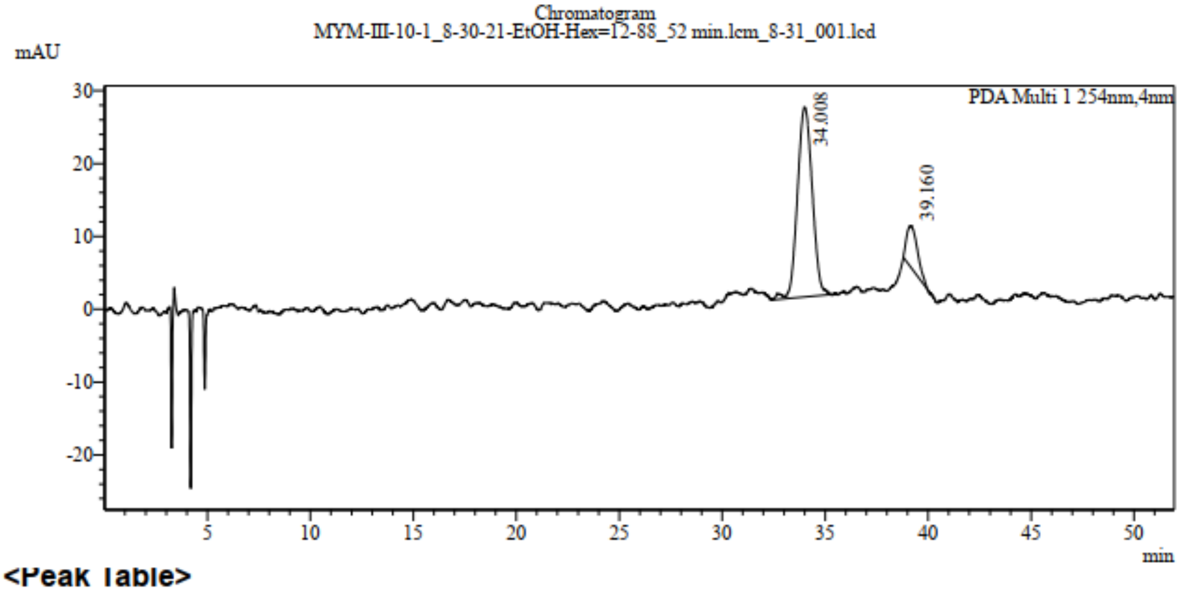


#### <Peak Table>

Peak Table MYM-III-10-1_8-30-21-EtOH-Hex=12-88_52 min.lcm_8-31_001.lcd

(3D)DAD Ch1 254nm

| Peak# | Ret. Time | Area% |
|---|---|---|
| 1 | 34.008 | 85.964 |
| 2 | 39.160 | 14.036 |
| Total | | 100.000 |

**HPLC chromatogram of S-MYM-III-10 at 20 minutes.** 10% S was converted to R after 20 minutes in solution. Normal phase chiral HPLC, Chiral pak IBN3 column (dimensions 4.6 mm x 150 mm, particle size 3 μm, mobile phase 10% EtOH-hexane).

#### <Sample Information>

| | | | |
|---|---|---|---|
| Sample Name | : rac-10-2 | | |
| Sample ID | : rac-10-2 | | |
| Data Filename | : rac-10-2_8-17-21-EtOH-Hex=12-88_80 min.lcm_8-17-2_001.lcd | | |
| Method Filename | : 8-17-21-EtOH-Hex=12-88_80 min.lcm | | |
| Batch Filename | : 8-17-2.lcb | | |
| Vial # | : Vial 2 | Sample Type | : Unknown |
| Injection Volume | : 20 uL | Level | : 1 |
| Date Acquired | : 8/17/2021 7:29:43 PM | Acquired by | : Md Yeunus Mian |
| Date Processed | : 8/17/2021 8:50:49 PM | Processed by | : Md Yeunus Mian |

#### <Chromatogram>

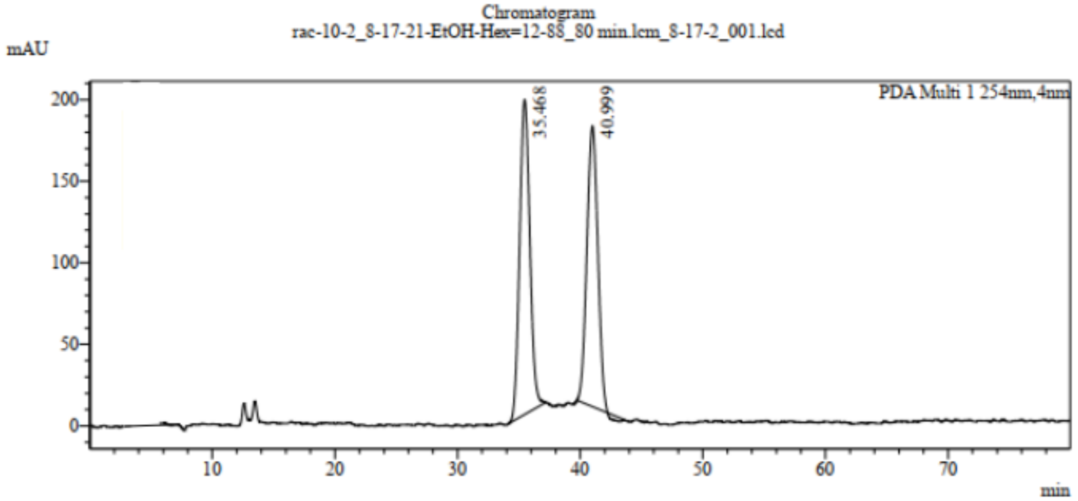


#### <Peak Table>

rac-10-2_8-17-21-EtOH-Hex=12-88_80 min.lcm_8-17-2_001.lcd

(3D)DAD Ch1 254nm

| Peak# | Ret. Time | Area% |
|---|---|---|
| 1 | 35.468 | 51.580 |
| 2 | 40.999 | 48.420 |
| Total | | 100.000 |

**HPLC chromatogram of S-MYM-III-10 at 50 minutes.** The racemization occurred within 50 minutes in solution. Normal phase chiral HPLC, Chiral pak IBN3 column (dimensions 4.6 mm x 150 mm, particle size 3 μm, mobile phase 10% EtOH-hexane)

SHIMADZU LabSolutions

### Analysis Report

#### <Sample Information>

| | | | |
|---|---|---|---|
| Sample Name | : MYM-III-10 (45) | | |
| Sample ID | : MYM-III-10 (45) | | |
| Data Filename | : 8-17-21_Yeunus, MYM-III-10 (45), EtOH-Hex=12-88_52 mi.lcd | | |
| Method Filename | : 8-30-21-EtOH-Hex=12-88_52 min.lcm | | |
| Batch Filename | : | | |
| Vial # | : Vial 2 | Sample Type | : Unknown |
| Injection Volume | : 20 uL | Level | : 1 |
| Date Acquired | : 8/31/2021 2:32:45 PM | Acquired by | : Md Yeunus Mian |
| Date Processed | : 8/31/2021 3:25:52 PM | Processed by | : Md Yeunus Mian |

#### <Chromatogram>

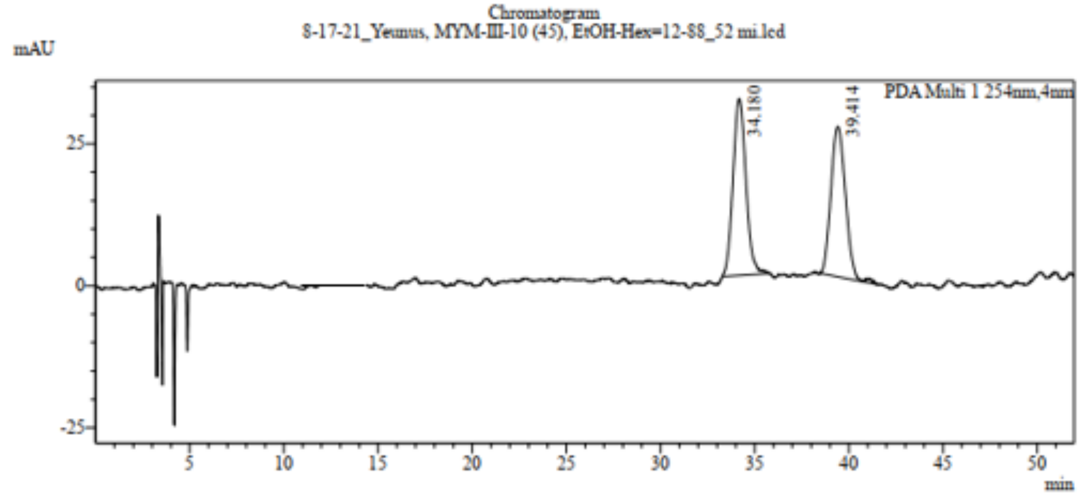


#### <Peak Table>

8-17-21_Yeunus, MYM-III-10 (45), EtOH-Hex=12-88_52 mi.lcd

(3D)DAD Ch1 254nm

| Peak# | Ret. Time | Area% |
|---|---|---|
| 1 | 34.180 | 50.989 |
| 2 | 39.414 | 49.011 |
| Total | | 100.000 |

**HPLC chromatogram of R-MYM-III-10 at 40 minutes.** Racemization occurs with 40 minutes in solution. Normal phase chiral HPLC, Chiral pak IBN3 column (dimensions 4.6 mm x 150 mm, particle size 3 μm, mobile phase 10% EtOH-hexane)

**Racemization of MYM-V-56:** We also investigated the enantiomers of MYM-V-56 in solution after separation using the preparative HPLC column mentioned in the previous section. Both enantiomers were kept in solution for 8 h and monitored by HPLC in analytical column at different time points. Both enantiomers were kept in ethanol solution for more than 8 hours but no evidence of racemization was found.

SHIMADZU LabSolutions **Analysis Report**

**<Sample Information>**

| | | | |
|---|---|---|---|
| Sample Name | : rac-56-2 | | |
| Sample ID | : rac-56-2 | | |
| Data Filename | : rac-56-2_8-17-21-EtOH-Hex=12-88_80 min.lcm_8-17-2_002.lcd | | |
| Method Filename | : 8-17-21-EtOH-Hex=12-88_80 min.lcm | | |
| Batch Filename | : 8-17-2.lcb | | |
| Vial # | : Vial 3 | Sample Type | : Unknown |
| Injection Volume | : 20 uL | Level | : 1 |
| Date Acquired | : 8/17/2021 8:50:51 PM | Acquired by | : Md Yeunus Mian |
| Date Processed | : 8/17/2021 10:11:57 PM | Processed by | : Md Yeunus Mian |

**<Chromatogram>**

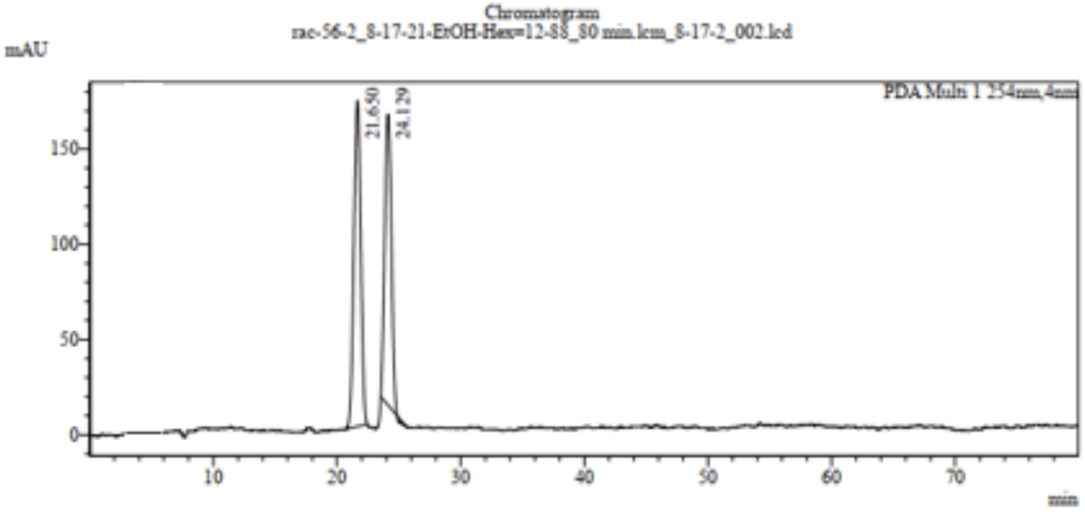


**<Peak Table>**

rac-56-2_8-17-21-EtOH-Hex=12-88_80 min.lcm_8-17-2_002.lcd

(3D)DAD Ch1 254nm

| Peak# | Ret. Time | Area% |
|---|---|---|
| 1 | 21.650 | 53.846 |
| 2 | 24.129 | 46.154 |
| Total | | 100.000 |

**HPLC chromatogram of racemic MYM-V-56.** Normal phase chiral HPLC, Whelk-01 column (4.6 mm x 25 cm, 5 μm particle size, mobile phase 12% EtOH-hexane.

**<Sample Information>**

| | | | |
|---|---|---|---|
| Sample Name | : V_56-1 | | |
| Sample ID | : V_56-1 | | |
| Data Filename | : V_56-1_8-31-21-EtOH-Hex=12-88_35 min.lcm_v-56_001.lcd | | |
| Method Filename | : 8-31-21-EtOH-Hex=12-88_35 min.lcm | | |
| Batch Filename | : v-56.lcb | | |
| Vial # | : Vial 1 | Sample Type | : Unknown |
| Injection Volume | : 20 uL | Level | : 1 |
| Date Acquired | : 8/31/2021 5:53:41 PM | Acquired by | : Md Yeunus Mian |
| Date Processed | : 8/31/2021 8:07:19 PM | Processed by | : Md Yeunus Mian |

**<Chromatogram>**

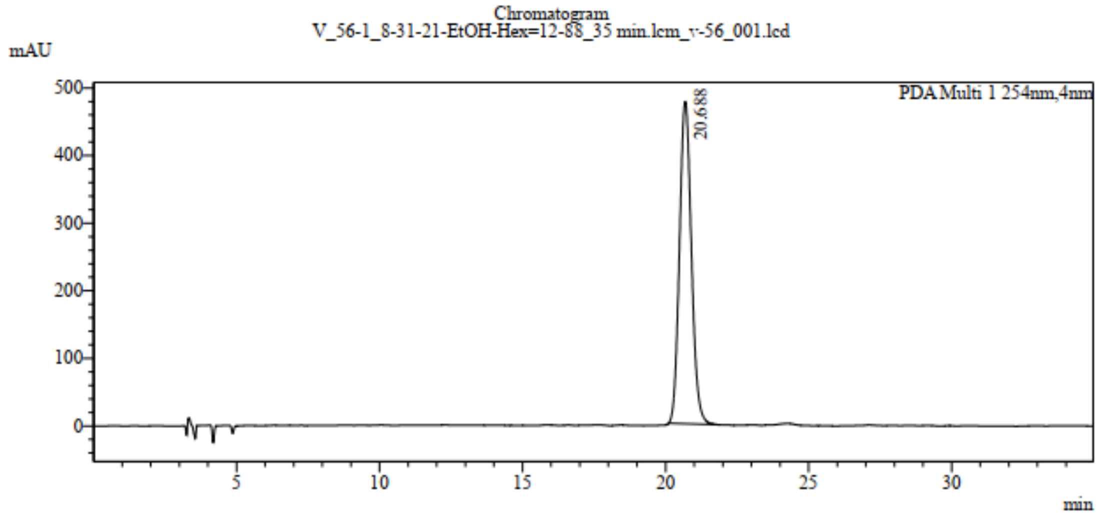


**<Peak Table>**

V_56-1_8-31-21-EtOH-Hex=12-88_35 min.lcm_v-56_001.lcd

(3D)DAD Ch1 254nm

| Peak# | Ret. Time | Area% |
|---|---|---|
| 1 | 20.688 | 100.000 |
| Total | | 100.000 |

**HPLC chromatogram of optically pure R-MYM-V-56 after 8 hr in ethanol solution.** No racemization was obtained like MYM-III-10. Normal phase chiral HPLC, Whelk-01 column (4.6 mm x 25 cm, 5 μm particle size, mobile phase 12% EtOH-hexane.

#### <Sample Information>

| | |
|---|---|
| Sample Name | : V_56-2 |
| Sample ID | : V_56-2 |
| Data Filename | : V_56-2_8-31-21-EtOH-Hex=12-88_35 min.lcm_v-56_002.lcd |
| Method Filename | : 8-31-21-EtOH-Hex=12-88_35 min.lcm |
| Batch Filename | : v-56.lcb |

| | | | |
|---|---|---|---|
| Vial # | : Vial 2 | Sample Type | : Unknown |
| Injection Volume | : 20 uL | Level | : 1 |
| Date Acquired | : 8/31/2021 6:29:52 PM | Acquired by | : Md Yeunus Mian |
| Date Processed | : 8/31/2021 7:06:02 PM | Processed by | : Md Yeunus Mian |

#### <Chromatogram>

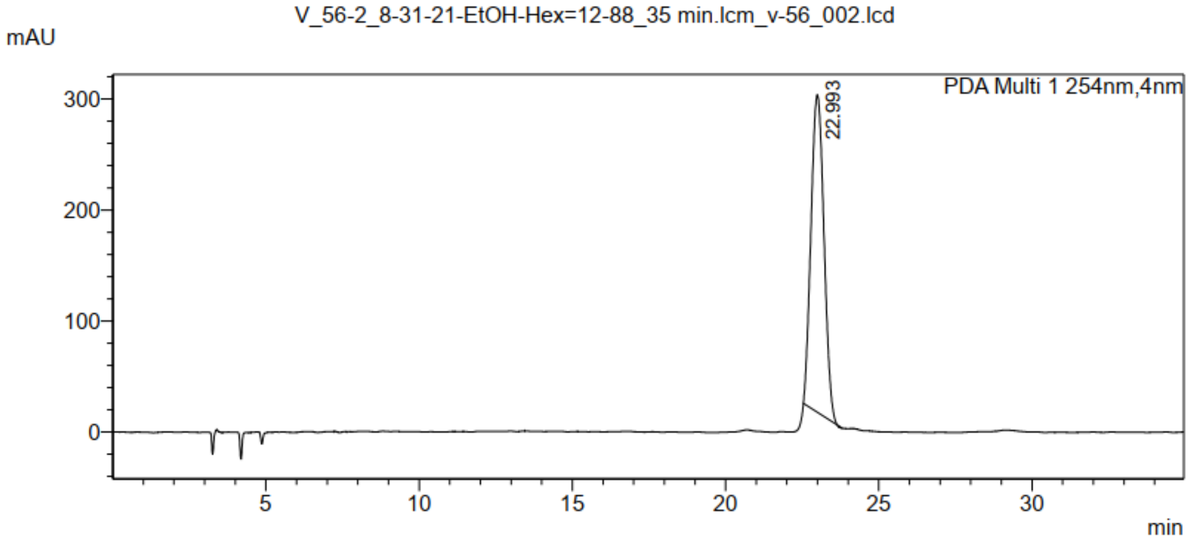


#### <Peak Table>

V_56-2_8-31-21-EtOH-Hex=12-88_35 min.lcm_v-56_002.lcd

(3D)DAD Ch1 254nm

| Peak# | Ret. Time | Area% |
|---|---|---|
| 1 | 22.993 | 100.000 |
| Total | | 100.000 |

**HPLC chromatogram of optically pure S-MYM-V-56 after 8 hr in ethanol solution.** No racemization was obtained unlike MYM-III-10. Normal phase chiral HPLC, Whelk-01 column (4.6 mm x 25 cm, 5 µm particle size, mobile phase 12% EtOH-hexane.

**X-ray crystal structure data for MYM-V-56.**

**Table 1**. Crystal data and structure refinement for MYM-V-56.

| | | |
|---|---|---|
| Identification code | cook195_twinb | |
| Empirical formula | $C_{16}H_{11}Cl_3FN_3O_3$ | |
| Formula weight | 418.63 | |
| Temperature | 293(2) K | |
| Wavelength | 1.54178 Å | |
| Crystal system | Monoclinic | |
| Space group | $P2_1$ | |
| Unit cell dimensions | a = 7.1939(2) Å | a= 90°. |
| | b = 8.3595(2) Å | b= 96.4475(12)°. |
| | c = 15.1285(4) Å | g = 90°. |
| Volume | 904.03(4) $Å^3$ | |
| Z | 2 | |
| Density (calculated) | 1.538 $Mg/m^3$ | |
| Absorption coefficient | 4.888 $mm^{-1}$ | |
| F(000) | 424 | |
| Crystal size | 0.236 x 0.185 x 0.080 $mm^3$ | |
| Theta range for data collection | 5.887 to 74.579°. | |
| Index ranges | -8<=h<=8, -10<=k<=10, -18<=l<=18 | |
| Reflections collected | 3150 | |
| Independent reflections | 3150 [$R_{int}$ = 0.032] | |
| Completeness to theta = 67.679° | 99.5 % | |
| Absorption correction | Semi-empirical from equivalents | |
| Max. and min. transmission | 0.7538 and 0.5237 | |
| Refinement method | Full-matrix least-squares on $F^2$ | |
| Data / restraints / parameters | 3150 / 1 / 236 | |
| Goodness-of-fit on $F^2$ | 1.025 | |
| Final R indices [I>2sigma(I)] | $R_1$ = 0.0454, $wR_2$ = 0.1306 | |
| R indices (all data) | $R_1$ = 0.0460, $wR_2$ = 0.1319 | |
| Absolute structure parameter | 0.009(12) | |
| Largest diff. peak and hole | 0.326 and -0.344 e.$Å^{-3}$ | |

**Table 2**. Atomic coordinates ( x $10^4$) and equivalent isotropic displacement parameters ($Å^2$x $10^3$) for MYM-V-56. U(eq) is defined as one third of the trace of the orthogonalized $U^{ij}$ tensor.

| | x | y | z | U(eq) |
|---|---|---|---|---|
| C(1) | 5926(6) | 667(4) | 7833(2) | 41(1) |
| C(2) | 5346(5) | 1514(4) | 8550(2) | 37(1) |
| C(3) | 4595(5) | 3016(5) | 8455(2) | 35(1) |
| C(4) | 4415(5) | 3767(4) | 7625(2) | 33(1) |
| C(5) | 5024(5) | 2944(4) | 6900(2) | 33(1) |
| C(6) | 5741(6) | 1392(4) | 7011(2) | 39(1) |
| C(7) | 3609(5) | 5402(4) | 7547(2) | 37(1) |
| N(8) | 4192(6) | 6505(4) | 7063(2) | 46(1) |
| C(9) | 5748(7) | 6190(5) | 6587(3) | 47(1) |
| C(10) | 5135(7) | 5100(5) | 5794(2) | 46(1) |
| N(11) | 4840(5) | 3565(4) | 6028(2) | 42(1) |
| C(12) | 2105(6) | 5865(5) | 8103(2) | 45(1) |
| C(13) | 618(7) | 4954(6) | 8265(3) | 54(1) |
| C(14) | -712(8) | 5479(9) | 8812(4) | 72(2) |
| C(15) | -511(11) | 6948(10) | 9189(5) | 93(2) |
| C(16) | 935(12) | 7924(9) | 9031(5) | 94(2) |
| C(17) | 2258(9) | 7381(6) | 8497(4) | 67(1) |
| N(18) | 5493(6) | 751(4) | 9428(2) | 50(1) |
| O(19) | 6630(8) | -311(6) | 9576(2) | 93(2) |
| O(20) | 4505(6) | 1204(6) | 9962(2) | 79(1) |
| F(21) | 6309(6) | 7625(3) | 6251(2) | 76(1) |
| O(22) | 4969(8) | 5566(5) | 5035(2) | 69(1) |
| Cl(23) | 186(2) | 3087(2) | 7775(1) | 87(1) |
| Cl(24) | 518(7) | 9942(9) | 6435(3) | 247(3) |
| Cl(25) | 284(5) | 9298(7) | 4546(2) | 171(2) |
| C(26) | 1090(20) | 8740(18) | 5646(8) | 150(5) |

**Table 3**. Bond lengths [Å] and angles [°] for MYM-V-56.

| | | | |
|---|---|---|---|
| C(1)-C(6) | 1.376(5) | C(1)-C(2) | 1.397(5) |
| C(1)-H(1) | 0.9300 | C(2)-C(3) | 1.369(5) |
| C(2)-N(18) | 1.466(4) | C(3)-C(4) | 1.396(5) |
| C(3)-H(3) | 0.9300 | C(4)-C(5) | 1.406(4) |
| C(4)-C(7) | 1.484(5) | C(5)-C(6) | 1.400(5) |
| C(5)-N(11) | 1.410(4) | C(6)-H(6) | 0.9300 |
| C(7)-N(8) | 1.277(5) | C(7)-C(12) | 1.495(5) |
| N(8)-C(9) | 1.422(6) | C(9)-F(21) | 1.381(4) |
| C(9)-C(10) | 1.532(6) | C(9)-H(9) | 0.9800 |
| C(10)-O(22) | 1.205(5) | C(10)-N(11) | 1.354(5) |
| N(11)-H(11) | 0.8600 | C(12)-C(13) | 1.358(7) |
| C(12)-C(17) | 1.399(7) | C(13)-C(14) | 1.404(7) |
| C(13)-Cl(23) | 1.741(6) | C(14)-C(15) | 1.355(11) |
| C(14)-H(14) | 0.9300 | C(15)-C(16) | 1.364(12) |
| C(15)-H(15) | 0.9300 | C(16)-C(17) | 1.391(8) |
| C(16)-H(16) | 0.9300 | C(17)-H(17) | 0.9300 |
| N(18)-O(20) | 1.197(5) | N(18)-O(19) | 1.211(6) |
| Cl(24)-C(26) | 1.646(14) | Cl(25)-C(26) | 1.763(13) |
| C(26)-H(26A) | 0.9700 | C(26)-H(26B) | 0.9700 |
| C(6)-C(1)-C(2) | 118.1(3) | C(6)-C(1)-H(1) | 120.9 |
| C(2)-C(1)-H(1) | 120.9 | C(3)-C(2)-C(1) | 122.1(3) |
| C(3)-C(2)-N(18) | 118.8(3) | C(1)-C(2)-N(18) | 119.1(3) |
| C(2)-C(3)-C(4) | 120.1(3) | C(2)-C(3)-H(3) | 119.9 |
| C(4)-C(3)-H(3) | 119.9 | C(3)-C(4)-C(5) | 118.5(3) |
| C(3)-C(4)-C(7) | 118.8(3) | C(5)-C(4)-C(7) | 122.7(3) |
| C(6)-C(5)-C(4) | 120.2(3) | C(6)-C(5)-N(11) | 116.6(3) |
| C(4)-C(5)-N(11) | 123.1(3) | C(1)-C(6)-C(5) | 120.9(3) |
| C(1)-C(6)-H(6) | 119.6 | C(5)-C(6)-H(6) | 119.6 |
| N(8)-C(7)-C(4) | 124.1(3) | N(8)-C(7)-C(12) | 116.1(3) |
| C(4)-C(7)-C(12) | 119.6(3) | C(7)-N(8)-C(9) | 119.0(3) |
| F(21)-C(9)-N(8) | 107.8(3) | F(21)-C(9)-C(10) | 107.4(3) |
| N(8)-C(9)-C(10) | 109.5(4) | F(21)-C(9)-H(9) | 110.7 |

| | | | |
|---|---|---|---|
| N(8)-C(9)-H(9) | 110.7 | C(10)-C(9)-H(9) | 110.7 |
| O(22)-C(10)-N(11) | 123.6(4) | O(22)-C(10)-C(9) | 122.9(4) |
| N(11)-C(10)-C(9) | 113.5(3) | C(10)-N(11)-C(5) | 126.3(3) |
| C(10)-N(11)-H(11) | 116.9 | C(5)-N(11)-H(11) | 116.9 |
| C(13)-C(12)-C(17) | 117.2(4) | C(13)-C(12)-C(7) | 126.2(4) |
| C(17)-C(12)-C(7) | 116.6(4) | C(12)-C(13)-C(14) | 122.3(5) |
| C(12)-C(13)-Cl(23) | 122.2(4) | C(14)-C(13)-Cl(23) | 115.5(4) |
| C(15)-C(14)-C(13) | 118.9(6) | C(15)-C(14)-H(14) | 120.5 |
| C(13)-C(14)-H(14) | 120.5 | C(14)-C(15)-C(16) | 121.0(5) |
| C(14)-C(15)-H(15) | 119.5 | C(16)-C(15)-H(15) | 119.5 |
| C(15)-C(16)-C(17) | 119.4(6) | C(15)-C(16)-H(16) | 120.3 |
| C(17)-C(16)-H(16) | 120.3 | C(16)-C(17)-C(12) | 121.1(6) |
| C(16)-C(17)-H(17) | 119.4 | C(12)-C(17)-H(17) | 119.4 |
| O(20)-N(18)-O(19) | 123.1(4) | O(20)-N(18)-C(2) | 119.1(4) |
| O(19)-N(18)-C(2) | 117.7(3) | Cl(24)-C(26)-Cl(25) | 116.2(9) |
| Cl(24)-C(26)-H(26A) | 108.2 | Cl(25)-C(26)-H(26A) | 108.2 |
| Cl(24)-C(26)-H(26B) | 108.2 | Cl(25)-C(26)-H(26B) | 108.2 |
| H(26A)-C(26)-H(26B) | 107.4 | | |

**Table 4**. Anisotropic displacement parameters (Å$^2$x 10$^3$) for MYM-V-56. The anisotropic displacement factor exponent takes the form: $-2p^2[h^2a^{*2}U^{11} + ... + 2 h k a^* b^* U^{12}]$

| | $U^{11}$ | $U^{22}$ | $U^{33}$ | $U^{23}$ | $U^{13}$ | $U^{12}$ |
|---|---|---|---|---|---|---|
| C(1) | 56(2) | 30(2) | 37(2) | 2(1) | 10(2) | 3(2) |
| C(2) | 46(2) | 37(2) | 29(2) | 2(1) | 7(1) | -2(2) |
| C(3) | 39(2) | 38(2) | 27(1) | -2(1) | 8(1) | -2(1) |
| C(4) | 40(2) | 32(2) | 29(1) | -3(1) | 10(1) | -3(1) |
| C(5) | 43(2) | 32(2) | 26(1) | -2(1) | 8(1) | -3(1) |
| C(6) | 54(2) | 34(2) | 29(1) | -2(1) | 10(1) | 3(2) |
| C(7) | 47(2) | 32(2) | 32(1) | -1(1) | 7(1) | 0(1) |
| N(8) | 67(2) | 35(2) | 39(1) | 1(1) | 16(2) | 2(2) |
| C(9) | 69(3) | 33(2) | 43(2) | 2(2) | 23(2) | -7(2) |
| C(10) | 66(3) | 41(2) | 33(2) | 6(1) | 16(2) | 4(2) |
| N(11) | 64(2) | 38(2) | 25(1) | -2(1) | 7(1) | 0(2) |
| C(12) | 52(2) | 42(2) | 43(2) | 2(1) | 12(2) | 7(2) |
| C(13) | 46(2) | 61(3) | 55(2) | 6(2) | 6(2) | 6(2) |
| C(14) | 52(3) | 94(4) | 73(3) | 9(3) | 20(2) | 9(3) |
| C(15) | 92(5) | 95(5) | 102(4) | 10(4) | 59(4) | 39(4) |
| C(16) | 122(6) | 66(4) | 105(5) | -19(3) | 61(4) | 17(4) |
| C(17) | 86(4) | 48(3) | 72(3) | -13(2) | 37(3) | 7(2) |
| N(18) | 71(2) | 44(2) | 33(1) | 9(1) | 9(1) | 6(2) |
| O(19) | 147(4) | 77(3) | 58(2) | 30(2) | 31(2) | 58(3) |
| O(20) | 101(3) | 96(3) | 44(2) | 25(2) | 32(2) | 29(3) |
| F(21) | 119(3) | 40(1) | 77(2) | 6(1) | 48(2) | -17(2) |
| O(22) | 121(3) | 56(2) | 34(1) | 13(1) | 19(2) | 6(2) |
| Cl(23) | 64(1) | 92(1) | 105(1) | -31(1) | 11(1) | -31(1) |
| Cl(24) | 185(4) | 317(7) | 231(4) | -154(5) | -19(3) | 87(5) |
| Cl(25) | 120(2) | 240(5) | 150(2) | -15(3) | -1(2) | -20(3) |
| C(26) | 149(10) | 129(9) | 185(11) | 40(9) | 71(9) | 15(8) |

**Table 5**. Hydrogen coordinates (x $10^4$) and isotropic displacement parameters ($Å^2$x $10^3$) for MYM-V-56.

| | x | y | z | U(eq) |
|---|---|---|---|---|
| H(1) | 6423 | -357 | 7909 | 49 |
| H(3) | 4202 | 3538 | 8944 | 41 |
| H(6) | 6096 | 843 | 6522 | 46 |
| H(9) | 6772 | 5700 | 6977 | 56 |
| H(11) | 4509 | 2904 | 5603 | 51 |
| H(14) | -1716 | 4830 | 8913 | 86 |
| H(15) | -1371 | 7296 | 9563 | 111 |
| H(16) | 1035 | 8944 | 9277 | 113 |
| H(17) | 3261 | 8035 | 8401 | 80 |
| H(26A) | 606 | 7682 | 5753 | 181 |
| H(26B) | 2441 | 8661 | 5697 | 181 |

**Table 6**. Torsion angles [°] for MYM-V-56.

| | | | |
|---|---|---|---|
| C(6)-C(1)-C(2)-C(3) | 0.6(6) | C(6)-C(1)-C(2)-N(18) | 178.7(4) |
| C(1)-C(2)-C(3)-C(4) | -1.2(6) | N(18)-C(2)-C(3)-C(4) | -179.2(3) |
| C(2)-C(3)-C(4)-C(5) | -0.1(5) | C(2)-C(3)-C(4)-C(7) | -179.5(3) |
| C(3)-C(4)-C(5)-C(6) | 1.8(5) | C(7)-C(4)-C(5)-C(6) | -178.8(3) |
| C(3)-C(4)-C(5)-N(11) | 177.4(3) | C(7)-C(4)-C(5)-N(11) | -3.2(6) |
| C(2)-C(1)-C(6)-C(5) | 1.2(6) | C(4)-C(5)-C(6)-C(1) | -2.4(6) |
| N(11)-C(5)-C(6)-C(1) | -178.3(4) | C(3)-C(4)-C(7)-N(8) | 141.3(4) |
| C(5)-C(4)-C(7)-N(8) | -38.1(6) | C(3)-C(4)-C(7)-C(12) | -34.4(5) |
| C(5)-C(4)-C(7)-C(12) | 146.2(4) | C(4)-C(7)-N(8)-C(9) | -2.1(6) |
| C(12)-C(7)-N(8)-C(9) | 173.7(4) | C(7)-N(8)-C(9)-F(21) | -169.5(4) |
| C(7)-N(8)-C(9)-C(10) | 74.1(5) | F(21)-C(9)-C(10)-O(22) | -8.9(7) |
| N(8)-C(9)-C(10)-O(22) | 107.9(5) | F(21)-C(9)-C(10)-N(11) | 169.7(4) |
| N(8)-C(9)-C(10)-N(11) | -73.6(5) | O(22)-C(10)-N(11)-C(5) | 179.5(5) |
| C(9)-C(10)-N(11)-C(5) | 0.9(6) | C(6)-C(5)-N(11)-C(10) | -143.4(4) |
| C(4)-C(5)-N(11)-C(10) | 40.8(6) | N(8)-C(7)-C(12)-C(13) | 141.2(4) |
| C(4)-C(7)-C(12)-C(13) | -42.8(6) | N(8)-C(7)-C(12)-C(17) | -39.2(6) |
| C(4)-C(7)-C(12)-C(17) | 136.9(4) | C(17)-C(12)-C(13)-C(14) | -0.8(7) |
| C(7)-C(12)-C(13)-C(14) | 178.8(4) | C(17)-C(12)-C(13)-Cl(23) | 176.7(4) |
| C(7)-C(12)-C(13)-Cl(23) | -3.7(6) | C(12)-C(13)-C(14)-C(15) | 0.2(8) |
| Cl(23)-C(13)-C(14)-C(15) | -177.5(5) | C(13)-C(14)-C(15)-C(16) | 1.4(11) |
| C(14)-C(15)-C(16)-C(17) | -2.3(13) | C(15)-C(16)-C(17)-C(12) | 1.6(12) |
| C(13)-C(12)-C(17)-C(16) | -0.1(9) | C(7)-C(12)-C(17)-C(16) | -179.8(6) |
| C(3)-C(2)-N(18)-O(20) | 21.6(6) | C(1)-C(2)-N(18)-O(20) | -156.5(5) |
| C(3)-C(2)-N(18)-O(19) | -157.9(5) | C(1)-C(2)-N(18)-O(19) | 24.0(6) |

**Table 7**. Hydrogen bonds for MYM-V-56 [Å and °].

| D-H...A | d(D-H) | d(H...A) | d(D...A) | <(DHA) |
|---|---|---|---|---|
| C(3)-H(3)...O(19)#1 | 0.93 | 2.57 | 3.492(5) | 172.2 |
| C(6)-H(6)...O(22)#2 | 0.93 | 2.41 | 3.156(4) | 137.7 |
| N(11)-H(11)...O(22)#2 | 0.86 | 2.23 | 2.990(5) | 147.1 |
| C(26)-H(26A)...Cl(25)#3 | 0.97 | 2.92 | 3.844(16) | 158.8 |

Symmetry transformations used to generate equivalent atoms:

#1 -x+1,y+1/2,-z+2    #2 -x+1,y-1/2,-z+1    #3 -x,y-1/2,-z+1

**HNMR of MYM-II-53 (2):**

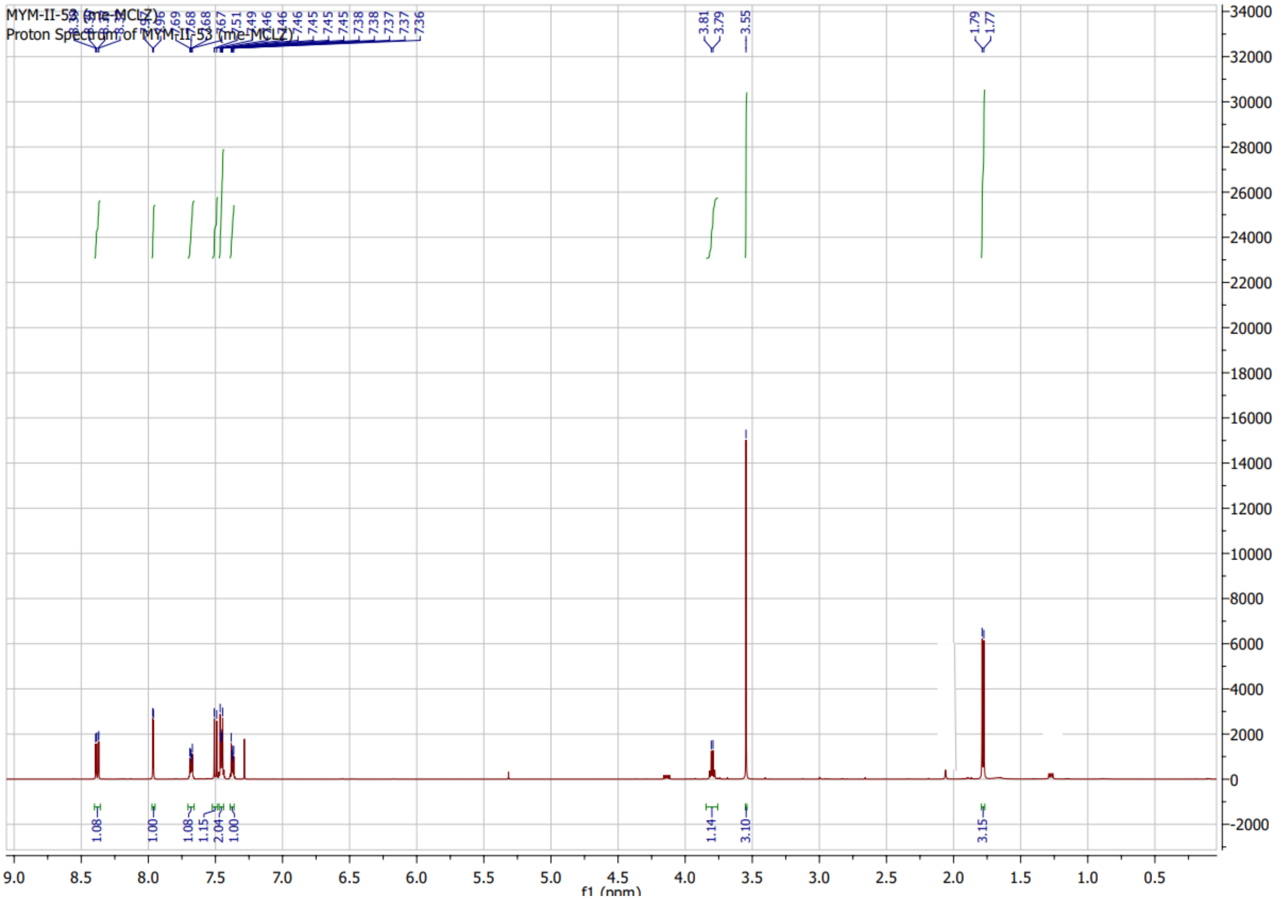

**CNMR of MYM-II-53 (2):**

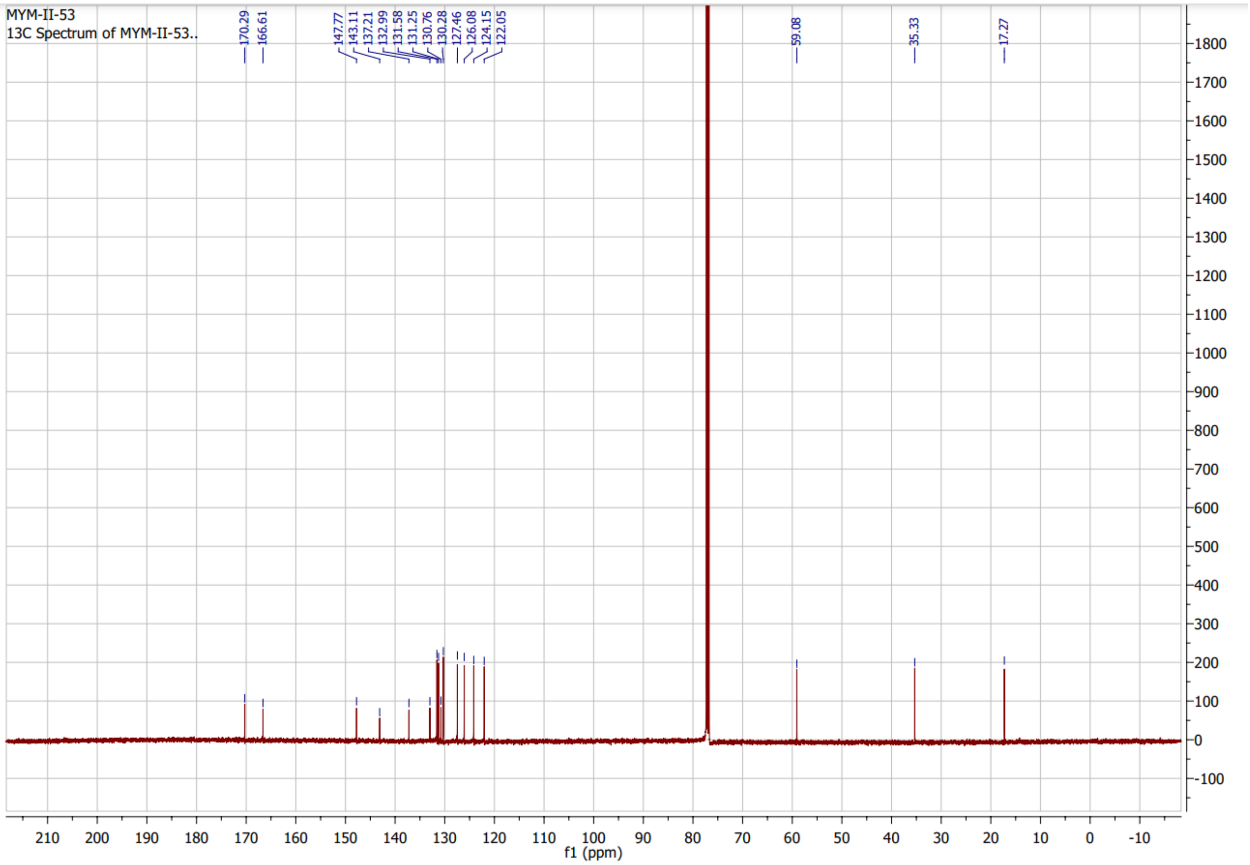

**HNMR of MYM-II-81 (4):**

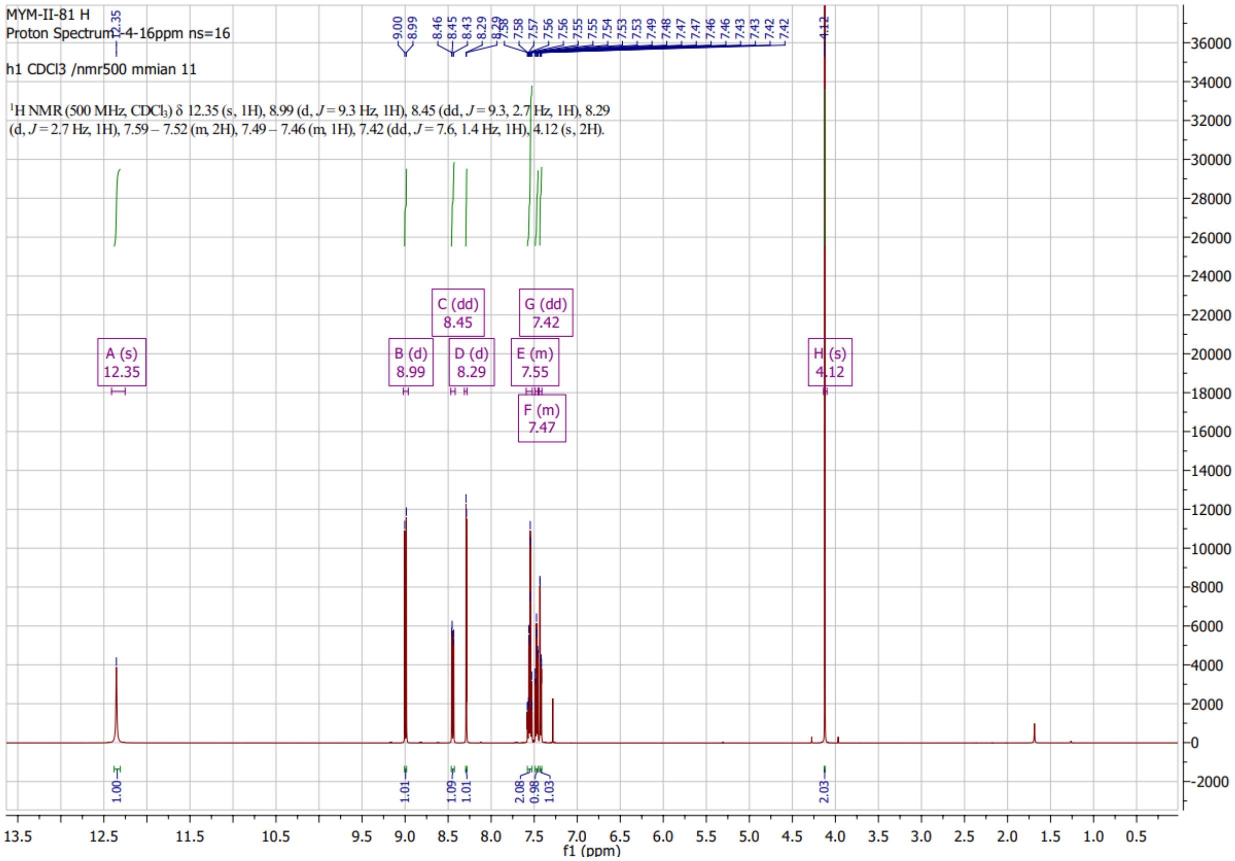

**CNMR of MYM-II-81 (4):**

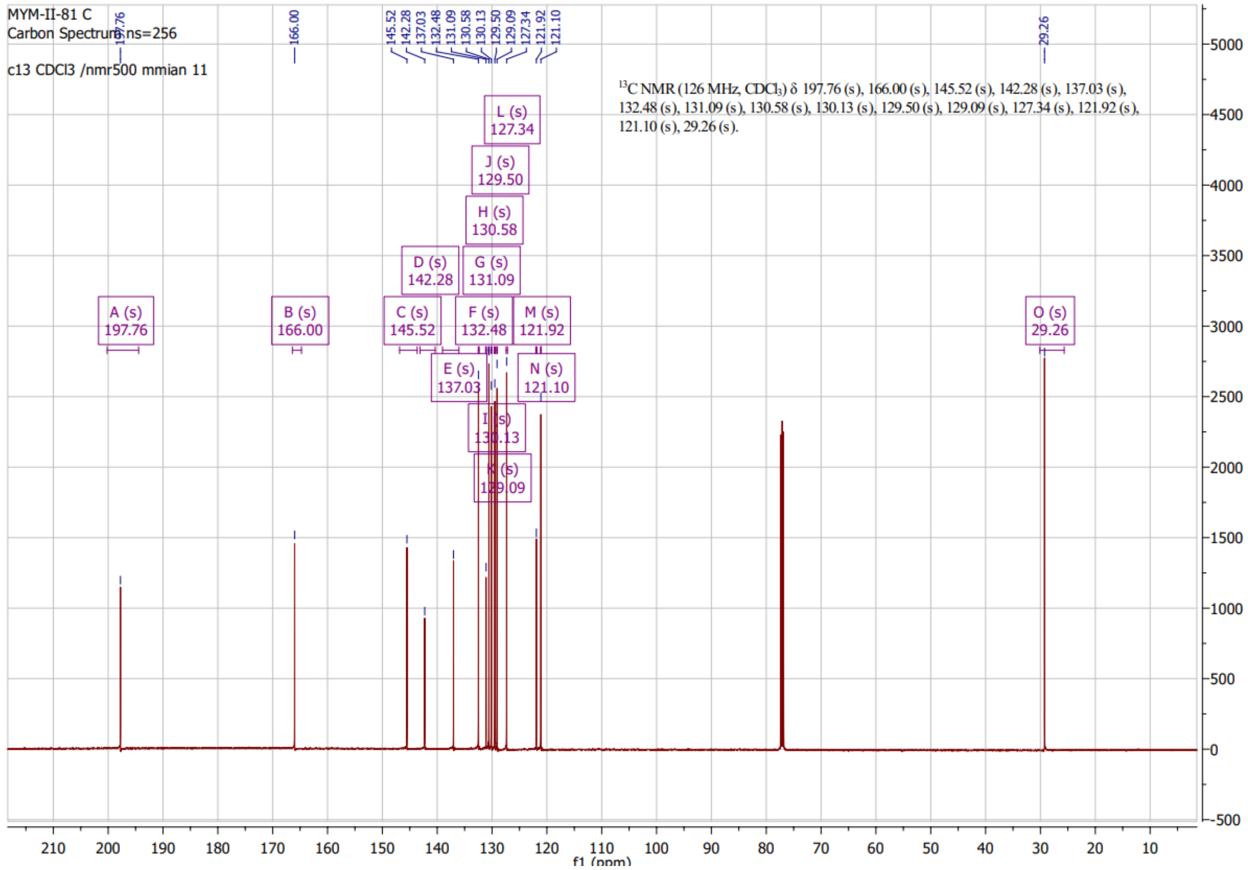

#### HNMR of MYM-III-50 (5)

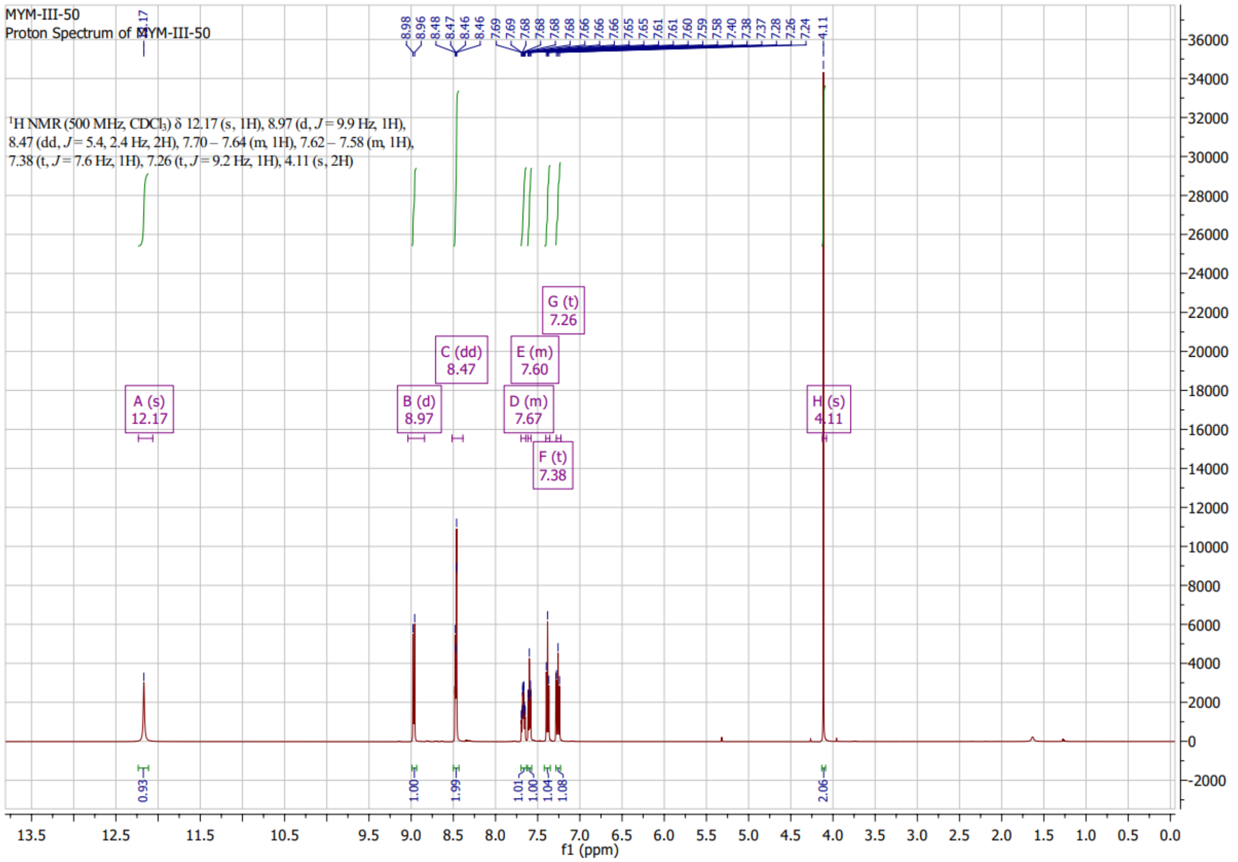

#### CNMR of MYM-III-50 (5):

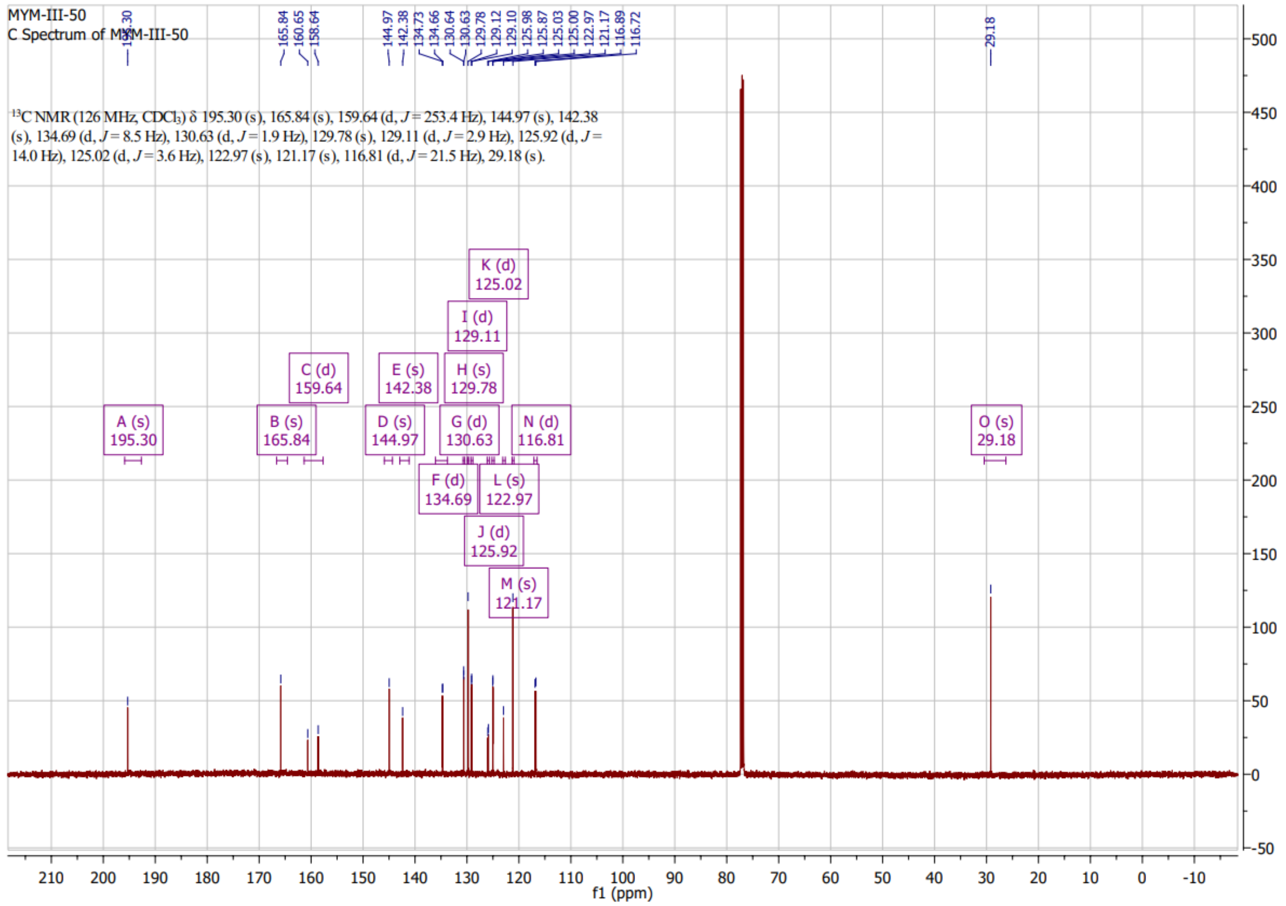

**HNMR of MYM-II-82(6)**

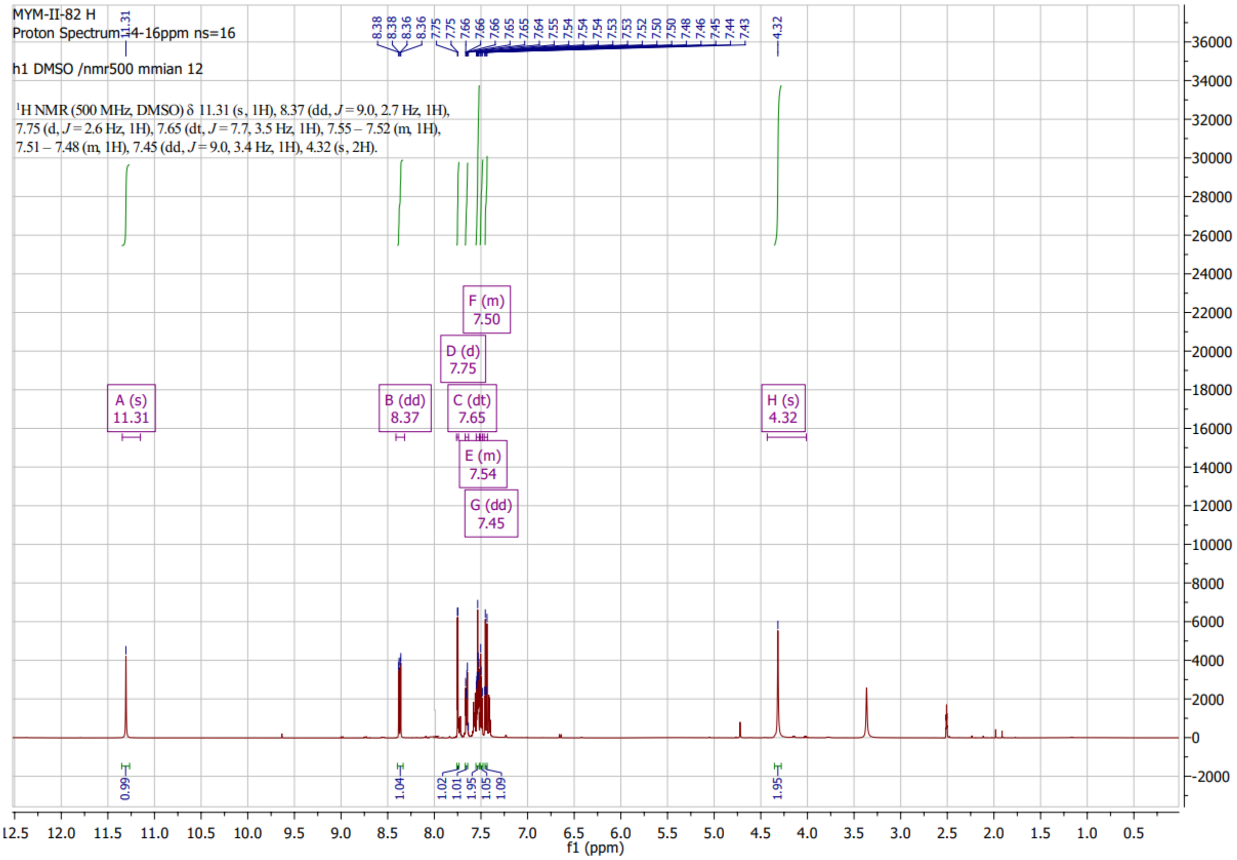

#### CNMR of MYM-II-82(6)

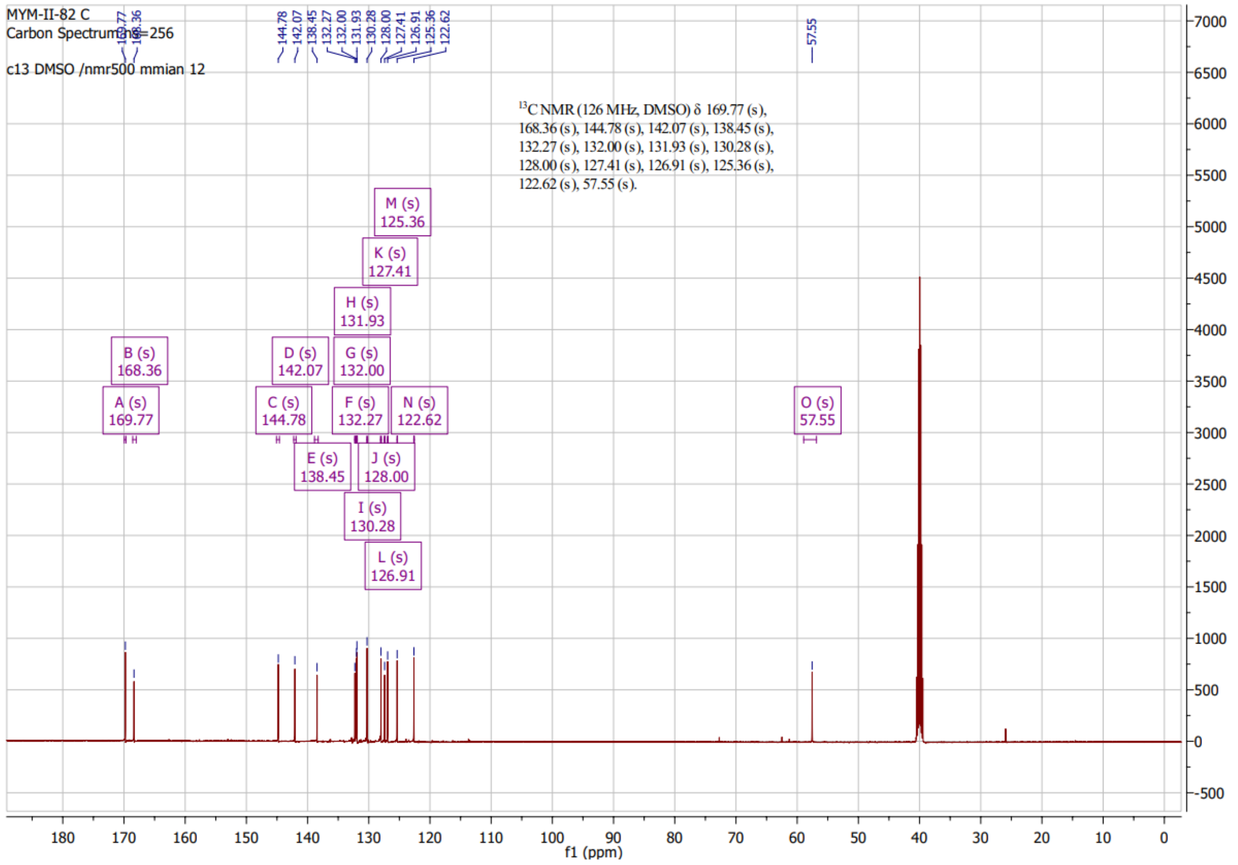

#### HNMR of MYM-III-55 (7)

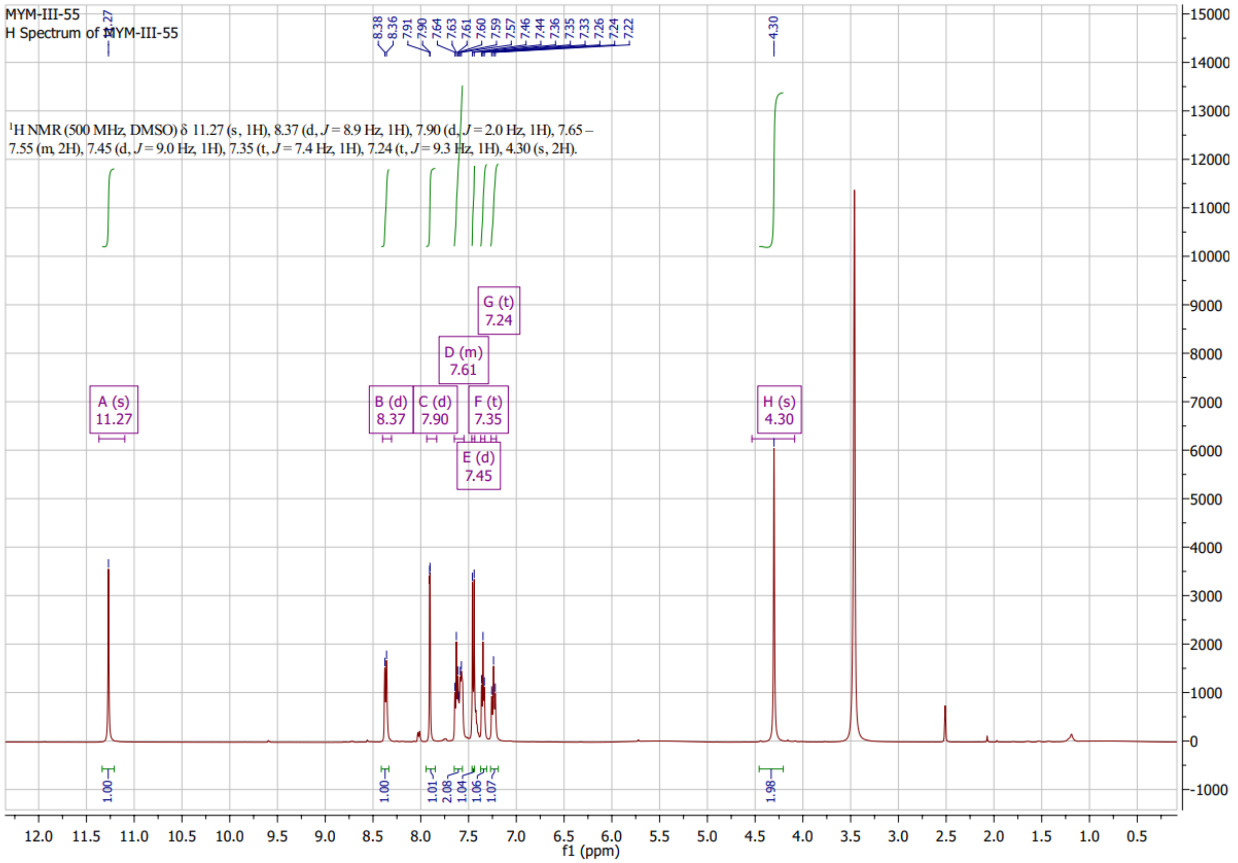

#### CNMR of MYM-III-55 (7)

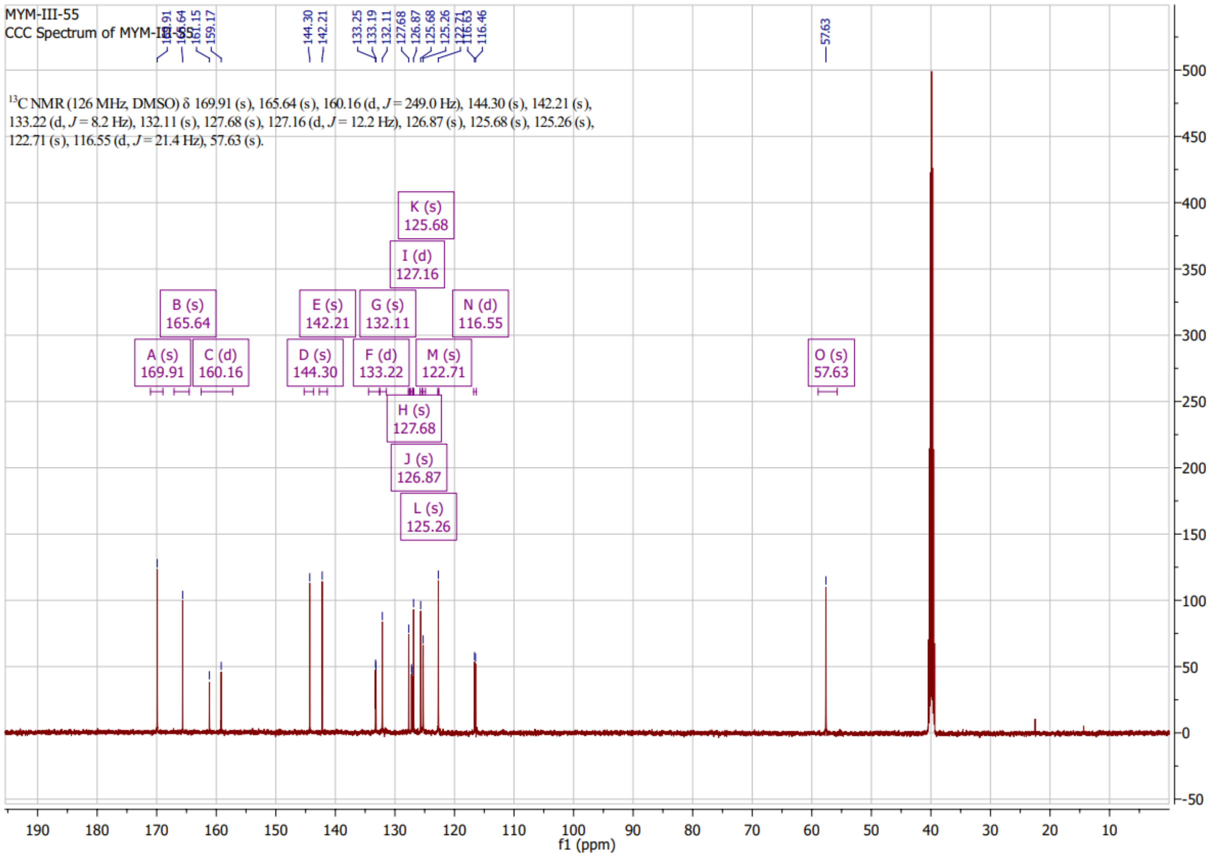

### HNMR of MYM-II-83 (8)

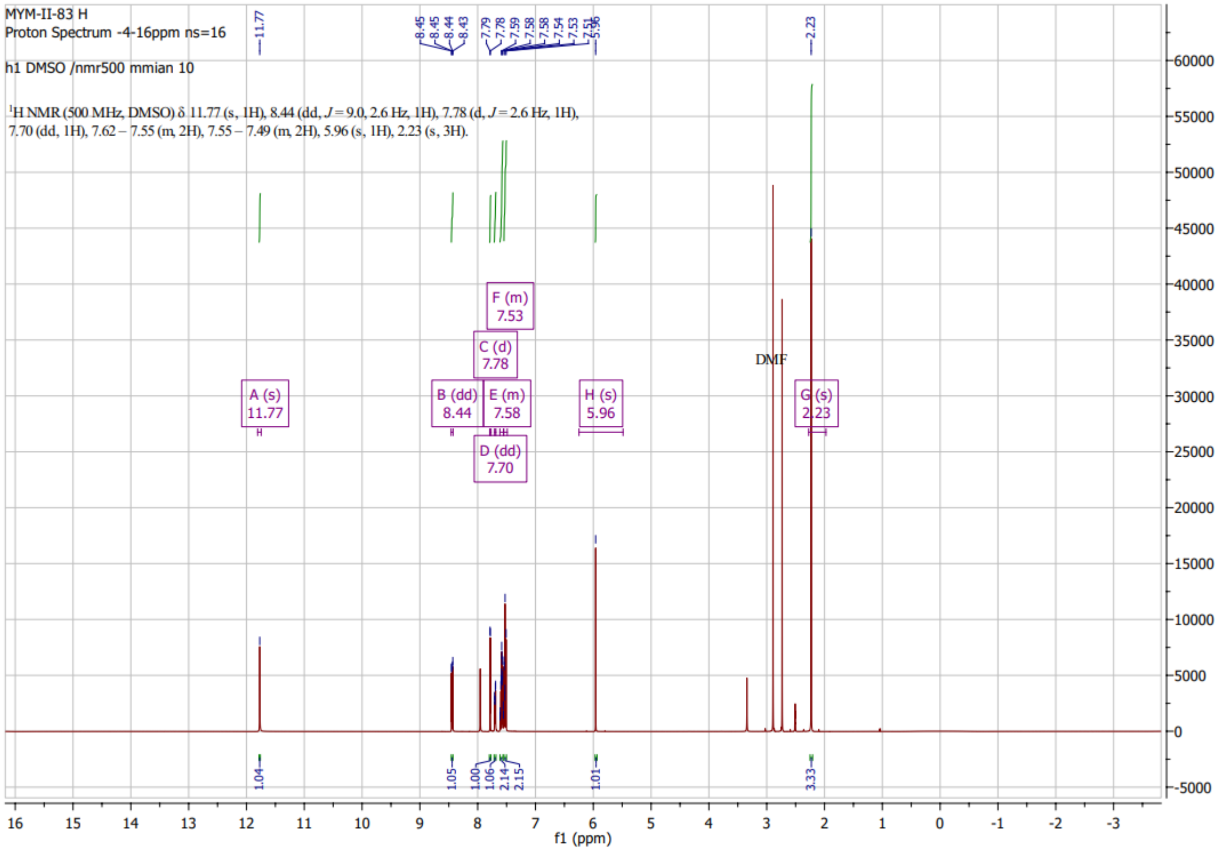

**CNMR of MYM-II-83 (8):**

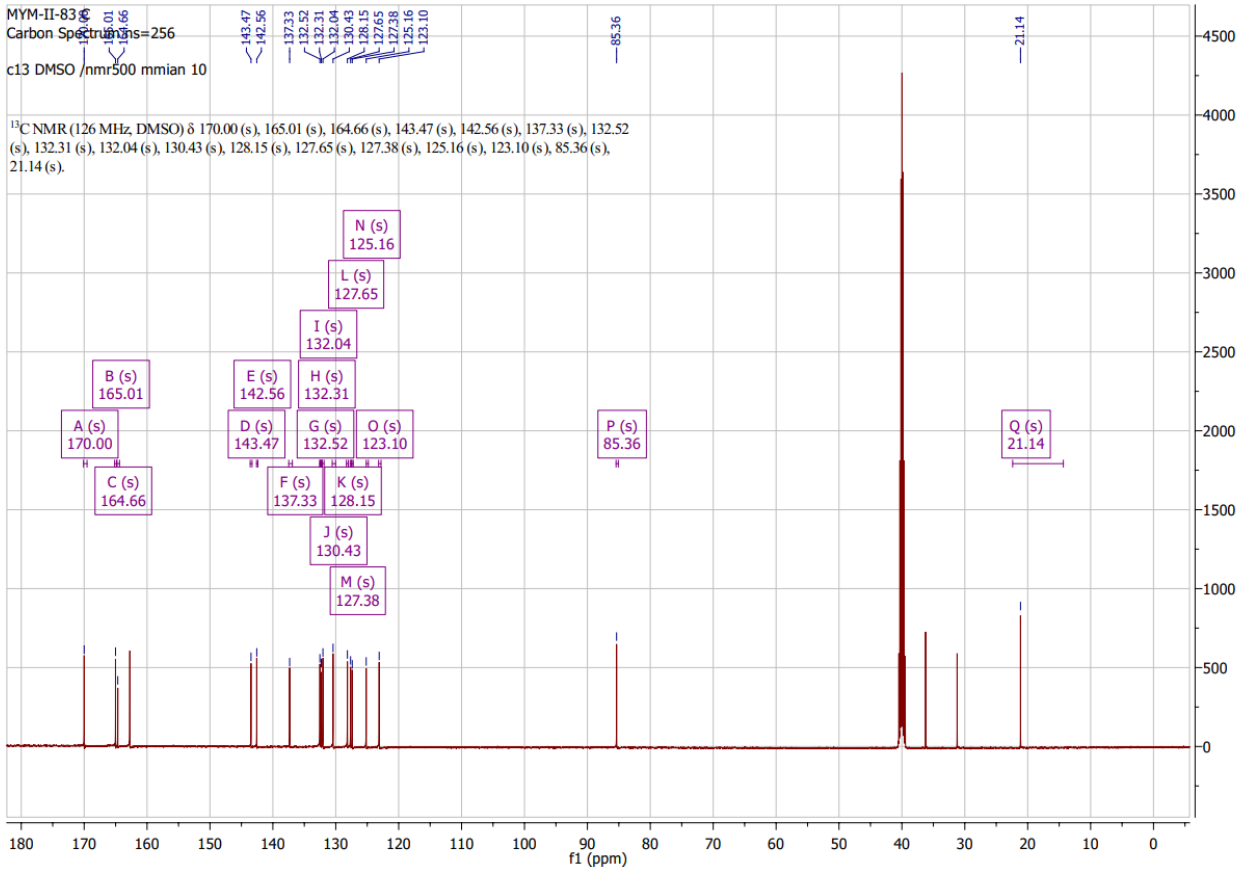

### HNMR of MYM-III-10 (10)

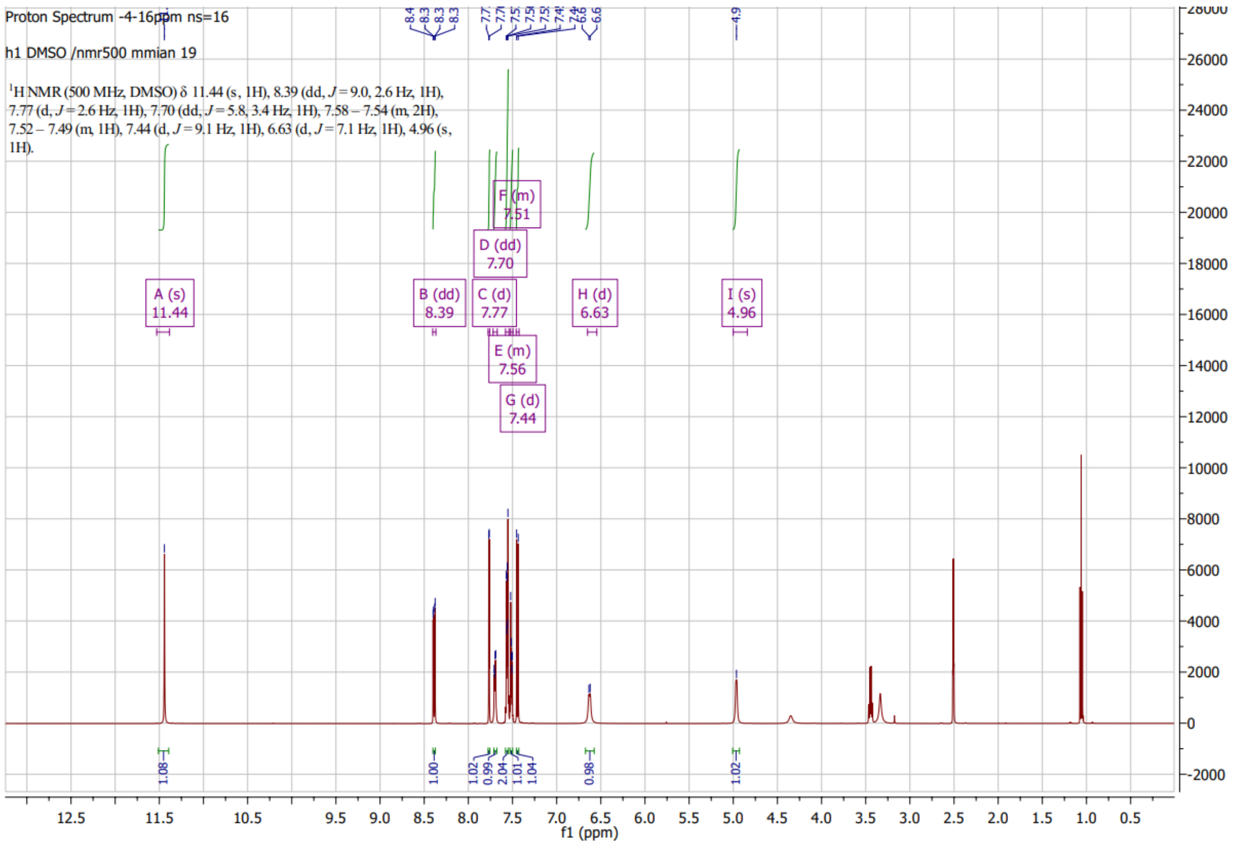

#### CNMR of MYM-III-10(10)

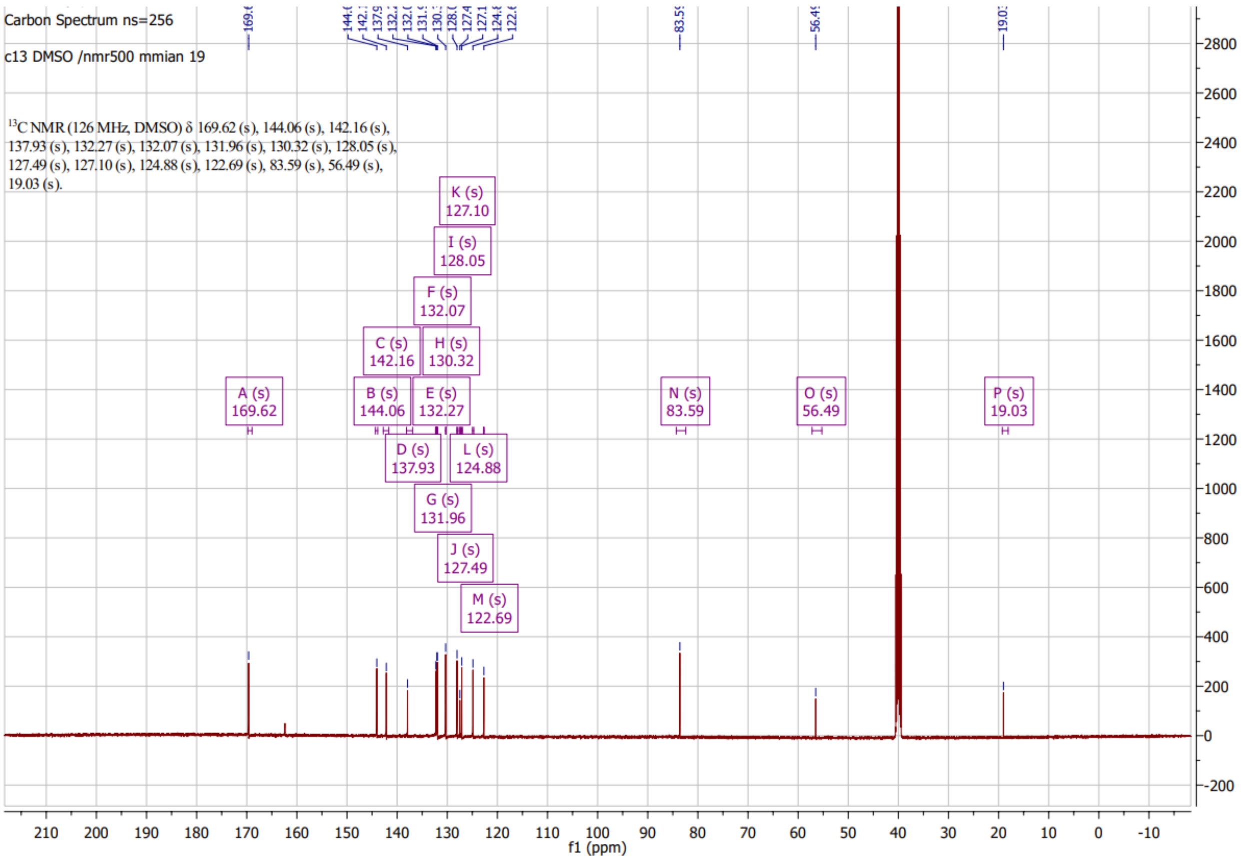

### HNMR of MYM-V-58 (11)

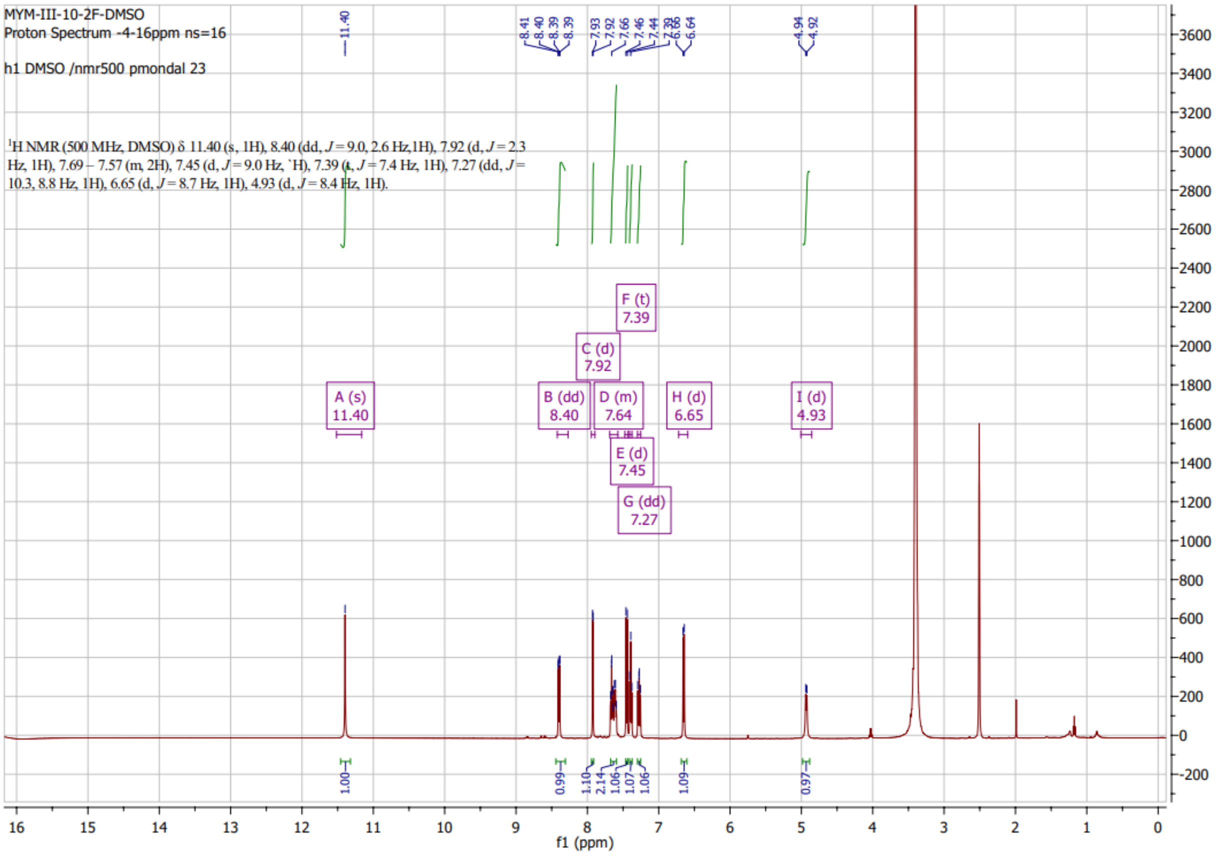

### HNMR of MYM-V-50 (13)

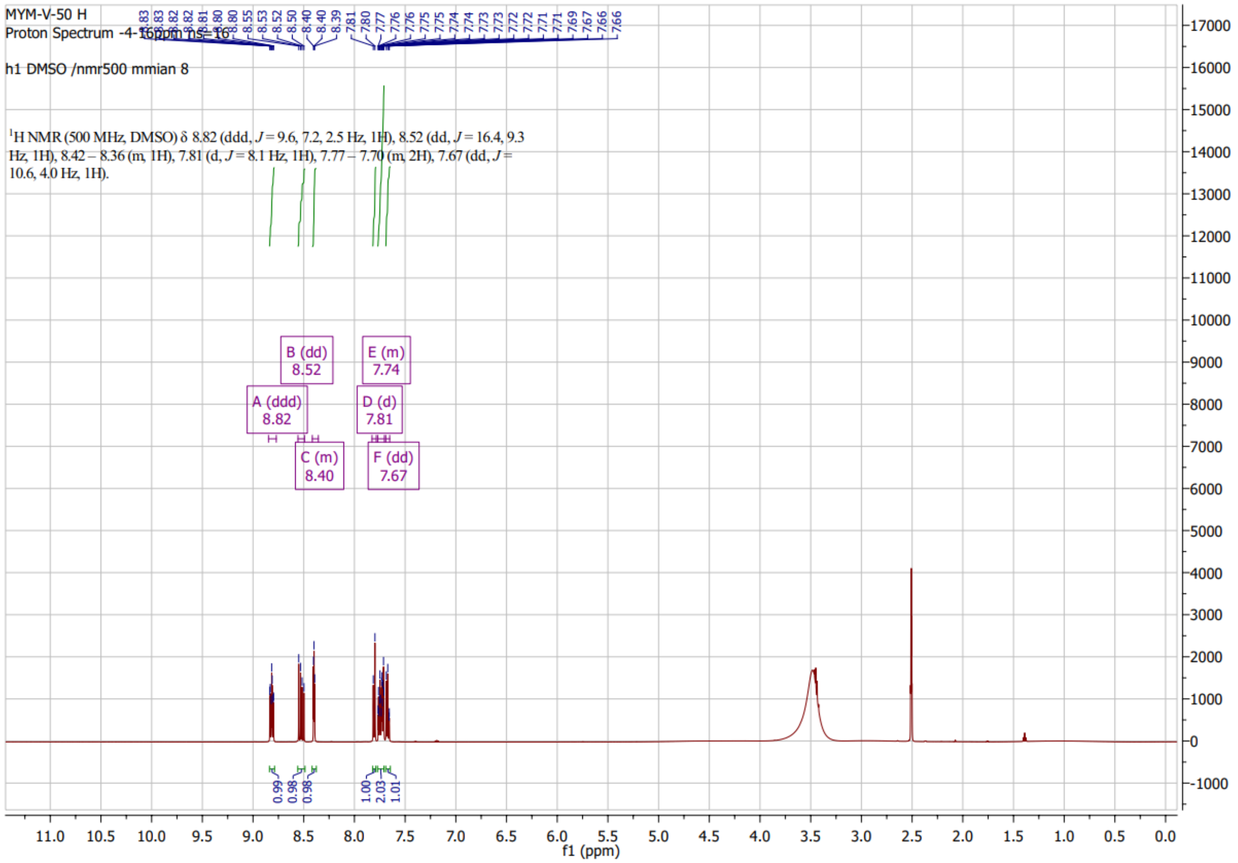

#### CNMR of MYM-V-50 (13)

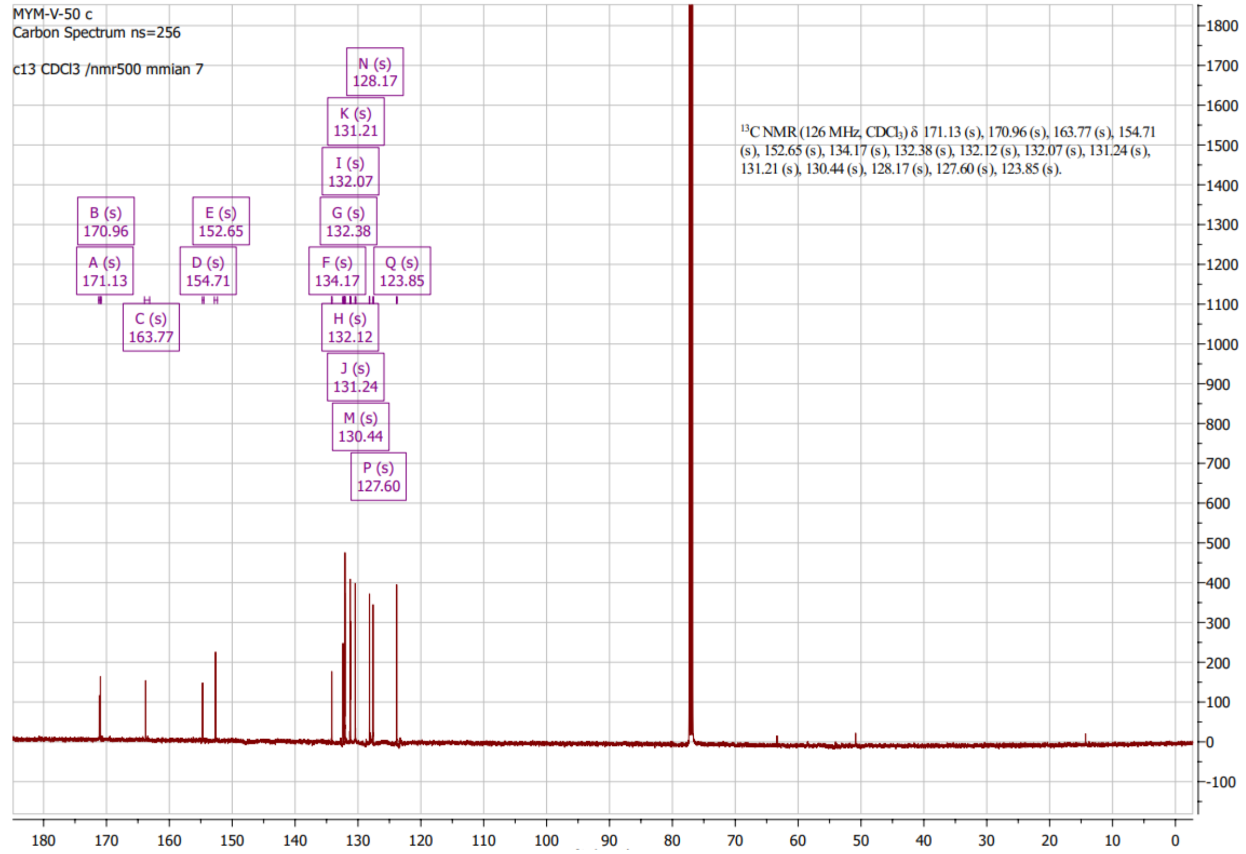

### HNMR of MYM-V-48 (15)

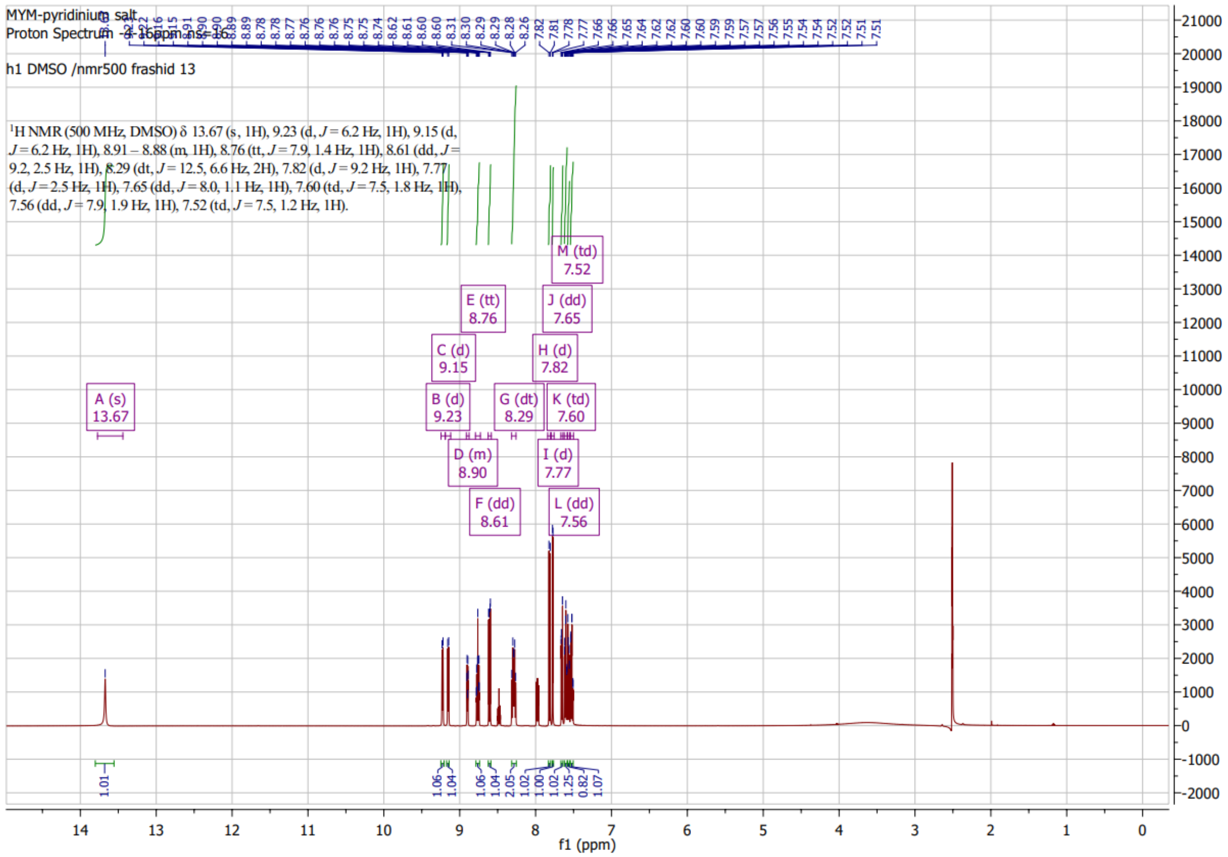

#### CNMR of MYM-V-48 (15)

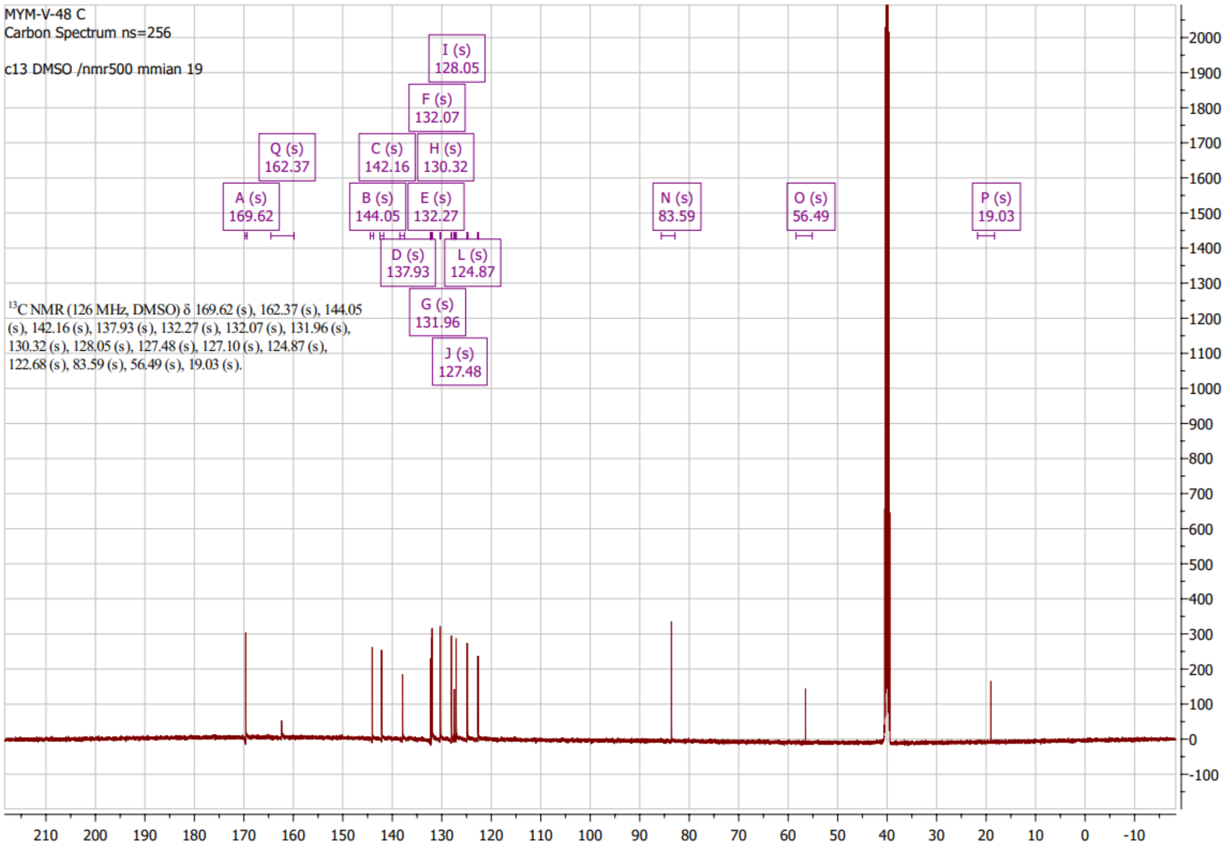

### HNMR of MYM-V-49 (16)

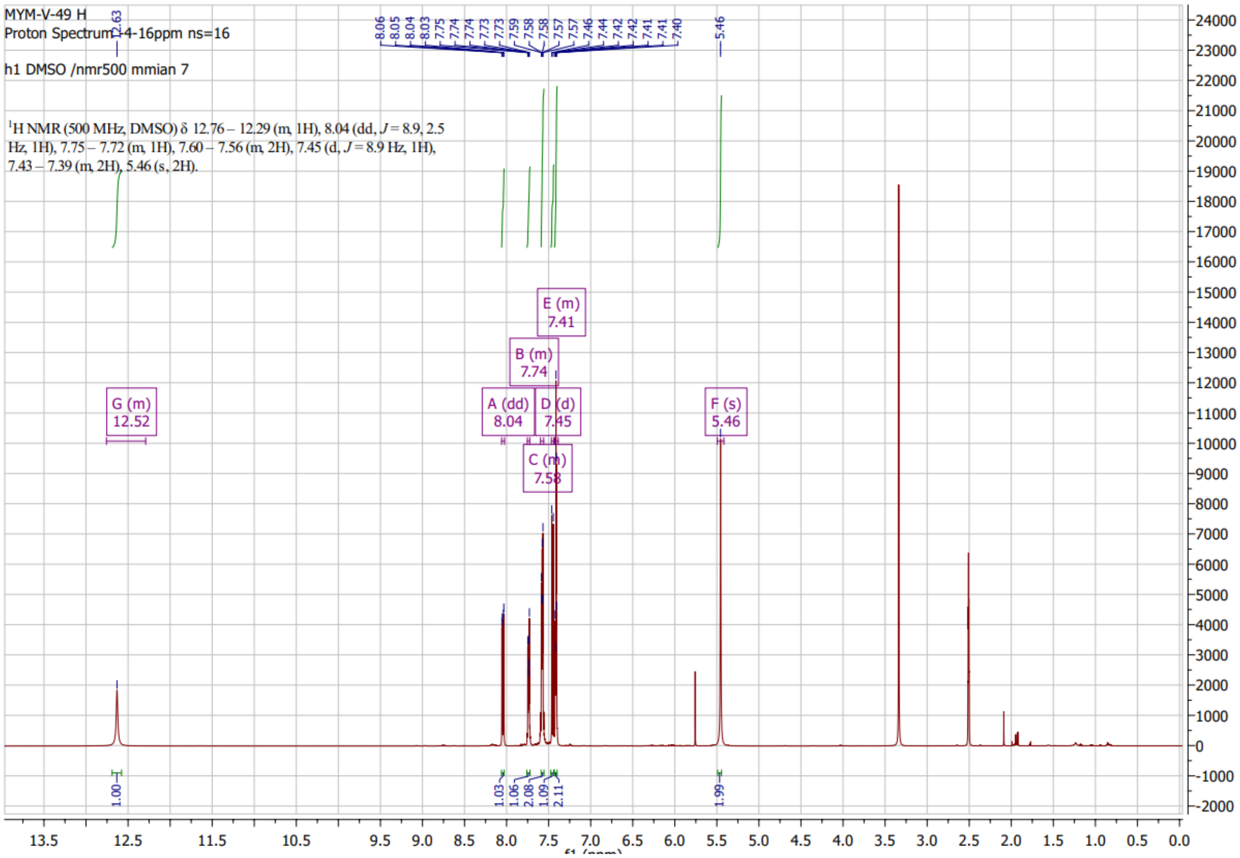

### CNMR of MYM-V-49 (16)

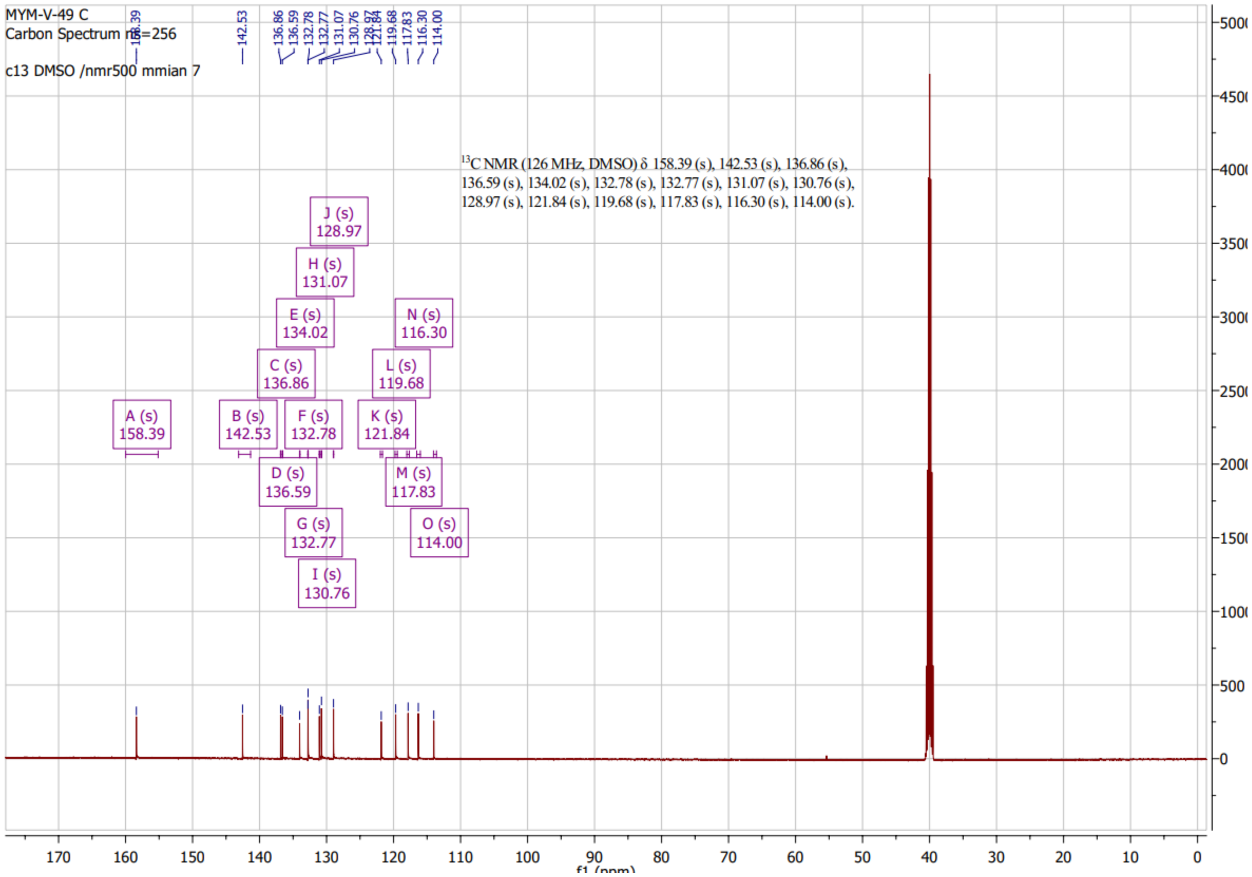

#### HNMR of MYM-V-56 (12)

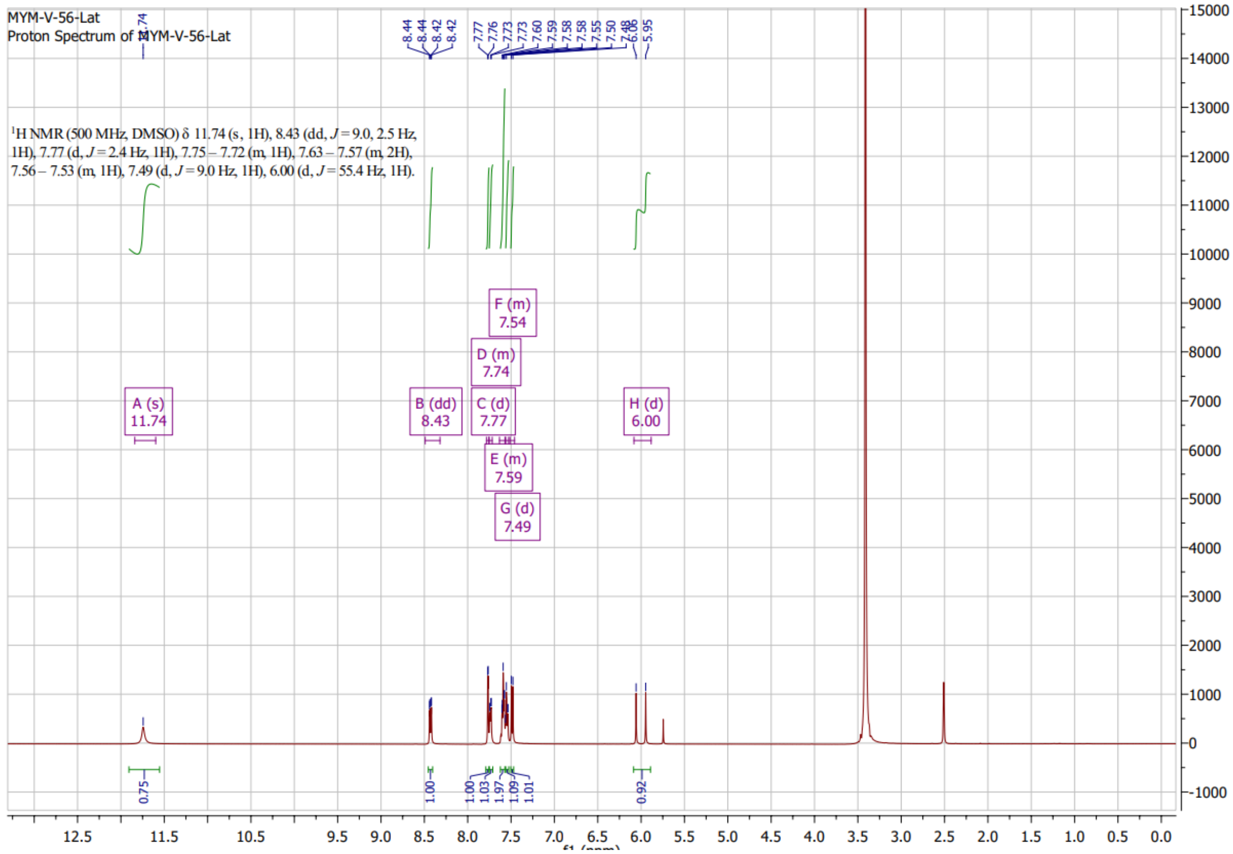

#### HNMR of MYM-II-72 (32)

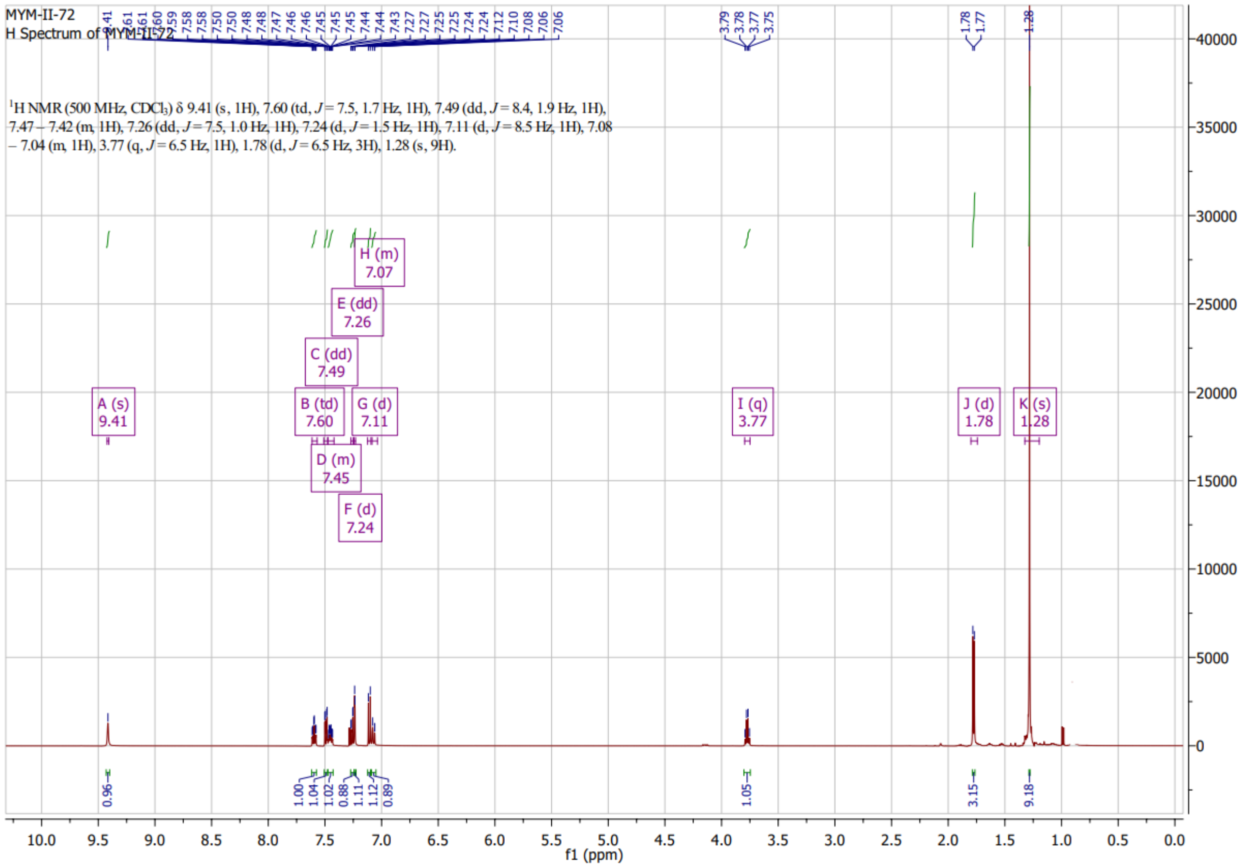

### CNMR of MYM-II-72 (32)

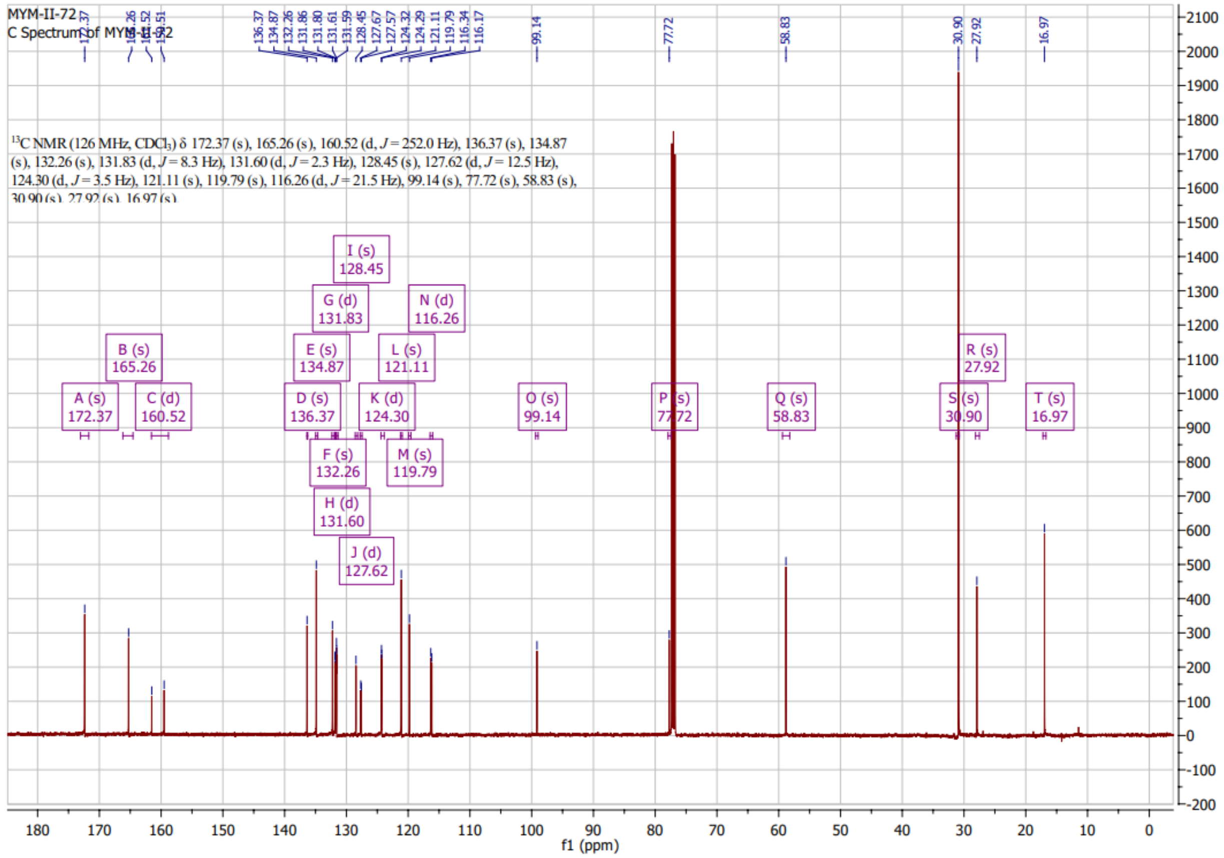

**HNMR of MYM-V-19 (24)**

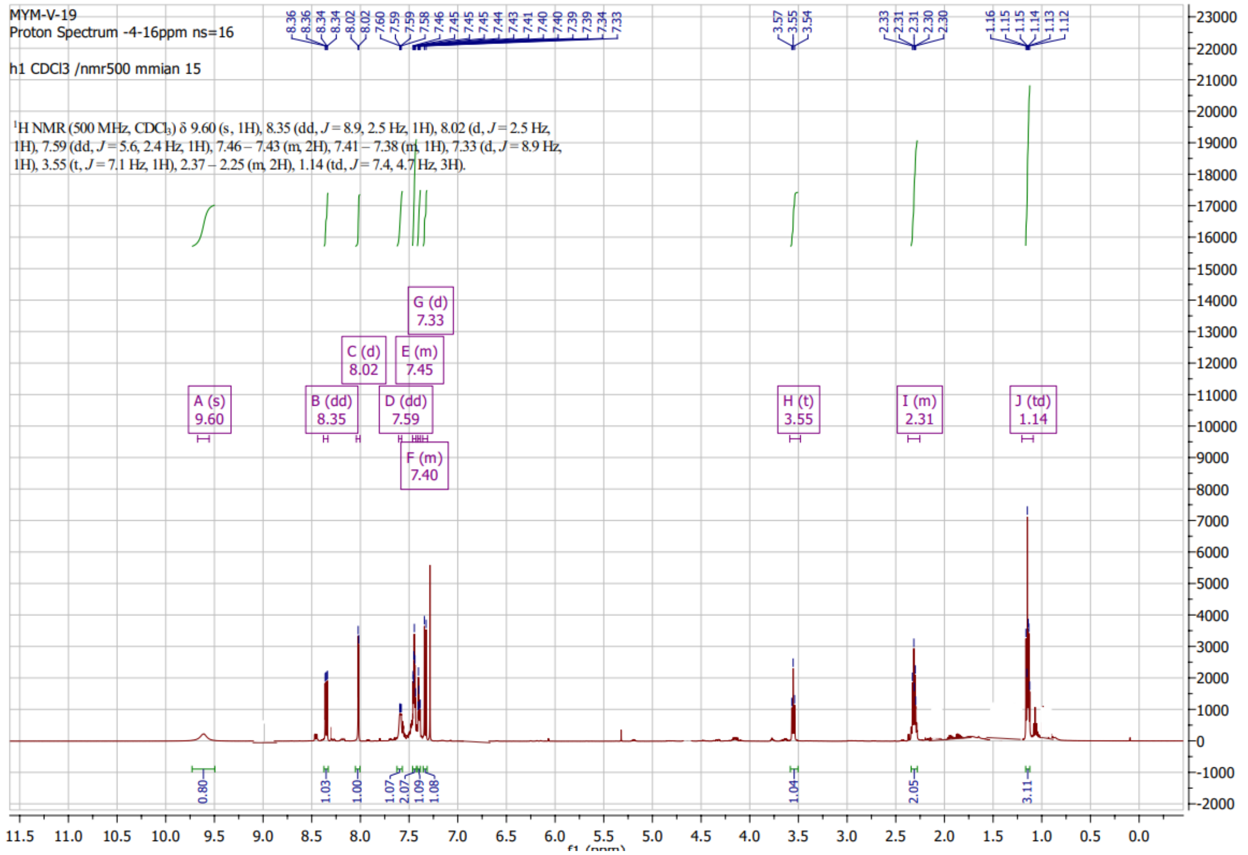

**CNMR of MYM-V-19 (24)**

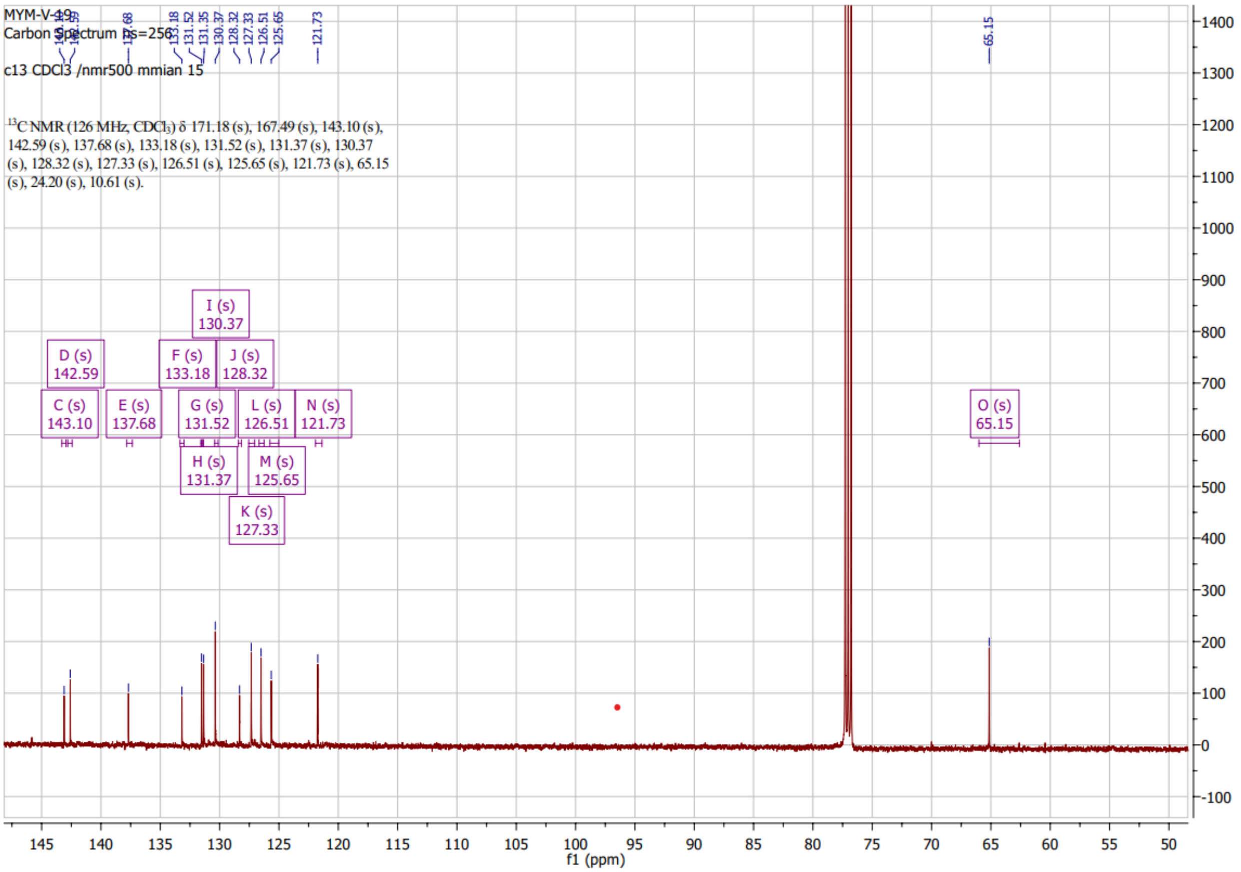

### HNMR of MYM-IV-12 (28)

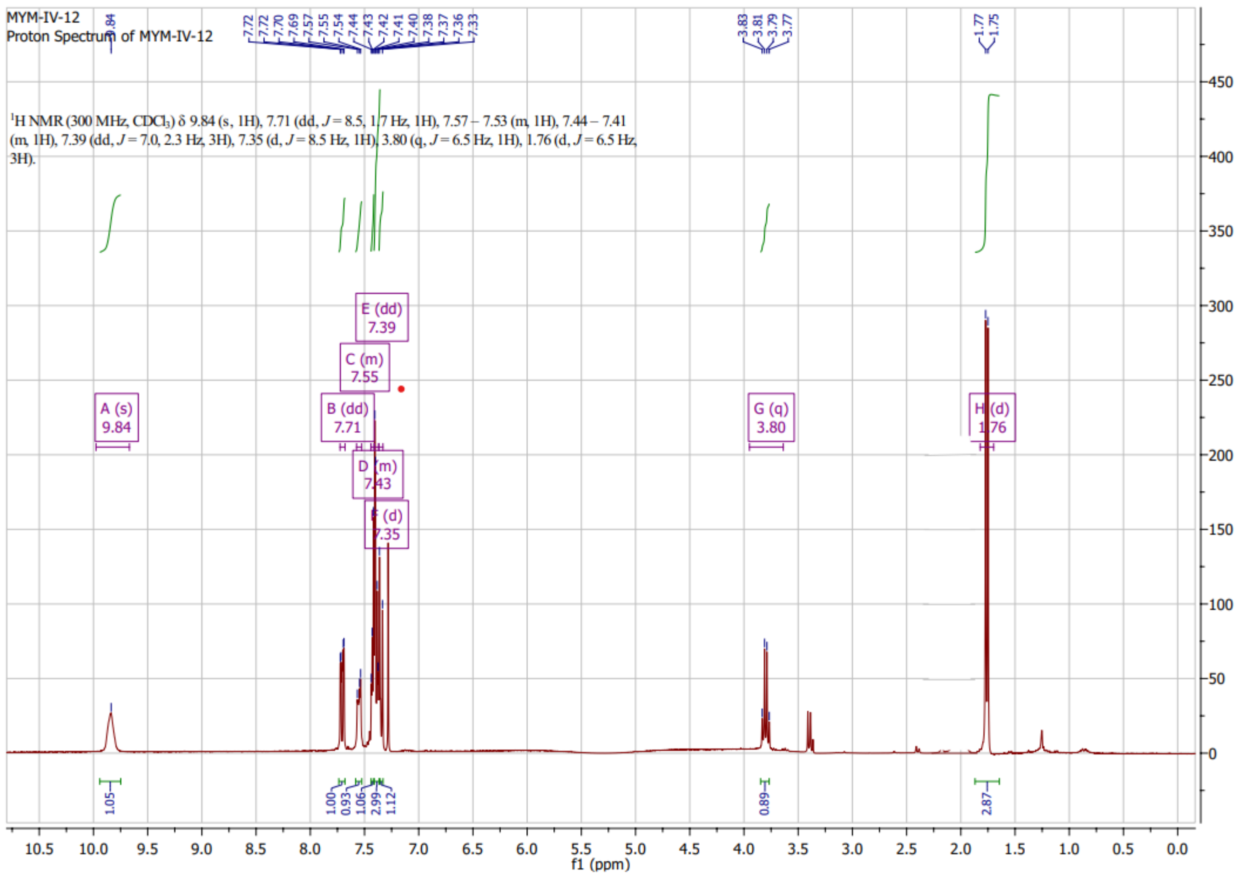

#### HNMR of MYM-V-51 (14)

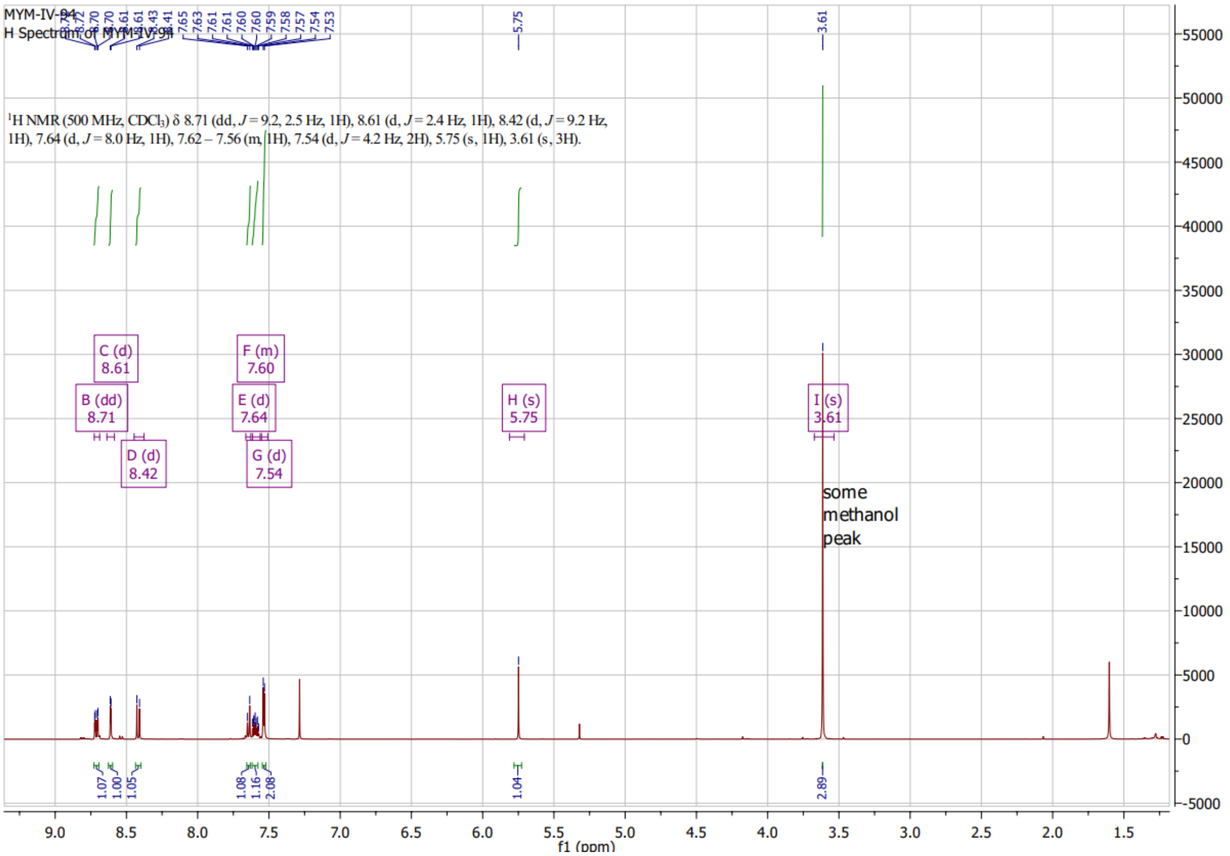

**CNMR of MYM-V-51 (14)**

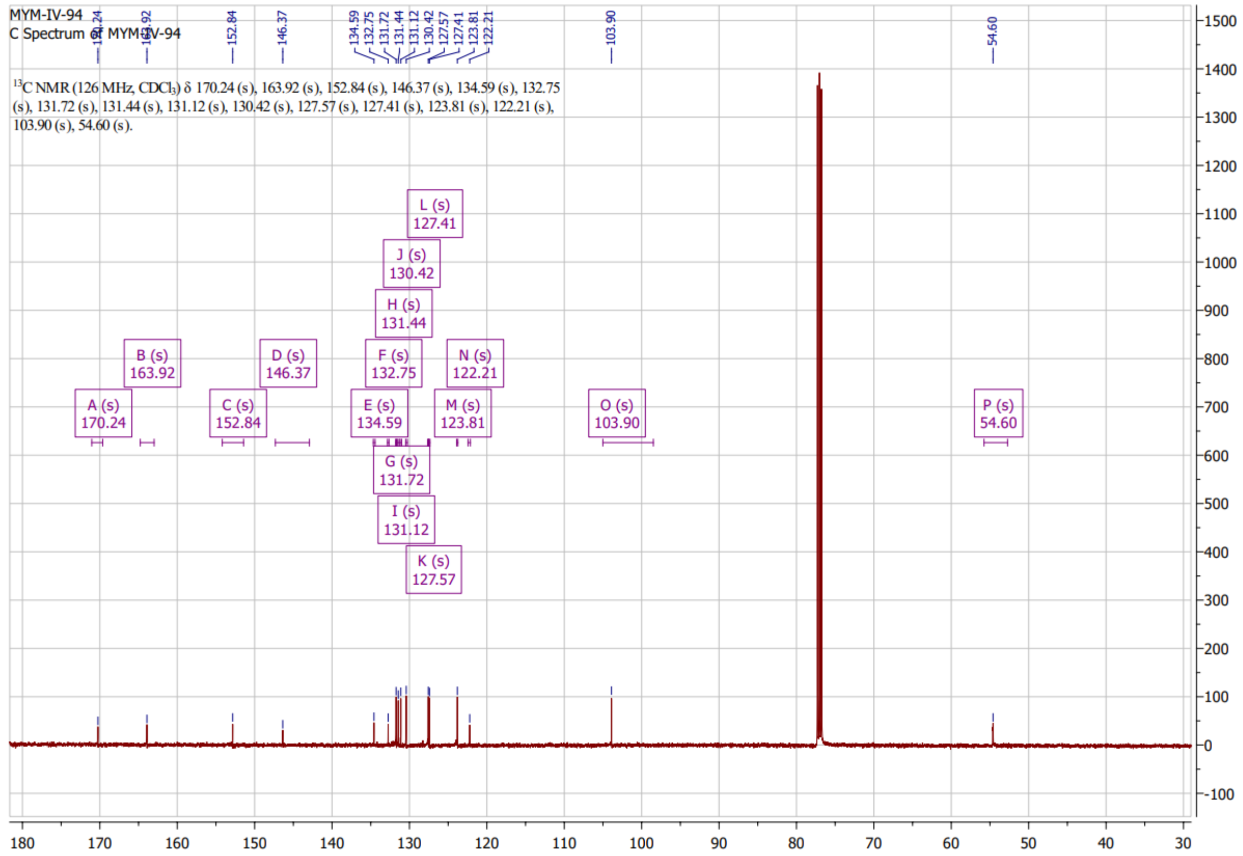

### HNMR of MYM-IV-02 (29)

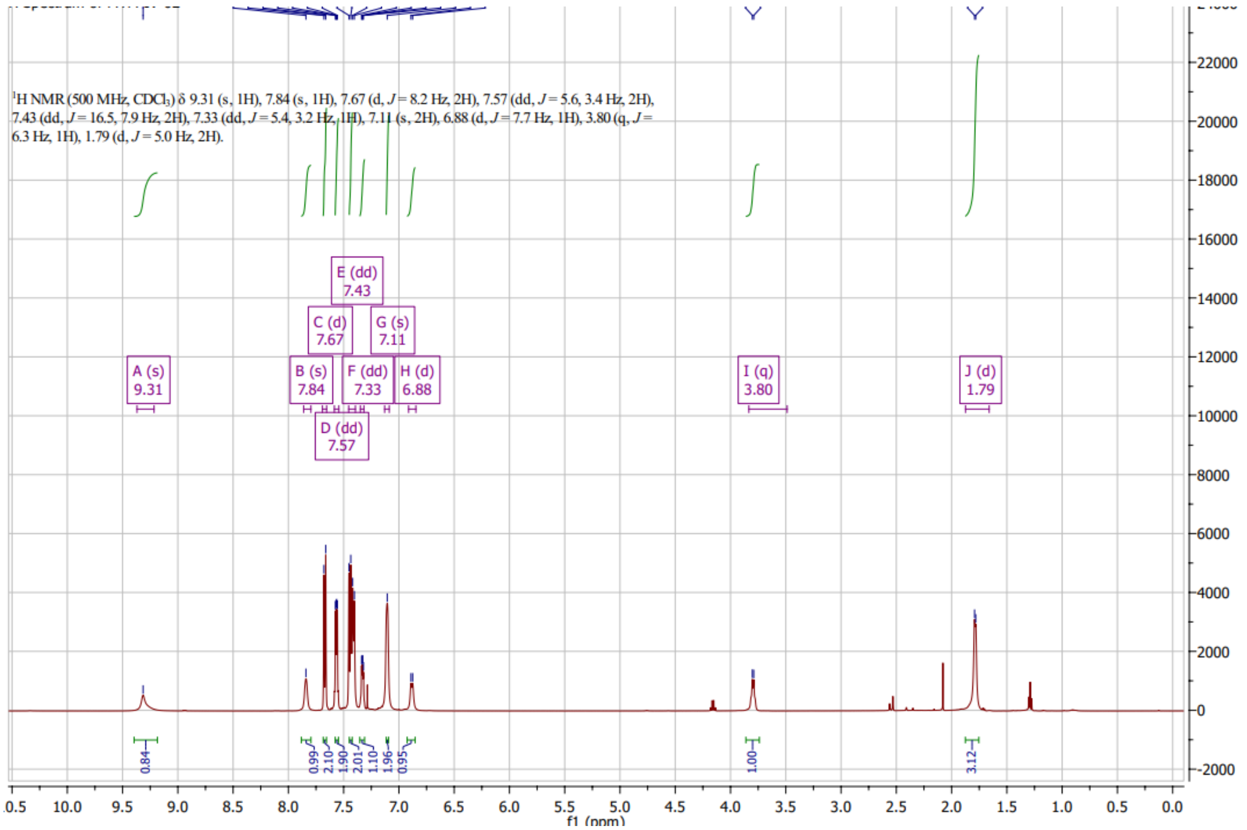

#### CNMR of MYM-IV-02 (29)

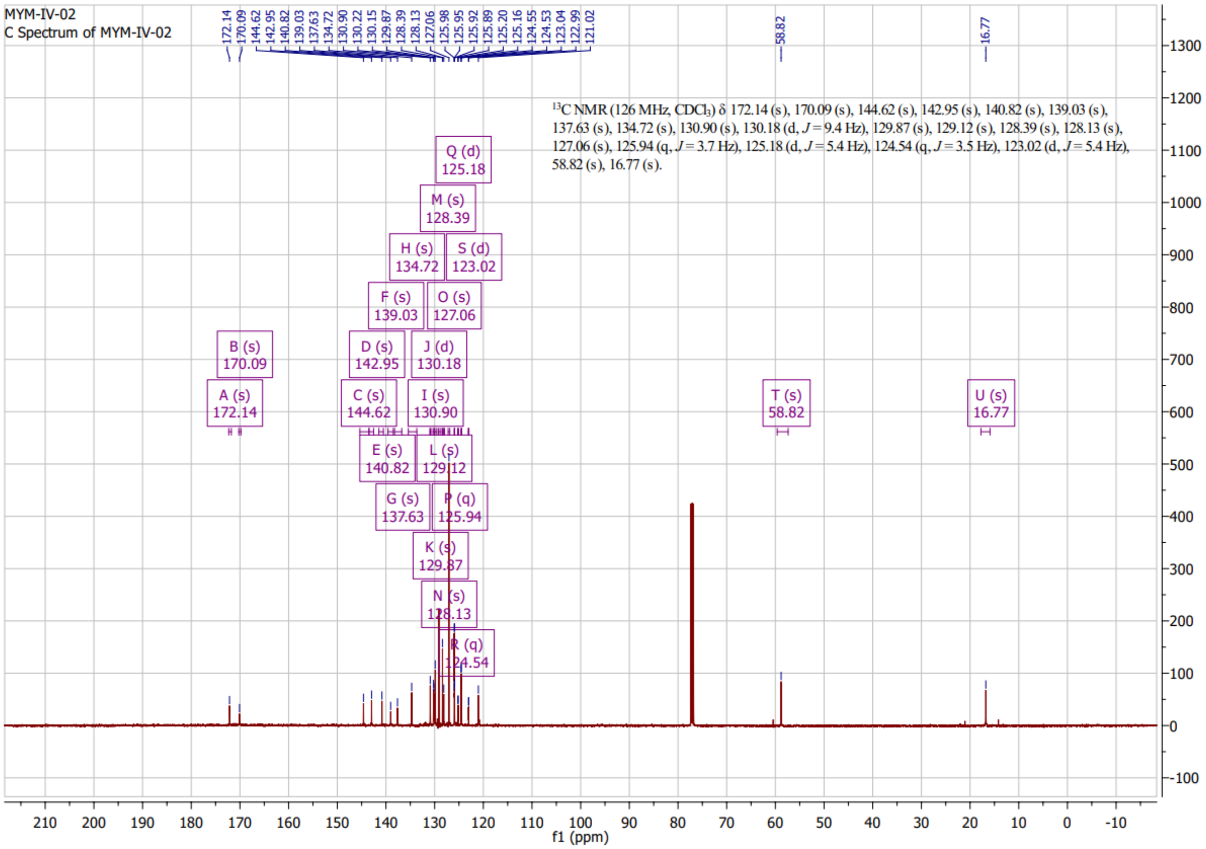

### HNMR of MYM-IV-01 (31)

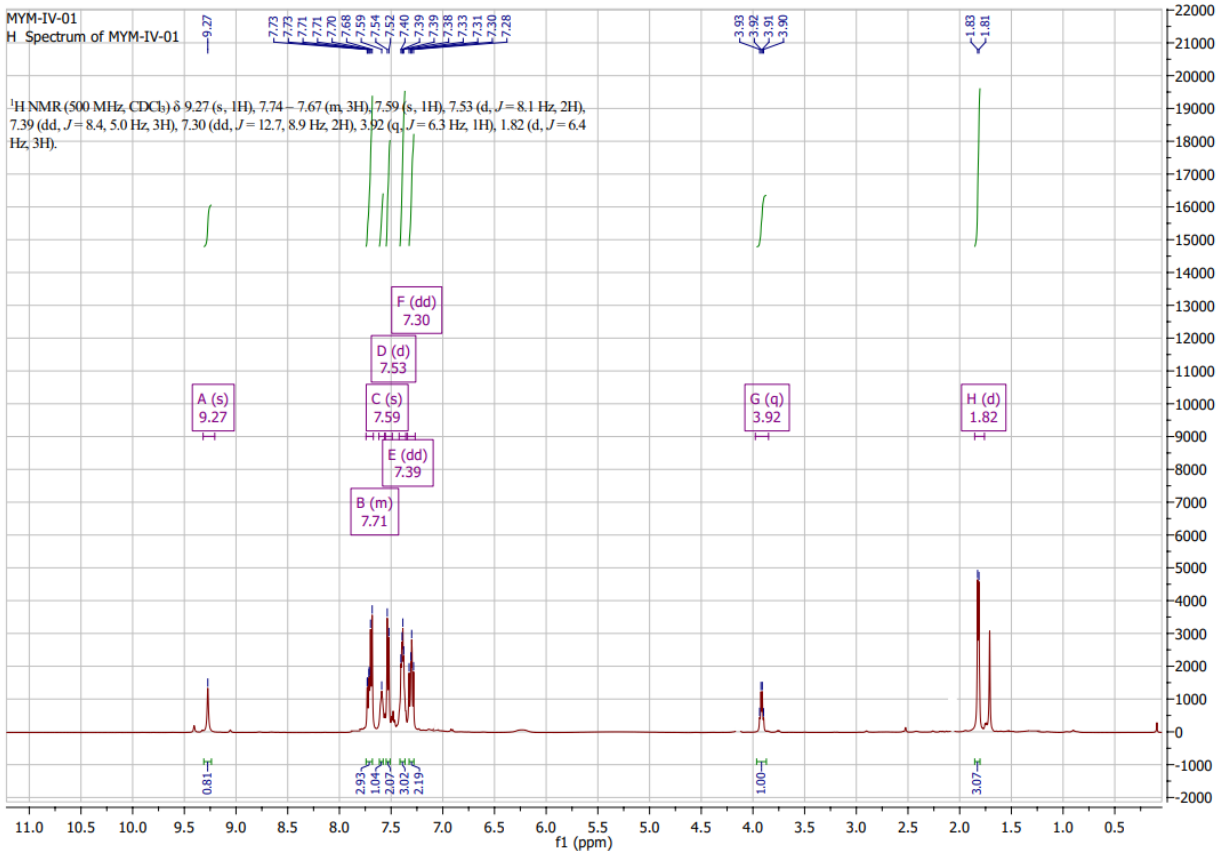

#### HNMR of MYM-IV-03 (30)

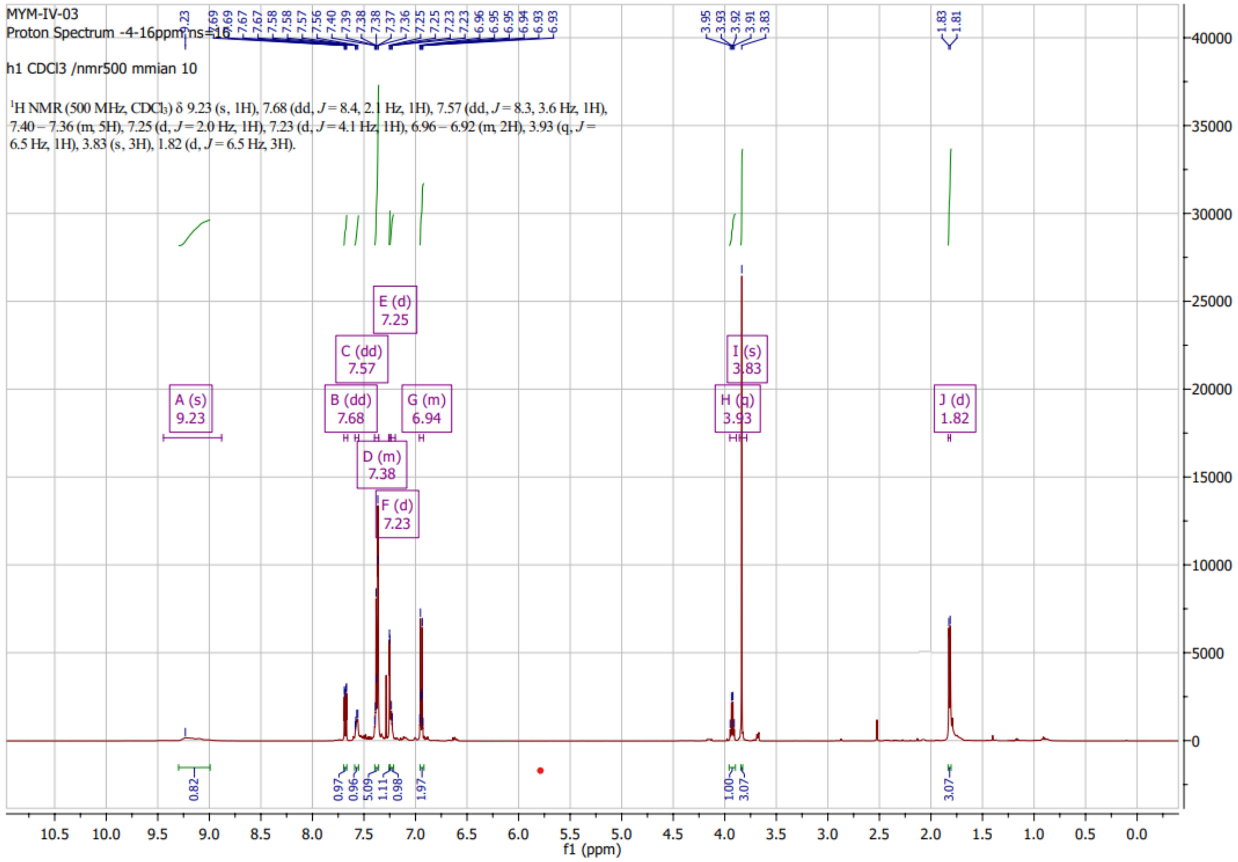

#### CNMR of MYM-IV-03 (30)

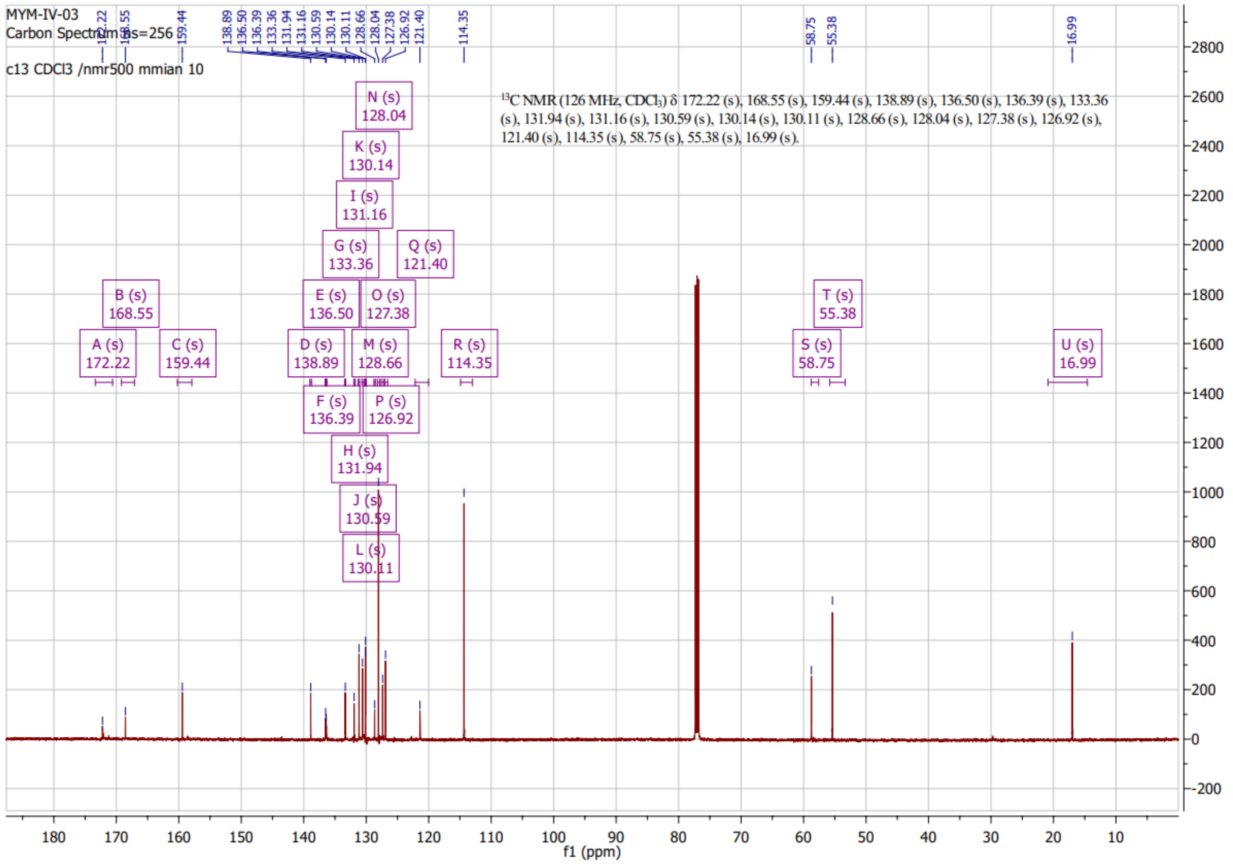

#### HNMR of MYM-III-98 (25):

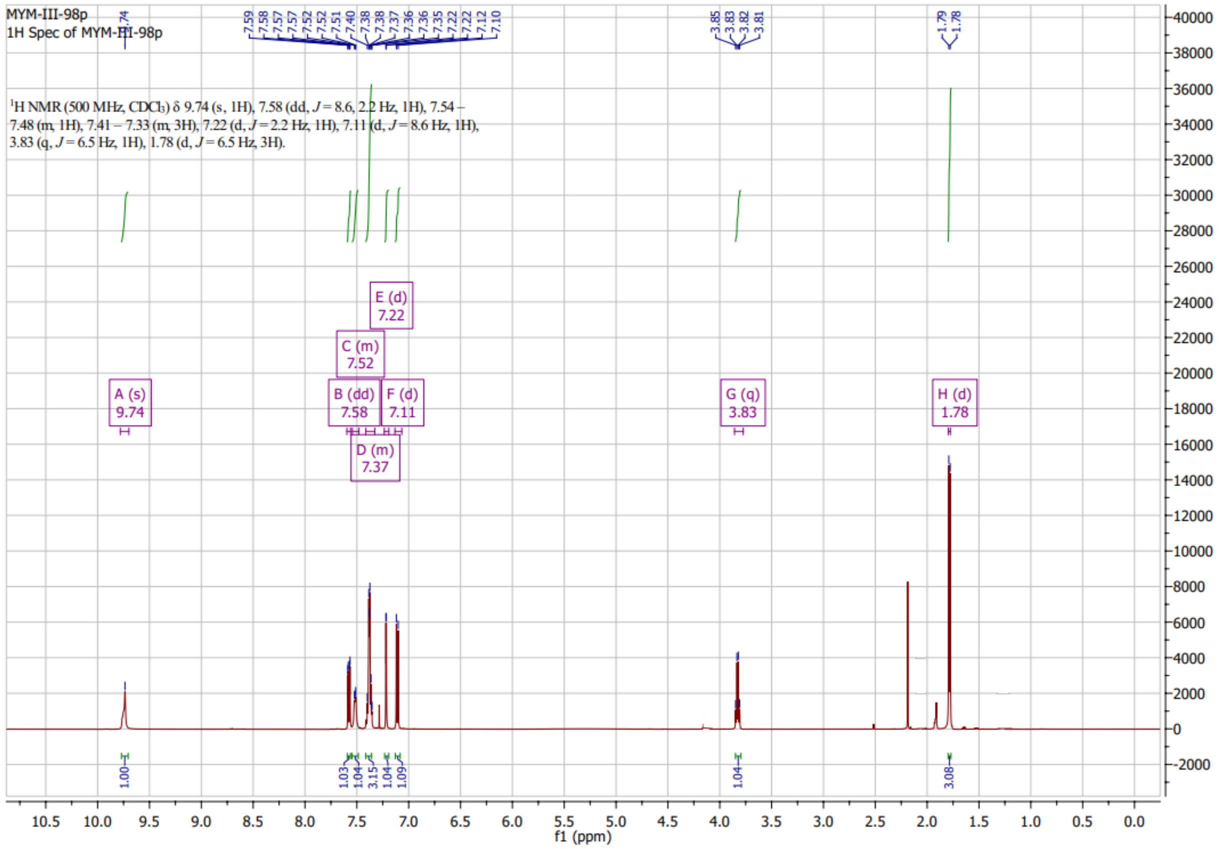

**CNMR of MYM-III-98 (25):**

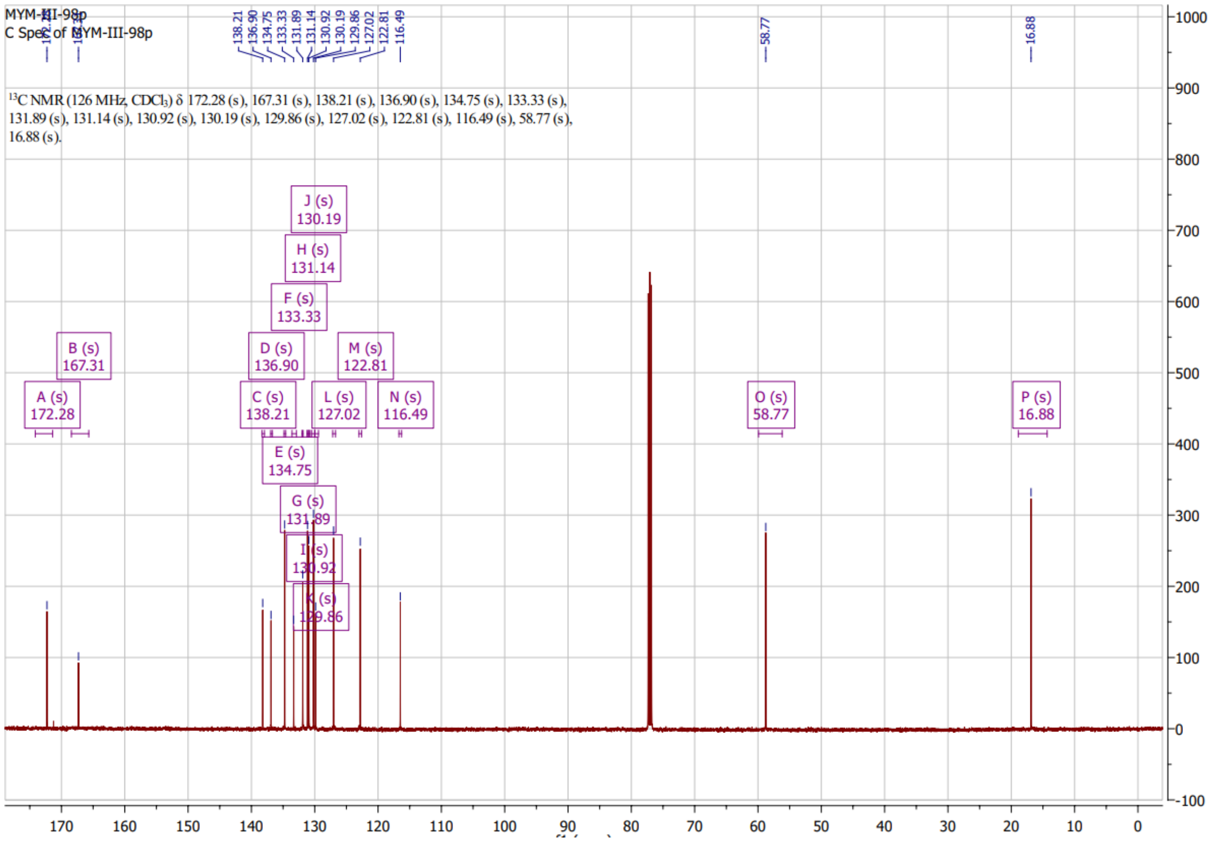



1. Huy C.P.; Pautet, G.V.; Hua, H. Separation of oxazepam, lorazepam, and temazepam enantiomers by HPLC on a derivatized cyclodextrin-bonded phase: Application to the determination of oxazepam in plasma. *Journal of Biochemical and Biophysical Methods* ***2002*** 54(1-3):287-99. DOI: 10.1016/S0165-022X(02)00123-9
